# First-in-Class Small Molecules Inhibit c-MET Activity Through Self-Regulatory Elements in Its Extracellular Domain

**DOI:** 10.64898/2026.05.01.722033

**Authors:** Okan Sezgin, Yeliz Yılmaz, Ezgi Bağırsakçı, Aykut Üren, Neşe Atabey, Serdar Durdağı

**Affiliations:** Laboratory for Innovative Drugs (Lab4IND), Computational Drug Design Center (HİTMER), Bahçeşehir University, 34734, İstanbul, Türkiye; Computational Biology and Molecular Simulations Laboratory, Department of Biophysics, School of Medicine, Bahçeşehir University, 34734, İstanbul, Türkiye; Izmir Biomedicine and Genome Center, Izmir, 35340, Türkiye; Department of Oncology, Georgetown University, Washington, DC., USA; Galen Research Center,Izmir Tinaztepe University, Buca, Izmir, Türkiye; Department of Medical Biology and Genetics, Faculty of Medicine, Izmir Tinaztepe University, 35340, Buca, Izmir, Türkiye; Molecular Therapy Laboratory, Department of Pharmaceutical Chemistry, School of Pharmacy, Bahçeşehir University, 34353, İstanbul, Türkiye; Quantitative System Biology Lab, Faculty of Medicine, Biruni University, İstanbul, Türkiye

## Abstract

Aberrant HGF–c-MET signaling is a major driver of hepatocellular carcinoma (HCC) progression and a clinically validated therapeutic axis, but current inhibitors predominantly target the intracellular kinase domain and remain vulnerable due to limited selectivity and resistance development. We therefore pursued an upstream strategy based on small molecules that target the extracellular HGF–c-MET interaction interface. We combined large-scale virtual screening of more than one million compounds from the ChemDiv and Enamine libraries with molecular dynamics (MD) simulations, steered MD, MM/GBSA profiling, and iterative lead optimization to identify candidate c-MET inhibitors targeting its extracellular (EC) domain. In HGF-stimulated HuH7 cells, selected compounds suppressed c-MET autophosphorylation, reduced cell viability, and inhibited long-term colony formation. Surface plasmon resonance (SPR) further confirmed direct binding of L083-1287 and 8008-3424 to the recombinant c-MET ectodomain. Mechanistic analyses identified previously unrecognized hotspot residues on the c-MET EC domain and a novel inhibitory network spanning multiple c-MET ectodomain interfaces. L083-0077 displayed the most consistent interaction pattern within this framework, including stabilization of key hotspot residues and preserved binding under acidic conditions relevant to the tumor microenvironment. Zebrafish xenograft assays with selected early hit compounds revealed compound-dependent developmental liabilities supporting the use of this model as an early *in vivo* prioritization step during lead optimization. These findings establish EC interface-directed c-MET inhibition as a promising therapeutic strategy in HCC and provide a mechanism-guided platform for the development of selective, upstream c-MET inhibitors with the potential to complement or overcome limitations of kinase-directed therapies.

## 1. Introduction

The hepatocyte growth factor (HGF)–c-MET signaling axis is a fundamental regulator of tissue homeostasis, embryonic development, and regenerative responses. Under physiological conditions, activation of the c-MET receptor by engagement of its ligand HGF coordinates diverse cellular programs, including proliferation, survival, motility, morphogenesis, angiogenesis, and wound healing.[1] These functions are essential for normal tissue repair and organ maintenance; however, the same signaling circuitry can be highjacked in pathological settings. In cancer, persistent or deregulated activation of HGF–c-MET signaling promotes tumor cell proliferation, epithelial–mesenchymal transition (EMT), invasion, metastatic dissemination, and resistance to therapy.[2–5] Aberrant c-MET signaling has been reported across a broad spectrum of malignancies, including hepatocellular carcinoma (HCC), gastric cancer, glioblastoma, and non-small-cell lung cancer (NSCLC), where it is frequently associated with aggressive clinical behavior and unfavorable prognosis.[6,7] These observations have established c-MET as both a biologically central oncogenic node and a therapeutically attractive target.

The oncogenic significance of c-MET is particularly compelling in hepatocellular carcinoma. HCC develops within a complex microenvironment shaped by chronic inflammation, fibrosis, altered growth-factor signaling, and progressive tissue remodeling, conditions in which HGF–c-MET signaling can become highly consequential.[6,7] In this context, c-MET contributes not only to tumor cell-intrinsic phenotypes such as proliferation and survival, but also to invasive behavior, vascular remodeling, and communication with stromal compartments.[1,6,7] Due to this broad biological effect, the pathway has remained a major focus in targeted drug discovery for more than two decades. Yet, despite sustained effort, durable pharmacological suppression of c-MET signaling remains challenging, particularly in tumors that adapt rapidly to selective pressure or that exploit compensatory growth-factor pathway activation.[6]

To date, the therapeutic landscape of c-MET inhibition has been dominated by small molecules directed against the intracellular tyrosine kinase domain. Several ATP-competitive inhibitors, including crizotinib, cabozantinib, and tepotinib, have advanced into clinical use and have demonstrated that c-MET is a druggable cancer target in principle.[8–11] Nevertheless, kinase-directed inhibition has important conceptual and practical limitations. First, the ATP-binding cleft of c-MET shares extensive structural similarity with other receptor tyrosine kinases (RTKs), making selective inhibition inherently difficult and increasing the risk of off-target pharmacology.[12] Second, as observed broadly across kinase-targeted therapies, treatment pressure can drive the emergence of resistance through gatekeeper mutations, amplification or rewiring of downstream effectors, and compensatory activation of parallel receptor systems. [13] Inhibitors directed at the extracellular domain (ECD) can mitigate mutation-driven resistance by sterically or allosterically hindering the ligand-induced dimerization required for downstream signaling. Third, ECD-targeted inhibitors that bind to the dimerization interface, physically preventing these disparate receptors from pairing and triggering downstream survival signals. Fourth, ECD inhibitors preclude receptor cross-talk by destabilizing the physical scaffolds and membrane hubs required for heteromeric signaling, rather than simply inhibiting kinase function. Together, these considerations suggest that alternative strategies capable of suppressing c-MET signaling earlier and more selectively may offer important therapeutic advantages.

An attractive but relatively underexplored alternative is to intervene at the extracellular (EC) step where HGF binds c-MET and initiates receptor activation. This interaction constitutes the obligate upstream event that precedes receptor dimerization, conformational rearrangement, and subsequent kinase-domain autophosphorylation. [12–14] As such, the EC HGF–c-MET interface offers a mechanistically distinct point of control. Blocking ligand–receptor engagement has the potential to prevent c-MET signaling at its origin, before activation propagates through intracellular kinase networks. This mode of intervention may also be less vulnerable to some classes of kinase-domain resistance mechanisms because it acts upstream of catalytic activation. In addition, the EC recognition surface of c-MET is structurally and chemically distinct from the highly conserved ATP-binding pockets shared among RTKs, providing the additional benefit of improved selectivity. [10–14] From a system biology perspective, EC blockade may also attenuate paracrine communication between tumor cells and their microenvironment, thereby influencing angiogenesis, stromal remodeling, and immune evasion in ways that kinase-directed inhibitors may not fully capture.[10–14]

Despite this conceptual appeal, most attempts to inhibit HGF–c-MET signaling at the EC level have relied on biologic modalities, such as neutralizing antibodies against HGF or c-MET and engineered protein-based antagonists.[14] These approaches have provided important proof of concept and, in some cases, have advanced into clinical evaluation.[14] However, biologics face well-recognized translational constraints. Their relatively large size can limit penetration into solid tumors and reduce access to spatially restricted receptor populations. They may also present formulation, immunogenicity, and exposure-related challenges that complicate broad therapeutic use.[15] Small molecules, in contrast, offer advantages in oral bioavailability, tissue distribution, synthetic tunability, and iterative optimization. For many areas of targeted therapy, they remain the most versatile and clinically scalable modality. The difficulty, however, is that EC receptor–ligand interfaces are often mediated by large, shallow, and conformationally dynamic protein–protein interaction (PPI) surfaces rather than by deep, well-enclosed small-molecule pockets.[16] The HGF–c-MET interface exemplifies this challenge, involving extended contact regions and a dynamic structural context associated with receptor activation. For this reason, the EC interface has been viewed as a difficult target for conventional small-molecule discovery.

Recent advances in structural biology and computational chemistry now provide an opportunity to revisit this assumption. High-resolution cryo-EM and crystallographic studies have substantially improved the structural definition of the HGF–c-MET EC complex and have begun to delineate the residues and interaction hotspots that govern ligand recognition and receptor activation. [12] In parallel, modern computational workflows, including large-scale virtual screening, molecular dynamics (MD) simulations, steered MD, and free-energy-based prioritization, have made it increasingly feasible to interrogate broad chemical spaces against dynamic and non-classical binding surfaces.[17,18] These approaches are especially valuable for targets in which binding competence depends not only on static structural complementarity but also on conformational adaptability, persistence of contact networks, and coupling to protein motions. In this sense, EC receptor interfaces once considered poorly suited for small-molecule intervention are becoming progressively more accessible to mechanism-informed discovery strategies. [12,17,18]

Within this emerging framework, c-MET represents a particularly compelling system because its EC domain is not simply a passive ligand-binding surface but a structurally organized regulatory unit whose conformational behavior is closely linked to receptor activation. [12]

This raises an important possibility: small molecules that engage carefully selected EC hotspots may not merely block HGF binding sterically, but may also reshape the conformational landscape of the receptor in ways that disfavor productive activation. If so, an effective EC inhibitor would be expected to combine at least three features: stable engagement of interface-relevant residues, compatibility with the dynamic architecture of the c-MET ectodomain, and measurable downstream functional consequences in biologically relevant cancer models. Developing such compounds therefore requires more than conventional docking alone. It requires an integrated framework that connects structural recognition, energetic stability, conformational dynamics, and cellular function.

In the present study, we developed such an integrative discovery strategy to identify and characterize small molecules that disrupt the HGF–c-MET EC interaction interface. By combining large-scale virtual screening of more than one million compounds with MD simulations, steered MD, and free-energy profiling, we prioritized candidates predicted to engage critical EC hotspot residues. [17,18] We then evaluated the resulting compounds experimentally in hepatocellular carcinoma cells, assessed their effects on c-MET activation and cell proliferation and invasion capabilities. We then examined direct binding to the recombinant c-MET ectodomain. In parallel, iterative hit-to-lead and lead-optimization cycles were used to refine scaffold properties and to distinguish compounds with broader interface activity from those with greater c-MET selectivity. This workflow was designed not only to identify active hit compounds, but also to establish whether the EC c-MET interface can be productively and reproducibly modulated by small molecules. Scheme 1 provides a comprehensive overview of the workflow and methodologies employed throughout this study.

Beyond small molecule lead compound discovery, this study also addresses a broader mechanistic question: How EC ligand binding and inhibitory ligand engagement are coupled to conformational regulation of c-MET. By integrating simulation-derived motion analyses with experimental validation, we identify previously unrecognized hotspot regions within the EC and define a mechanistic framework linking inhibitor binding to altered receptor dynamics. In doing so, the work expands the chemical and conceptual space of c-MET inhibition beyond kinase-directed blockade and supports the EC HGF–c-MET interface as a tractable target for small-molecule intervention. More broadly, it provides a model for how dynamic receptor–ligand interfaces can be interrogated and modulated through mechanism-guided small-molecule discovery. [12,17,18]

**Scheme 1.**
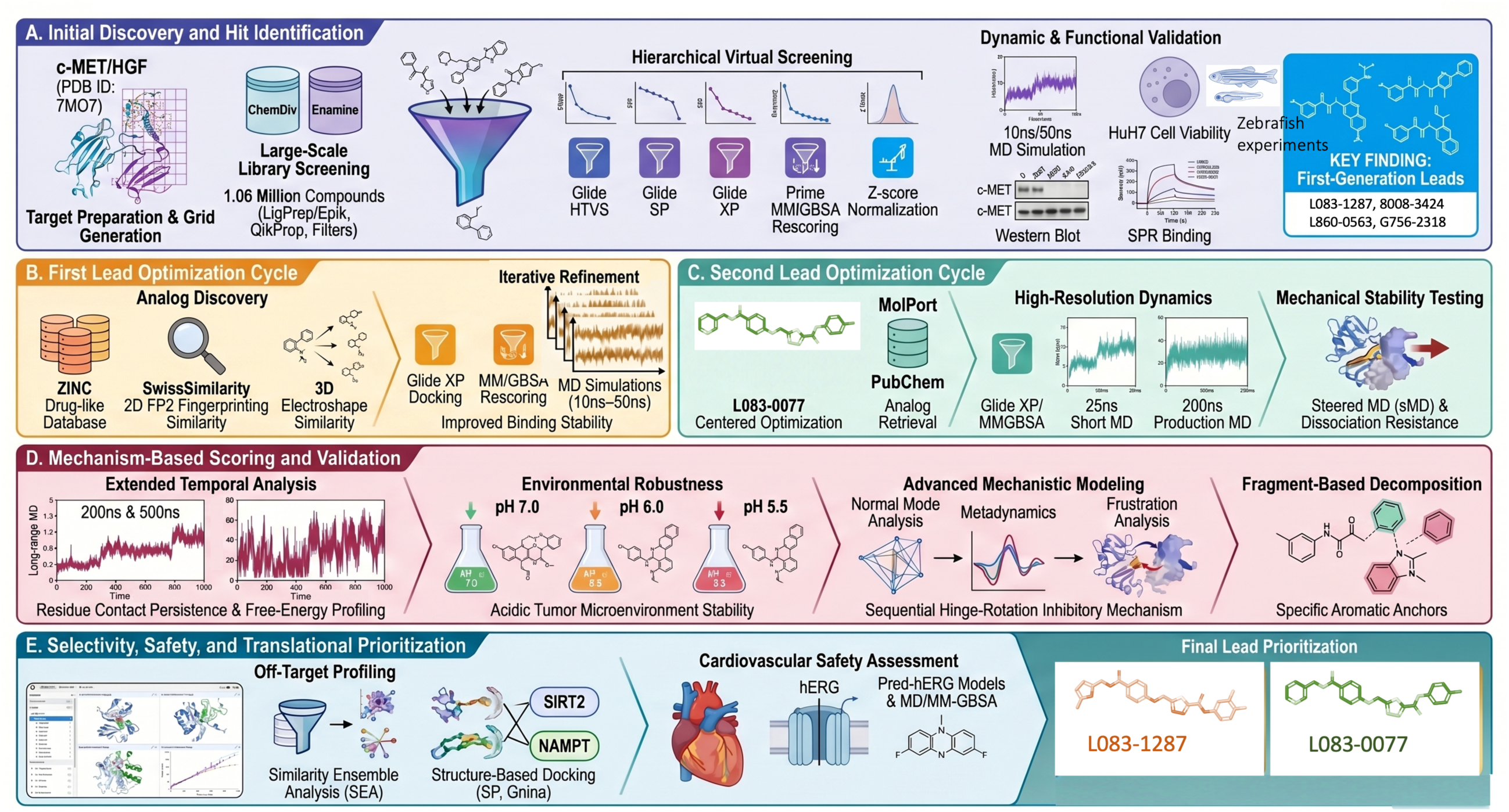
Summary of workflow and methodologies conducted throughout this study. (A) the steps involving the development of approaches tailored to the target structure, the execution of *in silico* analyses, and the experimental characterization of the identified hit compounds to evaluate their potential as lead compounds. (B) *In silico* and wet-lab experiments were performed to optimize the obtained lead compounds. (C) Optimizing pharmacophore characteristics observed in lead compounds during the first cycle using *in silico* methods to screen for new leads. (D) The stage of using mechanism-based scoring approaches in testing lead compounds. (E) Prior to their introduction into the literature, lead compounds underwent advanced technical research to investigate their off-target potential and binding efficiency.

## 2. Methods

### 2.1 Structural Preparation and Binding Site Definition

All structural modeling was conducted using the cryo-EM structure of the c-MET extracellular domain (ECD) (PDB ID: 7MO7) as the initial template. [12] The protein structure was prepared with the Schrödinger Protein Preparation Wizard by adding hydrogen atoms, assigning bond orders, and removing water molecules located more than 5 Å from heteroatom groups. Missing side chains and loop regions were modeled using Prime. [19] Disulfide bonds were manually assigned in accordance with reported post-translational modification data. [20] Protonation states were determined at physiological pH (7.4) using Epik [21] and further refined with PROPKA. [22] The hydrogen-bonding network was optimized, and the resulting structure was subjected to restrained energy minimization with the OPLS3e force field [23] until convergence reached an RMSD threshold of 0.30 Å. Binding-site residues were defined based on available structural and biochemical evidence from literature. [12,24] In particular, Lys47, Lys91, Phe112, and His114 from the HGF NK1 domain, together with Asn393 and Phe398 from the c-MET SEMA domain, were selected as interface hotspot residues. (Figure 1) These residues were used to define the centers of the binding-site grids. Three receptor grid systems were subsequently generated: one centered exclusively on HGF hotspot residues, one centered exclusively on c-MET hotspot residues, and one spanning the entire HGF–c-MET interface. Receptor grids were generated using the Receptor Grid Generation module [25] with OPLS3e-assigned charges, and side-chain flexibility was retained to account for local conformational variability.

**Figure 1.**
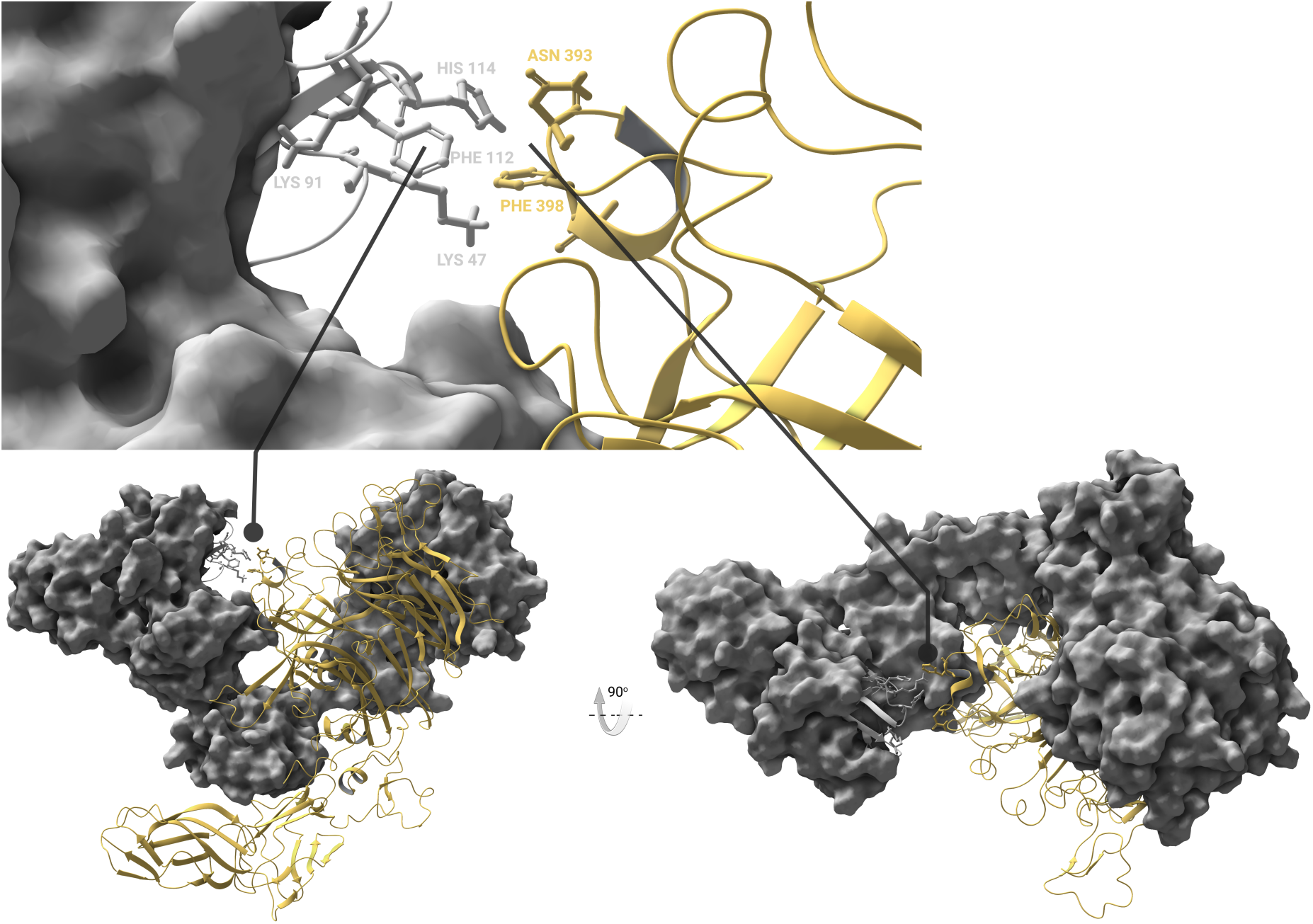
3D structures of the grids created from the region defined as the interface. The gold color-coded regions represent c-MET (Phe393, Asn398), while the dark-gray color-coded regions represent HGF (Lys47, Lys91, Phe112, His114).

### 2.2 Compound Libraries and Preprocessing

Two chemically diverse screening small molecule libraries were used in this study: the Enamine Hit Locator Library (460,160 compounds) [26] and the ChemDiv Representative Compounds Library (∼300,000 compounds). [27] Following preprocessing, including desalting, tautomer enumeration, and stereoisomer generation, the merged dataset comprised approximately 1.06 million structures. Bemis–Murcko clustering was applied to preserve scaffold diversity across the combined library. Compounds flagged by Pan-Assay Interference Compounds (PAINS) and Rapid Elimination of Swill (REOS) filters were excluded. [28] Ligands were prepared using Schrödinger LigPrep [29] and Epik [21] to generate relevant protonation states at pH 7.0 ± 2.0. All ligand structures were subsequently energy-minimized with the OPLS3e force field to obtain low-energy conformations prior to docking.

### 2.3 Virtual Screening Workflow

Virtual screening was conducted using a hierarchical workflow implemented in Schrödinger Maestro. Compounds were first filtered on the basis of QikProp [30] drug-likeness predictions to exclude molecules with unfavorable pharmacokinetic properties. The remaining set was further refined by applying Lipinski’s Rule of Five and removing compounds containing reactive functional groups. Docking was then performed sequentially with Glide using three levels of precision: High-Throughput Virtual Screening (HTVS) to rapidly eliminate the lowest-ranked 90% of compounds, Standard Precision (SP) docking for more detailed pose evaluation, and Extra Precision (XP) docking for final high-accuracy scoring. [25] Ligand flexibility was retained throughout the workflow, and selected side chains within the binding pocket were treated as rotatable to account for local conformational adaptability. Docking poses were subsequently rescored using Prime MM/GBSA [19,31] with the VSGB 2.0 implicit solvation model [32], and Z-score normalization was applied to standardize score distributions. (Figure S1) The highest-ranking compounds were then advanced for MD simulations.

### 2.4 MD Simulations

All-atom MD simulations were carried out using Desmond [33]. Each system was solvated in an explicit TIP3P water box under periodic boundary conditions and neutralized with Na^+^ and Cl^-^ ions. Simulations were performed in the NPT ensemble at 310 K and 1.01325 bar, using the Nosé–Hoover chain thermostat [34–36] for temperature control and the Martyna–Tobias–Klein barostat [37] for pressure regulation. Electrostatic interactions were calculated with a 9 Å cutoff, and the integration timestep was set to 2 fs, with long-range interactions updated every 6 fs. Short MD simulations were initially performed for 40 selected compounds, with each ligand simulated for 10 ns in five independent replicas initiated with different random seeds. Based on docking performance, binding free-energy estimates, and binary QSAR analysis, 31 ligands were subsequently advanced to extended 50 ns MD simulations, each performed in triplicate. (Table 1, Table S1).

**Table 1.**
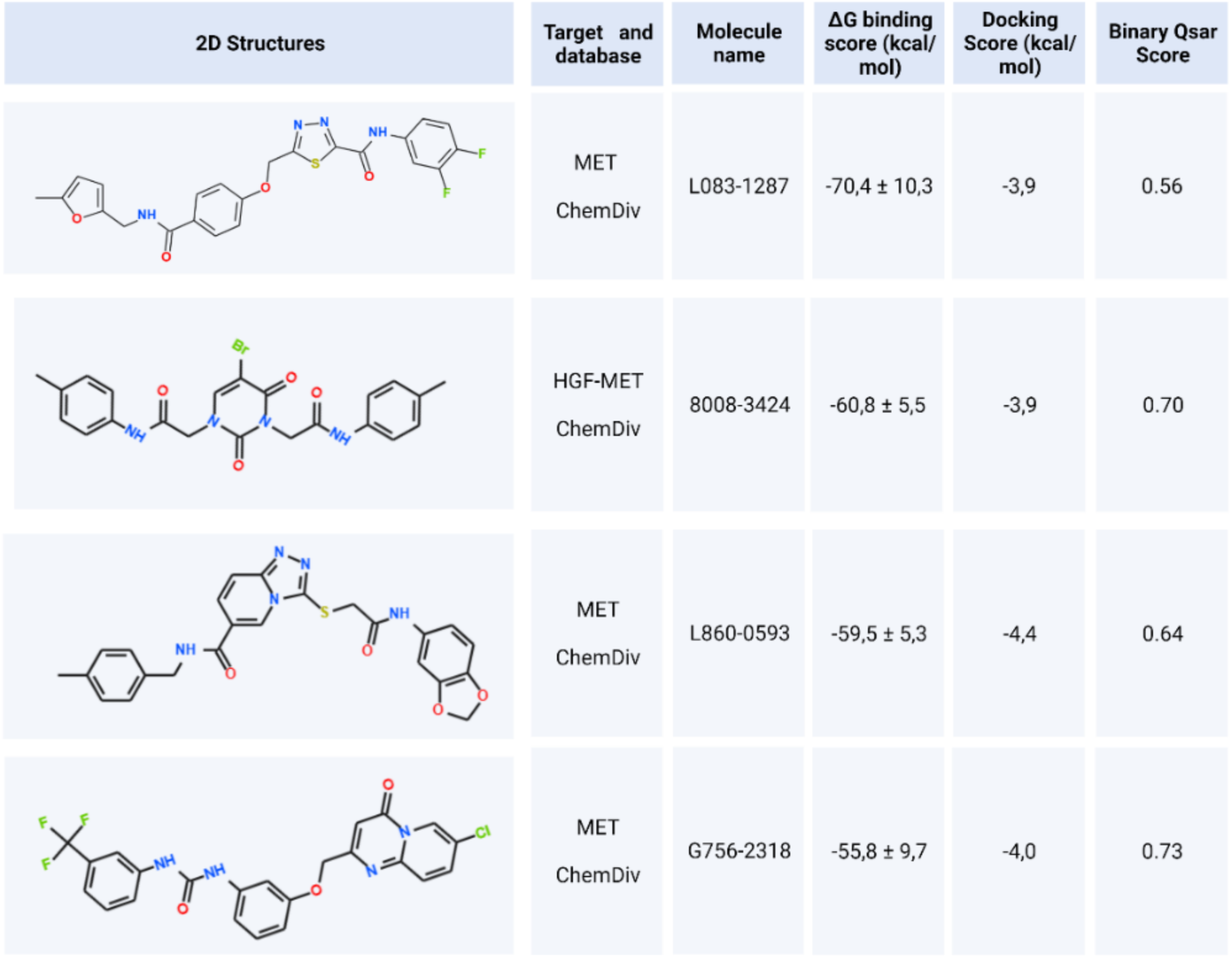
Candidate inhibitor molecules for inhibition of c-MET/HGF interaction obtained through Virtual Screening Workflow, short (10 ns) MD simulations, long (50 ns) MD simulations, and MM/GBSA from ChemDiv and Enamine libraries and their MM/GBSA and docking scores. Binary QSAR (Cancer-QSAR) therapeutic activity prediction results have also been presented. Cutoff is 0.5. Values higher than 0.5 indicate potentially active compounds. Training set consists of approved drugs. Model description: Training set N=886, Test set N=167, Sensitivity= 0.89, Specificity=0.83, Accuracy=0.86, MCC=0.72. Reference: Clarivate Analytics.

| 2D Structures | Target and database | Molecule name | $\Delta G$ binding score (kcal/mol) | Docking Score (kcal/mol) | Binary Qsar Score |
| --- | --- | --- | --- | --- | --- |
| | MET<br>ChemDiv | L083-1287 | $-70,4 \pm 10,3$ | -3,9 | 0.56 |
| | HGF-MET<br>ChemDiv | 8008-3424 | $-60,8 \pm 5,5$ | -3,9 | 0.70 |
| | MET<br>ChemDiv | L860-0593 | $-59,5 \pm 5,3$ | -4,4 | 0.64 |
| | MET<br>ChemDiv | G756-2318 | $-55,8 \pm 9,7$ | -4,0 | 0.73 |

### 2.5 Binding Free Energy Calculations (MM/GBSA)

Binding free energies were estimated using the Prime MM/GBSA module [19] in combination with the VSGB 2.0 implicit solvent model. [32] For the 10 ns trajectories, 100 frames were extracted from each simulation at 100 ps intervals, whereas for the 50 ns trajectories, 1,000 frames were sampled at 50 ps intervals. Calculations were carried out across all independent replicas, and the resulting values were averaged to obtain statistically robust estimates of ligand binding affinity. Compounds with mean binding free energies lower than −50 kcal/mol were classified as high-priority candidates for subsequent evaluation, with Z-score analysis used to support the selection process. (Figure S1)

### 2.6 sMD Simulations

To evaluate ligand dissociation energetics, sMD simulations were performed in GROMACS. [38] System setups of the sMD simulations were created by using CHARMM GUI with CHARMM36m force field. [39] Complexes were solvated in 0.15 M NaCl, energy-minimized with the steepest descent algorithm, and equilibrated under NPT conditions at 310 K with isotropic pressure coupling using the Berendsen barostat. Constant pulling forces were applied along a predefined unbinding pathway, with restraints of 20 kJ/mol applied to side-chain atoms and 400 kJ/mol to backbone atoms to preserve structural stability. The pulling speed and rate were adjusted according to the molecules as follows: For the pulling parameters, a spring constant of 100 kJ/mol/nm² and a pulling rate of 0.01 nm/ps were used. The thermostat used was Nose-Hoover, while the barostat was Parrinello-Rahman. The coupletype was set to isotropic, and the compressibility value was 4.5 × 10⁻⁵ bar⁻¹. The relaxation time was chosen as 1 ps. Free-energy differences were estimated from the pulling work using the Jarzynski equality. [40]

### 2.7 Binary quantitative structure-activity relationships (QSAR) Analysis

Binary QSAR predictions were carried out using the MetaCore/MetaDrug platform (Clarivate Analytics, https://portal.genego.com), which leverages large-scale datasets of small molecules and experimentally annotated biological activities to predict the functional profiles of previously uncharacterized compounds. Disease- and activity-specific QSAR models implemented within the platform were used to estimate the probability of relevant biological activity for the screened molecules. The cancer-focused QSAR model applied in this study was characterized by the following performance metrics: training set, *N* = 886; test set, *N* = 167; sensitivity = 0.89; specificity = 0.83; accuracy = 0.86; and MCC = 0.72. In parallel, toxicity risk was assessed using 26 distinct toxicity QSAR models available in MetaCore. For each toxicity endpoint, a predicted score above 0.5 was interpreted as indicative of potential toxicity. For the cancer activity model, compounds with scores of 0.5 or higher were classified as more likely to exhibit anticancer activity.

### 2.8 Cell Culture

HuH7 cells were cultured in Dulbecco’s Modified Eagle Medium (DMEM) and SNU-398 CMVGFP and METGFP cells were cultured in RPMI-1640 Medium supplemented with 10% fetal bovine serum (FBS), 1% L-Glutamine, 1% Penicillin/Streptomycin and 1% non-essential amino acids (Gibco). Cells were incubated at 37°C in a humidified atmosphere containing 5% CO₂. The HuH7 cell line (RRID: CVCL_0336) and SNU-398 cell line (RRID: CVCL_0077) were authenticated by single-tandem repeat (STR) profiling and routinely tested for mycoplasma contamination. For serum starvation, cells were incubated in DMEM supplemented with 2% FBS, 1% L-Glutamine, 1% Penicillin/Streptomycin, and 1% non-essential amino acids (Gibco) for 16 hours (h). All selected candidate c-Met inhibitors were dissolved in dimethyl sulfoxide (DMSO), thereby, DMSO was used in control groups when inhibitor treatment was omitted. HCC cells were starved for 16h followed by 2h induction with 10 ng/ml hepatocyte growth factor (HGF) (R&D Systems).Subsequently, cells were treated with the c-Met inhibitors at indicated times.

### 2.9 3-[4,5-dimethylthiazol-2-yl]-2,5-diphenyl tetrazolium bromide (MTT) Assay

The MTT assay was conducted as previously described. [41] Briefly, HuH7 and SNU-398 cells were seeded at a density of 4000 cells/cm² in 96-well plates. Adhered cells were starved for 16 hours, followed by induction with 10 ng/mL HGF for 2 hours. c-Met inhibitors were then added at concentrations of 3, 30, 300, 3000, 9000, and 30000 nM. Following 24h, 48h, and 72h inhibitor treatments, cells were incubated with 0.75 mg/ml MTT solution for 4h at 37 °C in 5% CO_2_ incubator. The formazan crystals formed during the incubation were solubilized in DMSO, and absorbance was measured at 570 nm using a Varioskan plate reader, with background correction at 720 nm. IC_50_ values for each inhibitor at the specified time points were calculated using GraphPad Prism v9.5.1. Inhibitor concentrations were transformed to their log10 values, and dose-response inhibition was assessed through non-linear regression analysis.

### 2.10 Colony Formation Assay

HuH7 cells were starved for 16 hours, counted, and seeded at a density of 80 cells/cm² into 24-well plates containing treatment media that includes 10 ng/mL HGF, and c-Met inhibitors at concentrations of 0.1, 1, and 5 μM. Treatment media was refreshed every 72 hours. DMSO was used in both the starvation and HGF control groups. Colony formation capacity was analyzed after 15 days of incubation, cells were fixed in methanol and stained with Giemsa dye. Plates were imaged using a BioRad ChemiDoc MP imaging system.

### 2.11 Western Blotting

Treated cells were lysed in ice-cold RIPA buffer (50 mM TrisCl pH 7.4, 150 mM NaCI, 1 mM EDTA pH 8.0, 1% NP-40) supplemented with protease inhibitor cocktail (Roche Diagnostics) and phosSTOP (Roche Diagnostics). The protein concentrations were determined using the bicinchoninic acid (BCA) assay (Thermo Scientific Pierce) according to the manufacturer’s instructions. Lysate samples (40 μg) were resolved on 5-8% SDS-PAGE gels, transferred onto membranes, and probed with antibodies against c-Met and the loading control Calnexin. Membranes were blocked with 5% bovine serum albumin (BSA) in TBS-Tween-20 (0.1%) for 1h at room temperature with gentle shaking. Primary antibodies against phospho-TYR1234-35 c-Met (1:1000, Abcam-ab278552), total c-Met (1:1000, Cell Signaling-CS3127), and Calnexin (1:1000, Santa Cruz-sc11397) were incubated in 3% BSA in TBS-T (0.1%) overnight at 4°C with gentle shaking. Secondary antibodies, goat anti-rabbit 680CW (1:7500, LICOR-926-68071) and goat anti-mouse AF800 (1:5000, Invitrogen-A11375), were incubated at room temperature for 2 hours. Immunoblots were visualized using multiplex imaging at DyLight 680 and DyLight 800 channels in BioRad ChemiDoc MP.

### 2.12 Zebrafish Toxicity Assay

Zebrafish embryos were dechorionated at 2 days post-fertilization (dpf) and incubated at E3 medium in 34°C incubator with the c-Met inhibitors (L083-1287, G756-2318 and 8008-3424) at concentrations ranging from 4X IC_50_, 2X IC_50_, IC_50_, 0.5X IC_50_, and 0.25X IC_50_. Specifically, zebrafish embryos were incubated with L083-1287 (65, 130, 260, 520, and 1040 nM); G756-2318 (1419, 2839, 5678, 11356, and 22712 nM); 8008-3424 (1599, 3198, 6397, 12794 and 25588 nM). E3 medium only and E3 medium with DMSO served as control groups. The media was refreshed in every 24 hours, and the experiment was concluded at 5 dpf. Throughout the study, embryos were monitored daily for viability, and dead zebrafish were recorded and removed from the media. At the end of the experiment, surviving embryos were assessed for developmental defects, internal bleeding, and cardiac edema. All zebrafish were fixed and analyzed using a stereomicroscope with a 4X objective. Survival analysis was conducted using GraphPad Prism v9.5.1, and representative images were prepared using Photoshop 26.2.0. The zebrafish experiments were performed as a part of service procurement at Izmir Biomedicine and Genome Research Center Zebrafish Facility. According to EU Directive 2010/63/EU on the protection of animals used for scientific purposes, early life-stage animals (including zebrafish up to 5 dpf) are not defined as protected. Since the experiments were terminated at 5dpf, ethical approval is exempted for this study.

### 2.13 Surface Plasmon Resonance (SPR) Analysis

All experiments were conducted using Biacore T200 with a CM5 chip at 25 °C. Recombinant cMet-ECD (Acro Biosystems, Catalog Number: MET-H5227) was used as the ligand and small molecules were used as analytes. Flow cell (FC) 1 was used as the reference for FC2. Anti-his antibody (Cytiva Catalog Number: 29234602, 1 mg/ml stock concentration) was diluted 1:20 in 10 mM acetate buffer at pH 4.5 and immobilized onto both FCs to a level of ∼12000 RU, using standard amine coupling chemistry. PBS-P (20 mM phosphate buffer pH 7.4, 137 mM NaCl, 2.7 mM KCl, 0.05% surfactant P20) was used as the immobilization running buffer.

cMet-ECD was diluted in PBS-P+1%DMSO to 0.05 mg/ml concentration and captured onto FC2 to a level of 730 RU. PBS-P+1%DMSO was used both as the capture running buffer and the kinetics running buffer. Based on the ligand immobilization level the calculated theoretical R_max_ value was 3.4 RU. The flow rate was maintained at 50 μL/min. The contact and dissociation for analyte bindings were 60s and 300s, respectively. One 20s pulse of glycine pH 1.5 was injected for surface regeneration. This regeneration condition takes away all captured ligands from the anti-his antibody surface. Therefore, fresh ligand was captured at the beginning of each cycle. Injected analyte concentrations were from 40 μM down to 5 μM (two-fold dilution). All analytes were injected in duplicate. K_D_ values were calculated using a 1:1 binding model in BiaEvaluation software. Double reference (buffer alone injection and FC1 reference surface binding) subtraction was used to account for any non-specific interaction that may occur between the small molecules and chip surface.

### 2.14 First Lead Optimization Cycle: Similarity Screening of Analogs

To identify structural analogs of the selected lead compounds (i.e., 8008-3424, L083-1287), similarity-based virtual screening was performed using the SwissSimilarity web server (http://www.swisssimilarity.ch/). The ZINC drug-like database, consisting of 9,205,113 commercially available compounds, was queried using two complementary methodologies: three-dimensional electroshape-based similarity and two-dimensional FP2 fingerprint-based similarity searches. For each lead compound, 400 structurally related molecules were retrieved, yielding a total of 800 candidates. No pre-filtering based on physicochemical properties was applied at this stage to maintain chemical diversity within the dataset. Docking simulations for all retrieved analogs were conducted with the Glide/XP flexible docking algorithm, which allowed ligands to explore multiple binding poses within the receptor pocket. Conformational sampling was enhanced to increase coverage of the potential binding space. Binding free energies of the top-ranked docking poses were estimated using Prime MM/GBSA calculations, applied to the ten best poses for each ligand. Compounds with favorable binding scores were subsequently subjected to MD simulations. For these MD simulations, previously described parameters in MD simulations were employed. Each selected analog was simulated for 10 ns, with five independent replicas generated using different random seeds to improve sampling and reduce bias. Trajectory frames were extracted every 100 ps, and MM/GBSA calculations were performed across 1000 frames per replica to evaluate average binding free energies. Compounds exhibiting binding free energies below the predefined threshold (-50 kcal/mol) were advanced to extended simulations. In these cases, 50 ns MD simulations were performed in three replicas per ligand, and frames were extracted every 50 ps to provide higher-resolution energy profiles. MM/GBSA binding free energy calculations were applied to these extracted frames per simulations.

### 2.15 Second Lead Optimization Cycle

Based on the *in silico* and experimental outcomes of the first optimization phase, L083-0077 was selected as the core scaffold for the second lead optimization cycle. Structural analogues of L083-0077 were retrieved from MolPort (https://www.molport.com) and PubChem databases (https://pubchem.ncbi.nlm.nih.gov/), excluding any compounds already investigated in the first cycle. A total of 303 candidate molecules were subjected to Glide XP docking, and the resulting poses were rescored using Prime MM/GBSA calculations. Using the MM/GBSA score of L083-0077 as the threshold, 34 compounds with higher predicted affinity were selected and evaluated by short 25 ns MD simulations (5 independent replicas) followed by MM/GBSA analysis over 250 frames as a filtering step. From these, 18 ligands exhibiting superior affinity relative to L083-0077 were advanced to 200 ns production MD simulations. Following the same filtering procedure, the top three candidates were subsequently subjected to Pulling MD simulations using a spring constant of 100 kJ mol⁻¹ nm⁻² and a pulling rate of 0.01 nm/ps, in order to assess their mechanical binding stability.

### 2.16 Fragment-Based Binding Free Energy Decomposition of Lead Compounds

To quantitatively dissect the binding behavior of the lead compounds L083-1287 and L083-0077, we employed a ligand-fragmentation strategy that partitioned each scaffold into structurally and energetically meaningful subunits. Fragmentation was guided by the local interaction capacity of binding-pocket residues, with the aim of preserving ligand regions capable of sustaining rigid and chemically diverse contacts, including electrostatic, hydrogen-bonding, hydrophobic, and aromatic interactions. This design allowed each fragment to represent a position-specific binding module with distinct recognition potential. MM/GBSA calculations were subsequently carried out for each fragment independently using trajectory data from three separate 200 ns MD simulations. This fragment-resolved energy decomposition enabled quantitative identification of the ligand substructures that contribute most strongly to binding stability and thus informed structure-based optimization of the lead scaffolds.

### 2.17 Diffusion-Based Ligand Efficiency Assessment with Reference to Collision Theory

Diffusion coefficient calculations were carried out using the diffusion analysis tool implemented in the Desmond module of the Schrödinger Suite. For each candidate inhibitor, the ligand was extracted from its corresponding protein–ligand complex and simulated as an isolated molecule in bulk solvent.[42] Each system was then subjected to 500 ns MD simulations, from which the self-diffusion coefficient was determined. [43,44] Ligand mobility was further characterized by monitoring the time evolution of the mean square displacement (MSD) [45], and heat map representations were generated to provide a more detailed view of temporal diffusion behavior.

### 2.18 Ligand-Based Similarity-Driven Specificity Prediction via Off-Target Affinity Projection

Following the processing of data obtained from the SEA web server (https://sea.bkslab.org/) 3D PDB structures of proteins associated with the identified off-targets were examined to explore potential cross-reactivity of the lead compounds. [46] Binding regions were first analyzed using the co-crystallized inhibitor-protein complexes, and the same sites were subsequently screened with the lead compounds. To benchmark the binding behavior, positive control molecules were re-docked into their native binding sites using Glide XP, and comparative docking experiments were performed with the lead compounds. Lead compounds were then subjected to three independent 20 ns MD simulations, followed by MM/GBSA free energy calculations to assess their relative binding affinities and stabilities. In addition, for targets, where no co-crystallized ligand structure was available, a comprehensive surface-based screening approach was employed to gain insights into their potential specificity profiles, detect possible allosteric effects, and evaluate the aggregation-collapse tendencies of the protein in response to lead compound binding. To achieve this, each candidate off-target was individually subjected to 100 short (10 ns) MD simulations initiated with different random seeds. The resulting trajectories were clustered using TTClust [47], and 30 representative frames per target were extracted. From each representative structure, potential binding pockets were identified using MDpocket [48], followed by size-based filtering and pocket enrichment analysis to define viable ligand-binding regions. Grid boxes were then generated to cover all such pockets across the selected frames. Each lead compound was docked into every frame-pocket combination using two complementary docking strategies Glide SP (physics-based) and Gnina (AI-based). [49] Docking poses exceeding a predefined binding threshold were retained, subjected to MD simulations, and evaluated using MM/GBSA binding free energy calculations. The resulting energetic profiles provided a quantitative basis for inferring ligand specificity in terms of relative affinities toward predicted off-targets.

### 2.19 *In Silico* Evaluation of hERG Blocking Potential of Lead Compounds

To assess the potential cardiotoxicity of the lead compounds L083-1287 and L083-0077, their propensity to inhibit the hERG potassium channel was evaluated. *In silico* prediction of hERG liability was performed using the Pred-hERG web server, which employs QSAR-based classification models trained on experimentally validated datasets and implemented through artificial intelligence algorithms. [50] In addition, a comparative structure-based analysis was carried out using three established hERG blockers, E-4031 [51], pimozide [52], and astemizole [53], to examine their structural similarity and interaction characteristics. Based on these analyses, astemizole was selected as the reference hERG blocker for subsequent molecular-level investigations. Astemizole, together with L083-1287 and L083-0077, was then subjected to three independent 100 ns MD simulations, followed by MM/GBSA binding free-energy calculations. The resulting interaction profiles and binding-energy estimates were compared with literature-defined key residues implicated in hERG blockade, enabling a comprehensive assessment of the lead compounds relative to a representative hERG inhibitor.

### 2.20 Analysis of Ligand Binding Behavior under pH-Relevant Physiological Conditions of the Tumor Microenvironment

To investigate the impact of tumor-associated acidosis on ligand binding, receptor models were prepared under two physiologically relevant acidic conditions representative of the tumor microenvironment, namely pH 6.0 ± 1.0 and pH 5.5 ± 0.5. [54,55] These pH ranges were selected to reflect the EC acidification commonly observed in solid tumors as a consequence of hypoxia, altered metabolic flux, and dysregulated respiratory activity. For each protonation condition, receptor structures were prepared accordingly and used as docking templates for ligands previously prioritized through integrated *in silico*, *in vitro*, *in vivo*, and SPR-based validation. The resulting ligand–receptor complexes were then subjected to 50 ns MD simulations in triplicate for each system to examine the stability and adaptability of binding under acidic conditions relevant to the cancer milieu. To ensure robust sampling of the conformational ensemble, 1000 evenly distributed frames were extracted from each trajectory and analyzed by MM/GBSA to estimate binding free energies across the tested pH conditions. This workflow enabled a quantitative comparison of ligand affinity and binding stability as a function of acidification, thereby providing insight into the extent to which tumor-relevant pH variation, typically spanning pH 5.5–6.5, modulates ligand–target recognition and complex persistence.

### 2.21 Extended MD simulations

To further interpret the experimental findings obtained from the *in vitro* and *in vivo* assays, extended MD simulations were performed for seven prioritized lead compounds. Each ligand–protein complex was subjected to 200 ns all-atom MD simulations in order to evaluate the long-timescale stability of binding, persistence of key intermolecular contacts, and ligand-induced conformational responses of the receptor. For the most promising lead compound, L083-0077, the simulation time was further extended to 500 ns to enable a more comprehensive assessment of binding durability, conformational adaptability, and the reproducibility of its interaction network over an expanded temporal window. All simulation systems were prepared and analyzed using the same protocol described in Section 2.4.

### 2.22 Elucidation of the c-MET Inhibition Mechanism and Mechanistic Validation of Lead Compound Activity

To define the molecular basis of c-MET inhibition, normal mode analyses (NMA) were performed following a comprehensive review of the structural literature, with particular attention to unresolved questions regarding binding-site localization and the putative molecular clamp mechanism governing the transition of c-MET from the active, HGF-bound state to an inactive conformation. The cryo-EM-derived monomeric structure of the c-MET ectodomain (PDB ID: 7MO7) [12] was used as the primary reference. To capture the broader conformational landscape of the receptor, this structure was analyzed together with additional EC-domain models, including the AlphaFold3-predicted full-length ectodomain model encompassing residues 25–925, the experimentally resolved structure spanning residues 25–567 (PDB ID: 1SHY), and the structure covering residues 42–658 (PDB ID: 2UZX). [56–59] Integrating these complementary structural models enabled evaluation of large-scale domain motions and facilitated identification of conformational features potentially associated with receptor autoinhibition. To further investigate the structural transition from the extended to the collapsed state, and to determine whether the inhibitory behavior of the experimentally validated lead compound was consistent with the intrinsic regulatory dynamics of the receptor, extensive MD simulations were performed. Specifically, eight independent 250 ns MD simulation replicas were carried out to characterize the reproducibility and stability of conformational transitions across distinct trajectories. This ensemble-based approach allowed us to examine whether ligand binding reinforced naturally accessible inactive-state motions or induced alternative inhibitory conformational states. Collectively, these analyses provided a mechanistic framework for linking lead-compound binding to the dynamic regulation of c-MET ectodomain architecture and inhibitory function.

### 2.23 Sequential Angular Hinge-Rotation Mechanism

The c-MET ectodomain contains multiple hinge regions, regulatory loops, and disulfide-constrained segments that together confer substantial conformational plasticity while preserving overall structural integrity. This combination of flexibility and architectural stability enables large-scale domain rearrangements that are likely to be functionally coupled to receptor activation and inhibition. Accordingly, for a target such as c-MET, conventional energy-based analyses are informative primarily as an initial filtering step and are insufficient on their own to resolve the mechanistic basis of conformational regulation. To characterize these dynamic transitions in greater detail, and to determine whether the same intrinsic motions remain accessible in the presence of bound inhibitors, metadynamics simulations were performed in Desmond using the OPLS3e force field, starting from the initial frame of the 7MO7 structure. To capture the principal conformational motions associated with receptor rearrangement, collective variables (CVs) were defined on the basis of domain-level rotational and translational movements. In total, four centers of mass were used to parameterize the metadynamics framework. A dihedral CV was first constructed to monitor angular motion around the Gly517–Gly519 region, which was hypothesized to act as a local hinge during the transition between extended and more compact receptor states. This dihedral incorporated atoms from residues Val184, Leu420, His251, Ala320, Ile442, Ile491, and Val55 located in the central region of the SEMA domain, together with Gly517 and Gly519, and residues Lys567, Ile583, Val603, Lys623, Thr612, Ile639, and Gln648 positioned in the central portion of the IPT1 domain between the PSI domain and Helix26. (Figure S2) This CV was therefore designed to track relative rotational displacement between the SEMA and IPT1 regions across the putative hinge axis. The Gaussian width for this dihedral CV was set to 5.0°. As a second descriptor of conformational progression, a distance CV was defined between Lys306 (atom 4436; hydrogen) and Glu616 (atom 9252; negatively charged oxygen). This coordinate was selected to monitor domain closure and the formation of long-range intramolecular contacts associated with the transition from a more extended to a more collapsed structural state. The width parameter for this distance CV was set to 0.05 Å. Together, the dihedral and distance CVs enabled simultaneous tracking of rotational and compaction-related motions, thereby providing a mechanistically interpretable representation of the conformational landscape. Metadynamics simulations were carried out at 310 K with a Gaussian hill height of 0.03 kcal/mol and a deposition interval of 0.09 ps. To enhance exploration of the free-energy surface, reduce the likelihood of kinetic trapping, and improve sampling of alternative conformational states, well-tempered metadynamics was applied with bias modulation across different energy levels. This setup allowed systematic interrogation of the angular hinge-rotation behavior underlying c-MET conformational regulation and enabled assessment of whether ligand binding preserves, perturbs, or redirects these intrinsic mechanistic motions. To complement the metadynamics analysis, local energetic frustration was evaluated using Frustratometer 2.0. This analysis was used to identify residue networks under competing energetic constraints and to map regions in which conformational transitions may be facilitated by locally frustrated interactions. Integration of metadynamics-derived conformational sampling with frustration analysis therefore provided a mechanistic framework for defining the sequential hinge-rotation behavior of c-MET and for assessing how inhibitor binding may reshape this intrinsic inhibitory landscape.

### 2.24 Holo-Apo Sequencial Mechanism Comparison

Holo and apo metadynamics trajectories were analyzed using an identical set of CVs to allow direct comparison of their underlying mechanistic landscapes. To characterize long-range interaction networks across the full protein surface, frustration analysis was performed using the Frustratometer 2.0 workflow [60], and time-resolved frustration profiles were monitored throughout the simulations. Representative trajectories were selected from independent metadynamics runs that converged on similar mechanistic features and were subsequently used for downstream analysis. For each simulation frame, residue-level interaction energies were computed using the AWSEM (Associative Memory, Water-Mediated, Structure-Based Energy Model) coarse-grained formalism, as implemented in Frustratometer 2.0 through a LAMMPS backend. On the basis of these energy calculations, both configurational and mutational frustration indices were determined. In this representation, each residue was modeled using its C_α_ atom to capture backbone geometry, a C_β_ pseudo-atom to encode side-chain properties, and the backbone carbonyl oxygen to account for hydrogen-bonding contributions, while the remaining atomic degrees of freedom were treated implicitly within the energy function. Residue–residue interactions were evaluated using a 14.5 Å cutoff. Interaction pairs that recurred most frequently across the trajectories were identified, and the top 100 highest-occurrence pairs were selected for further analysis. These dominant interaction networks were then examined together with locally highly frustrated regions to identify spatial clustering patterns and to define potential pivot points around which mechanistically relevant conformational rearrangements are organized.

## 3. Results

### 3.1 Elucidation of Promising Hit Molecules through Virtual Screening

To comprehensively interrogate the chemical space surrounding potential small-molecule inhibitors of the c-MET/HGF signaling axis, we established a multistage virtual screening pipeline that integrated broad library coverage, chemically informed preprocessing, stringent property-based filtering, hierarchical docking, and energy-based rescoring. The screening campaign was initiated using two complementary and chemically diverse commercial chemical libraries: ChemDiv’s 300k Representative Compounds Bemis–Murcko Clustering Algorithm Library and the Enamine Hit Locator Library, which initially comprised 300,960 and 460,160 compounds, respectively. These libraries were selected to maximize both scaffold diversity and screening tractability, thereby enabling broad exploration of drug-like chemical space while minimizing redundancy at the core chemotype level. Before docking, both libraries were subjected to extensive ligand preparation to generate chemically realistic molecular representations under biologically relevant conditions. Using LigPrep-based preprocessing, each parent entry was expanded to account for stereochemical, tautomeric, and protonation-state variability that could influence molecular recognition at the target interface. Specifically, stereoisomers were enumerated where applicable, alternative tautomeric states were generated, and protonation states consistent with physiological pH were assigned. This expansion step was intended to improve the fidelity of docking by ensuring that each compound was represented in forms likely to exist under assay-relevant conditions. As a result, the ChemDiv collection increased from 300,960 to 411,812 structures, whereas the Enamine collection expanded from 460,160 to 652,588 entries, substantially broadening coverage of accessible conformational and ionization space. The expanded libraries were then subjected to a sequential curation workflow designed to retain chemically tractable and pharmacologically relevant molecules while excluding liabilities that could compromise downstream prioritization. As an initial filtering step, QikProp analysis was used to calculate an extensive set of physicochemical and ADME-relevant descriptors, including lipophilicity, aqueous solubility, molecular weight, hydrogen-bond donor/acceptor capacity, and related features associated with drug-likeness. This first-pass filter minimally reduced the ChemDiv library to 407,082 compounds, indicating that the starting collection was already enriched in molecules with broadly acceptable physicochemical profiles. More restrictive filters were then applied by enforcing Lipinski’s Rule of Five and removing compounds bearing reactive, unstable, or otherwise problematic functional groups. These sequential criteria reduced the ChemDiv dataset to 272,401 compounds. A comparable filtering workflow was applied to the Enamine-expanded library, reducing it from 652,588 to 611,687 structures. Importantly, this refinement process was not intended simply to reduce library size, but to improve the signal-to-noise ratio of the screening campaign by enriching for molecules with a more favorable balance of chemical stability, developability, and target engagement potential, while still preserving broad scaffold-level diversity. The refined compound sets were subsequently docked against three independently constructed receptor grids designed to represent distinct mechanistic intervention points within the c-MET/HGF signaling system: the NK1 domain of HGF, the SEMA domain of c-MET, and the composite c-MET/HGF interface. These three docking scenarios were selected to capture complementary pharmacological strategies, including direct disruption of ligand recognition, blockade of receptor-side hotspot engagement, and perturbation of the broader protein–protein interaction (PPI) surface required for productive c-MET activation. By distributing the screening campaign across these mechanistically distinct receptor environments, we aimed to identify not only conventional binders to localized hotspot regions but also compounds capable of modulating interface stability or altering the geometry of ligand–receptor recognition. For the ChemDiv library, 272,401 filtered compounds were first subjected to Glide high-throughput virtual screening (HTVS), a rapid initial docking stage used to eliminate compounds with the weakest predicted binding performance. This step reduced the library to 128,041 molecules, which were subsequently advanced to Glide standard-precision (SP) docking for more refined pose generation and scoring. From this set, 11,130 compounds progressed to extra-precision (XP) docking, that provided a higher-stringency evaluation of binding geometry and interaction complementarity. The XP stage yielded 4,393 HGF-directed candidates, 4,013 c-MET-directed candidates, and 2,724 interface-directed candidates. These top-ranking docking poses were then rescored using single-pose MM/GBSA calculations to incorporate an additional energetic assessment beyond empirical docking scores alone, enabling more discriminating prioritization of putative binders. This rescoring step ultimately reduced the ChemDiv-derived candidate pool to 1,112 compounds. An analogous hierarchical workflow was applied to the Enamine collection, resulting in 18,348 compounds reaching the XP docking stage, of which 1,848 were retained following MM/GBSA-based refinement. To further improve prioritization robustness and minimize false-positive enrichment, we implemented a hybrid ranking framework that integrated empirical docking scores, MM/GBSA-derived binding free-energy estimates, and Z-score normalization across the screened datasets. This combined strategy allowed us to identify compounds that were not only highly ranked in absolute terms, but also statistically enriched relative to the broader scoring distributions within each screening branch. Such normalization was particularly important given the use of multiple receptor grids and independent library subsets, where raw score comparisons alone can be misleading. By integrating orthogonal scoring layers with distribution-based normalization, we sought to prioritize compounds showing both strong predicted affinity and reproducible performance across the screening workflow. Using this multilevel selection strategy, we ultimately identified 252 high-confidence candidate molecules across the three docking scenarios, including 83 compounds prioritized against HGF, 84 against c-MET, and 85 against the c-MET/HGF interface. These compounds represented the most compelling subset emerging from the large-scale screening campaign, as they combined favorable predicted binding energetics, acceptable physicochemical characteristics, and robust statistical support. Collectively, this workflow enabled progressive reduction of more than one million ligand representations into a focused set of mechanistically diverse and computationally prioritized candidates suitable for downstream MD simulations and experimental validation. (Figure 2)

**Figure 2.**
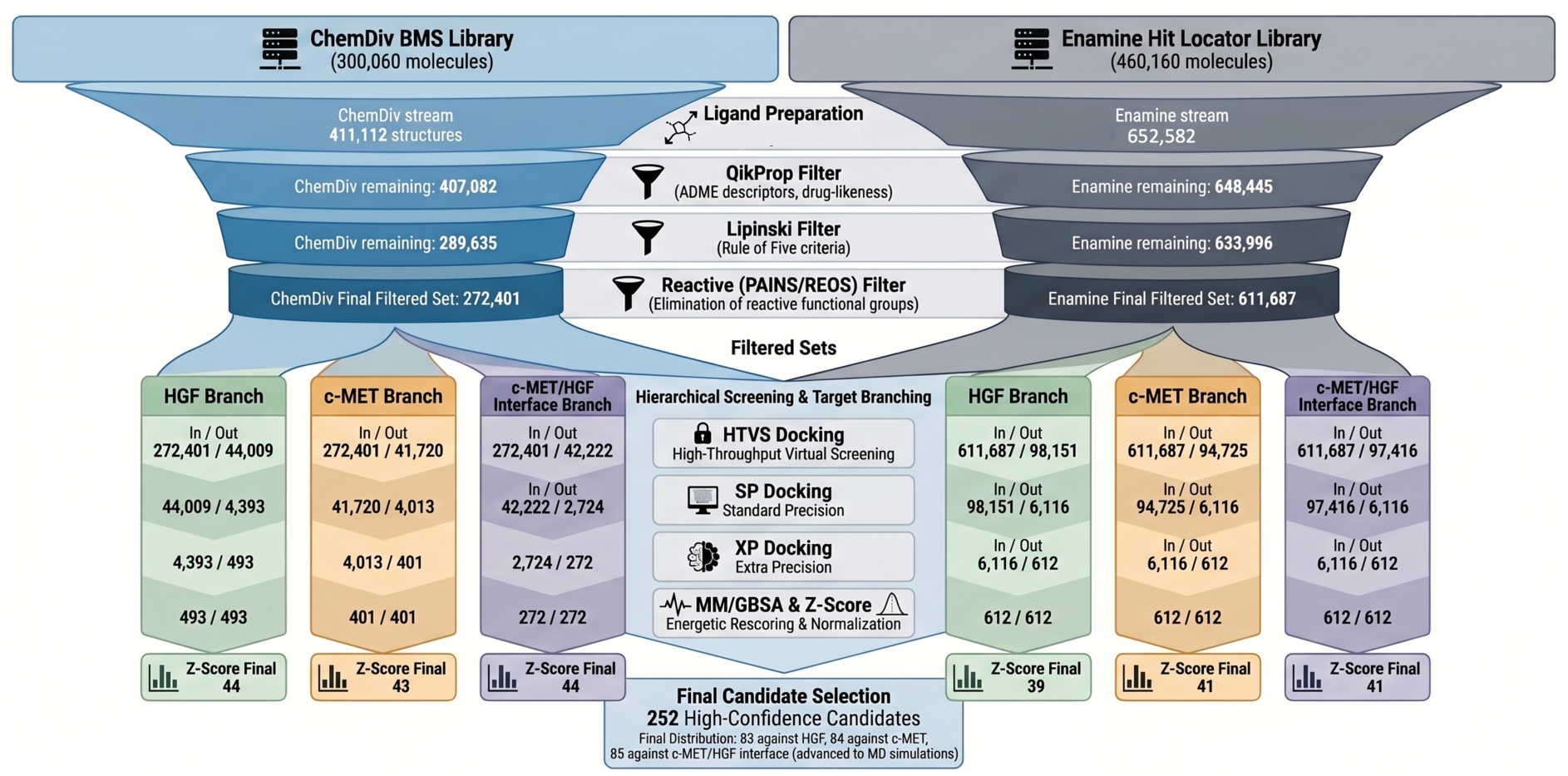
The diagram illustrates the stepwise application of various filtering criteria and virtual screening algorithms on molecules from the Enamine Hit Locator and ChemDiv BMS libraries. Molecules that successfully pass each stage proceed to the subsequent step of the workflow. The target binding sites are defined as c-MET (Phe393, Asn398), HGF (Lys47, Lys91, His112, His114), and the c-MET/HGF complex (Phe393, Asn398, Lys47, Lys91, Phe112, His114).

### 3.2 MD Simulations and MM/GBSA Calculations

Following prioritization of 252 high-confidence compounds from the virtual screening campaign, all-atom MD simulations were performed to evaluate the dynamic stability, binding persistence, and conformational compatibility of the resulting protein–ligand complexes under near-physiological conditions. As an initial post-docking refinement step, each compound was subjected to 10 ns MD simulations performed in five independent replicas using distinct random seed assignments. This replicate-based design was implemented to reduce sensitivity to initial coordinates and stochastic effects, thereby providing a more statistically robust representation of ligand behavior within the three docking environments, namely the c-MET SEMA region, the HGF NK1 region, and the broader c-MET/HGF interface. Across most systems, the equilibration phase was completed within approximately 2-4 ns, after which potential energy profiles and major structural fluctuations reached stable regimes suitable for comparative analysis. To quantify ligand retention and energetic favorability, frame-based MM/GBSA calculations were carried out using 100 trajectory frames per simulation. This first MD filtering stage allowed discrimination between compounds that retained stable engagement with key hotspot residues and those that rapidly lost favorable contact patterns after docking-based placement.

On the basis of these short-timescale simulations, 31 compounds among 252 ligands were selected for extended 50 ns MD simulations performed in triplicate. This second simulation tier was designed to provide a deeper assessment of the temporal persistence of intermolecular interactions, the adaptability of each ligand to local pocket flexibility, and the reproducibility of binding behavior over longer timescales. The longer trajectories also enabled more reliable characterization of whether ligand binding was maintained through stable anchoring interactions or instead reflected transient, docking-derived poses that were not sustained dynamically. Following experimental prioritization through *in vitro* and *in vivo* analyses, seven lead compounds (L083-1287, 8008-3424, L860-0593, G756-2318, 8016-2998, Y043-7353, and Y020-8864) were subsequently subjected to 200 ns MD simulations to enable molecular-level interpretation of their biological activity. (see Table S3) These long-timescale simulations were used to resolve persistent contact networks, ligand-induced conformational responses, and the durability of binding modes that could plausibly explain the observed pharmacological behavior.

This multistage simulation workflow progressively refined the candidate set and identified four compounds as the most promising leads: L083-1287, 8008-3424, L860-0593, and G756-2318. Among these, L083-1287 exhibited the most favorable average binding free energy, reaching values as low as −70.4 kcal/mol, consistent with a highly stable binding mode. Structural interaction analysis further showed that L083-1287 formed a persistent interaction network within the c-MET SEMA domain, engaging residues Phe241, Tyr245, Leu269, Cys385, Gln387, and Asn399. The recurrence and stability of these contacts suggest a well-defined anchoring mechanism that stabilizes the ligand within the pocket and supports prolonged occupancy throughout the simulation interval. (Figure 3) Particularly notable was the repeated involvement of Cys385 and Gln387, which emerged as energetically and structurally important binding determinants despite being less emphasized in earlier hotspot-focused models.

**Figure 3.**
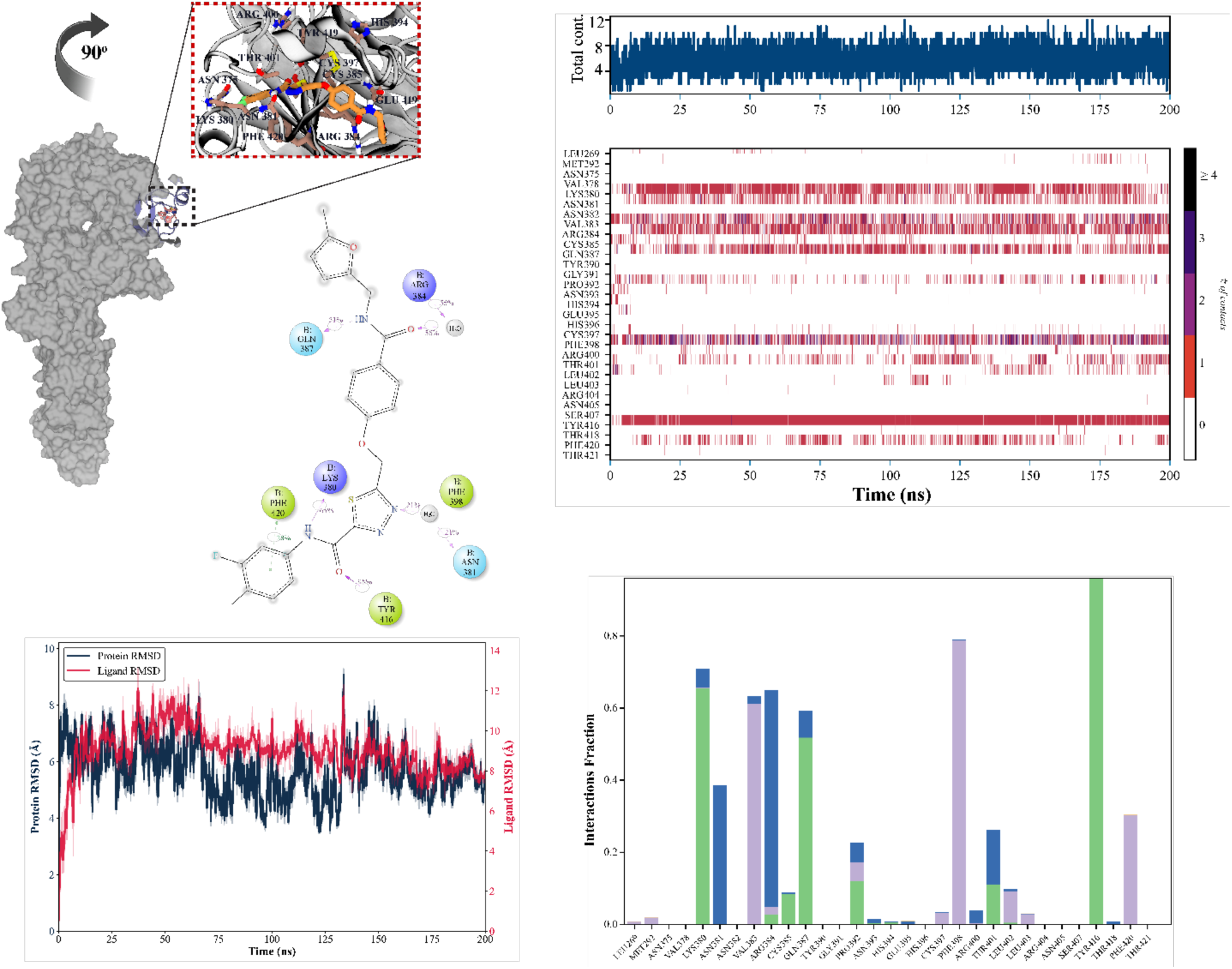
(A) 3D and 2D ligand interaction of diagrams of the selected hit compound (L083-1287) at the MET site. (B) Numeric residue contact map of the protein-ligand complex; x axis shows time by unit of the ns; y-axis shows residue types of the c-MET. Contact bar represents contact number of the residue-ligand (C) Protein-Ligand RMSD; protein RMSD values were plotted by backbone atoms, Ligand RSMD values were plotted by all atoms. x-axis illustrates time (ns), y-axis represents protein and ligand RSMD by unit of Å. (D) Interaction fraction values of the ligand for residues of the c-MET. x-axis depicts residue of the c-MET and y-axis indicates the interaction fraction, defined as the fraction of MD frames in which each residue forms a contact with the ligand.

In contrast to the more pocket-confined binding profile of L083-1287, compound 8008-3424 displayed a broader, dual-interface interaction pattern. This ligand simultaneously engaged c-MET residues Phe393 and Asn398 and HGF residues Lys47, Lys91, Phe112, and His114, indicating that its activity may arise from direct perturbation of the receptor–ligand recognition interface rather than occupation of a single localized site. (Figure S3) Such dual-site engagement is mechanistically important because it suggests the possibility of stabilizing a nonproductive interface geometry or sterically disrupting the contact network required for efficient HGF-dependent receptor activation. Additional simulation and interaction analyses for other lead compounds are presented in the Supplementary Information (Figures S4–S9), and the corresponding docking scores, MM/GBSA-derived binding free energies, and predicted anticancer activity values are summarized in Table 1.

Taken together, these data demonstrate that integration of MD simulations with MM/GBSA rescoring provides a stringent and mechanistically informative framework for distinguishing transient docking hits from ligands capable of sustaining stable, biologically meaningful interaction networks. Beyond simple ranking by binding energy, the simulation workflow revealed ligand-specific modes of engagement, differentiated pocket-binding from interface-bridging behavior, and identified previously underappreciated residues such as Cys385 and Gln387 as potential structural hotspots for extracellular c-MET targeting. These findings not only strengthened prioritization of the lead series for experimental testing but also provided an early mechanistic basis for subsequent hit-to-lead optimization.

The lead compound L083-1287 exhibited a well-defined and stable binding profile within the active pocket of the target protein throughout the 200-ns MD simulations. RMSD analysis showed that, following the initial equilibration phase (∼10 ns), the protein backbone maintained overall structural integrity with fluctuations between 2-6 Å, while the ligand RMSD stabilized around 6-8 Å, indicating a consistent binding orientation without any dissociation from the pocket. (Figure 3) These observations confirm that the ligand dynamically adapts to the flexibility of the binding cavity while preserving its overall stability. The interaction timeline revealed several residues forming persistent contacts with the ligand, including Asn393, Phe398, Tyr404, Arg404, Asn405, Asn425, and Lys380. Among these, Asn393 and Phe398 displayed high contact occupancies (>80%) and were identified as critical hotspot residues maintaining the ligand’s anchoring position within the binding interface. (Figure 3) The 2D interaction diagram indicated that the ligand established dual hydrogen bonds with Asn393 and engaged in π-π stacking interactions with Phe398, while Tyr404 and Arg404 contributed additional electrostatic stabilization. This combination of polar and hydrophobic contacts suggests a hybrid binding mechanism, where aromatic stacking and hydrogen bonding cooperatively reinforce the interaction network. The interaction fraction histogram further supported these findings, showing consistently high contact frequencies along the Asn393-Phe398-Tyr404 axis, which forms the core interaction triad governing the ligand’s positional stability.

Overall, these results demonstrate that L083-1287 binds with high affinity and conformational stability, forming a persistent interaction network that anchors the ligand firmly within the binding pocket. The Asn393 and Phe398 residues act as dominant hotspot regions, ensuring the durability and specificity of binding. Collectively, these data position L083-1287 as a strong and thermodynamically favorable lead candidate for further optimization in the inhibitor design pipeline.

### 3.3 sMD Simulations

To complement equilibrium simulations, sMD was performed on the seven best candidates identified from long-term MD simulations. The goal of this analysis was to assess the mechanical strength of ligand binding and the relative resistance of protein-ligand complexes to external perturbation. Each complex was subjected to a pulling force of 100 kJ/mol/nm applied along the ligand exit pathway, and the dissociation trajectories were analyzed over an average duration of 450-500 ps. The resulting force-time profiles revealed striking differences in binding resilience. Compounds such as 8008-3424, Y020-8828, and L083-1287 demonstrated exceptional resistance to forced dissociation at the c-MET-HGF interface, consistent with their dual-site binding mechanisms and broad engagement of interface residues. (Figure 4) In contrast, molecules including L860-1860 and 8016-2998 dissociated more readily, suggesting weaker mechanical stability despite favorable equilibrium free energies. Notably, 8008-3424 exhibited incomplete dissociation within the standard simulation timeframe, requiring extended force application to achieve separation of the ligand. This observation highlights the potential of 8008-3424 to act as a highly stable inhibitor capable of withstanding conformational fluctuations and competitive binding pressures within the tumor microenvironment. These findings underscore the value of integrating sMD into the inhibitor discovery workflow. While equilibrium simulations identify favorable energetic profiles, sMD provides complementary insight into the kinetic resilience of binding, thereby discriminating between compounds with similar free energy values but divergent mechanical stabilities.

**Figure 4.**
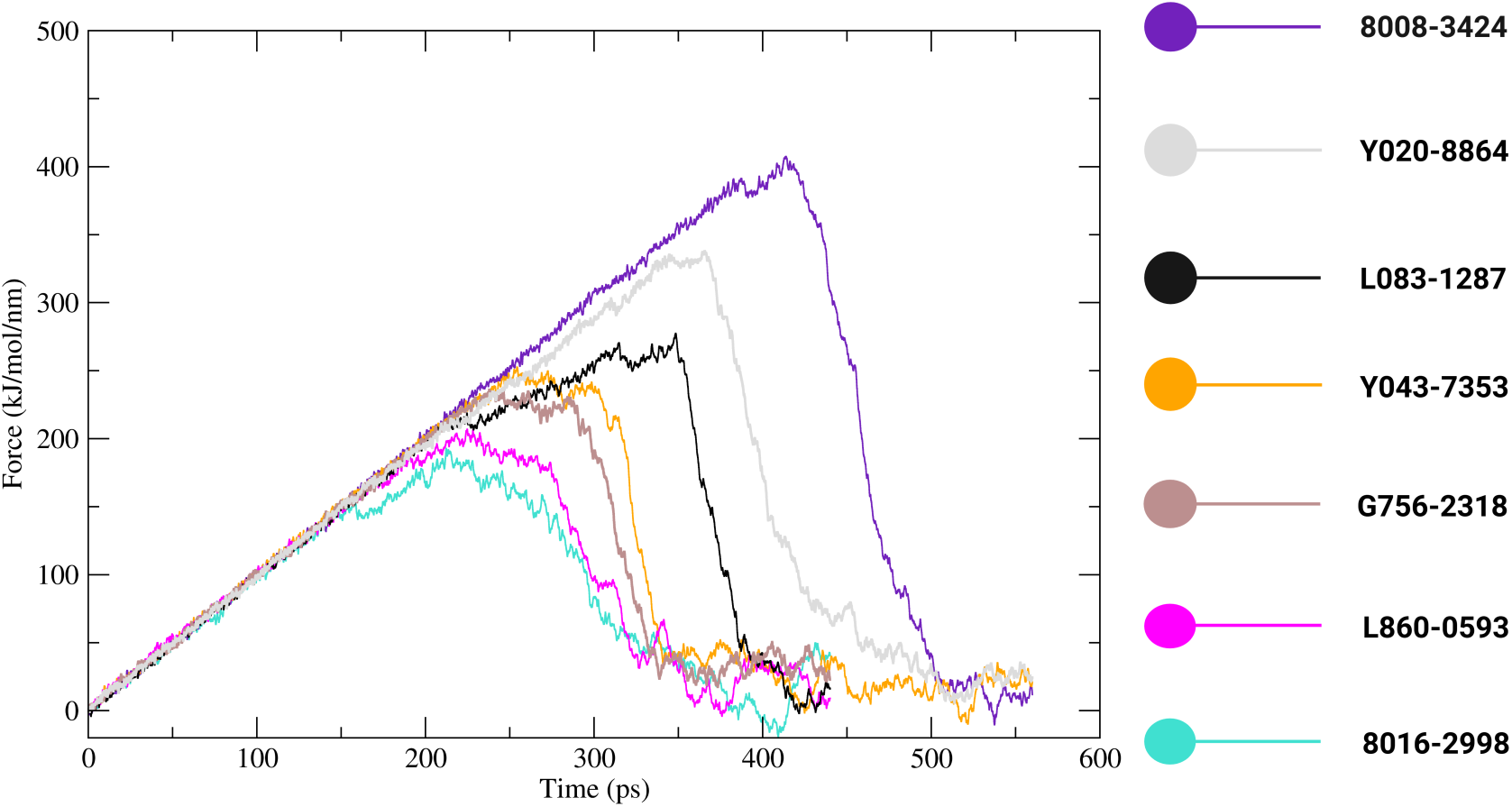
Results of sMD simulations performed on the seven molecules 8008-3424, Y020-8864, L083-1287, Y043-7353, G756-2318, L860-0593, and 8016-2998 that exhibited the lowest binding scores, as determined from a 50 ns MD simulation. The molecules are color-coded for clear distinction. On the plot, x-axis reveals time (ps) and y-axis highlights pulling force by unit of the kJ·mol⁻¹·nm⁻¹.

### 3.4 Determining the effects of c-MET inhibitors on cell survival *in vitro* and *in vivo*

c-MET, as the sole receptor for the HGF, plays pivotal roles in hepatocyte proliferation, survival, and migration, thereby contributing significantly to the development and progression of HCC. [61] Given its critical function in HCC, the impact of selected c-MET inhibitors was tested on HCC cell survival *in vitro* and on zebrafish development *in vivo*. Using a well-established HCC cell line, HuH7, the efficacy of six selected compounds (Y020-8864, L860-0593, L083-1287, 8016-2998, G756-2318, and 8008-3424) were evaluated. The half-maximal inhibitory concentration (IC_50_) values were determined via MTT assay at 24, 48, and 72 hours post-treatment. (Figure 5A) Mimicking an aggressive tumor microenvironment in HCC, cells were treated with HGF (10 ng/ml) after starvation. As the IC_50_ calculations suggested, Y020-8864 and G756-2318 displayed inconsistent inhibitory effects on cell survival. To elucidate the long-term effects of the six c-MET inhibitors, the colony formation capacities of HuH7 cells were evaluated under HGF stimulation. According to their IC_50_ value range, cells were treated at 0.1 μM, 1 μM, and 5 μM concentrations for 15 days. Even at 0.1 μM, L860-0593, L083-1287, 8016-2998, and 8008-3424 exhibited substantial suppressive effects compared to HGF controls. (Figure 5B) Notably, at 5 μM, a significant reduction in colony formation capacity was observed for almost all inhibitors.

**Figure 5.**
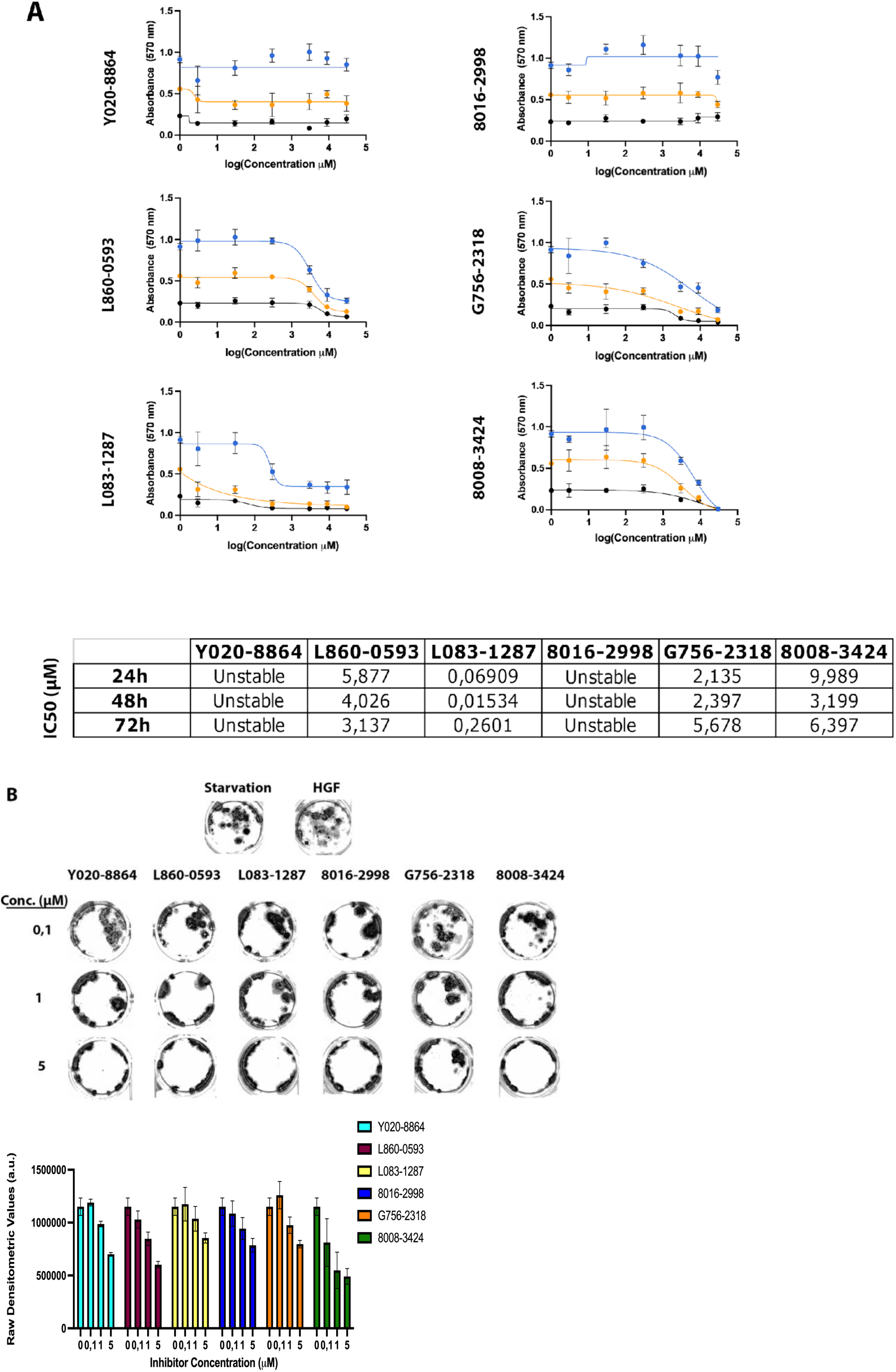
The Effect of c-Met Inhibitors on Cell Survival and Proliferation. **A**. MTT analysis of HuH7 cells when incubated with c-Met inhibitors for 24, 48 and 72 hours in concentrations ranging from 3, 30, 300, 3000, 9000 and 30000 nM. DMSO served as control and cells were treated with 10 ng/ml HGF. Absorbance at 570 nm was measured in Varioskan Plate Reader. The inhibitory concentration 50 (IC_50_) values were determined using GraphPad Prism v9.5.1. Inhibitor concentrations were log10-transformed, and dose-response inhibition was calculated using non-linear regression analysis. **B.** Colony formation assay determined the effect of c-Met inhibitors on cell survival in 15 days. Cells were treated with inhibitors at concentrations of 0.1 μM, 1 μM, and 5 μM, refreshed every 72 hours. DMSO served as the control, and cells were also treated with 10 ng/mL HGF. Colonies were stained with Giemsa and imaged using the BioRad ChemiDoc MP imaging system. The densitometric quantifications are shown on the right graph.

### 3.5. Mechanistic insights into how the inhibitors contributed to c-Met activation

To determine the effects of inhibitors on c-Met activation, phosphorylation at Tyr1234/35 was investigated. HuH7 cells were starved and subsequently treated with HGF (10 ng/mL) alongside the inhibitors that showed consistent effects on suppression of survival: L860-0593, L083-1287, G756-2318, and 8008-3424. The inhibitors were tested at 3 different concentrations (0.1 μM, 1 μM, and 5 μM) and cell lysates were analyzed by western blotting. (Figure 6A) All inhibitors showed a significant reduction in c-Met activation, except L860-0593. Especially, 5 μM concentration was found to be highly effective in suppressing c-Met activity. Among all, L083-1287 was identified as the most potent inhibitor considering its inhibitory capacity of HCC cell survival and c-Met activity.

**Figure 6.**
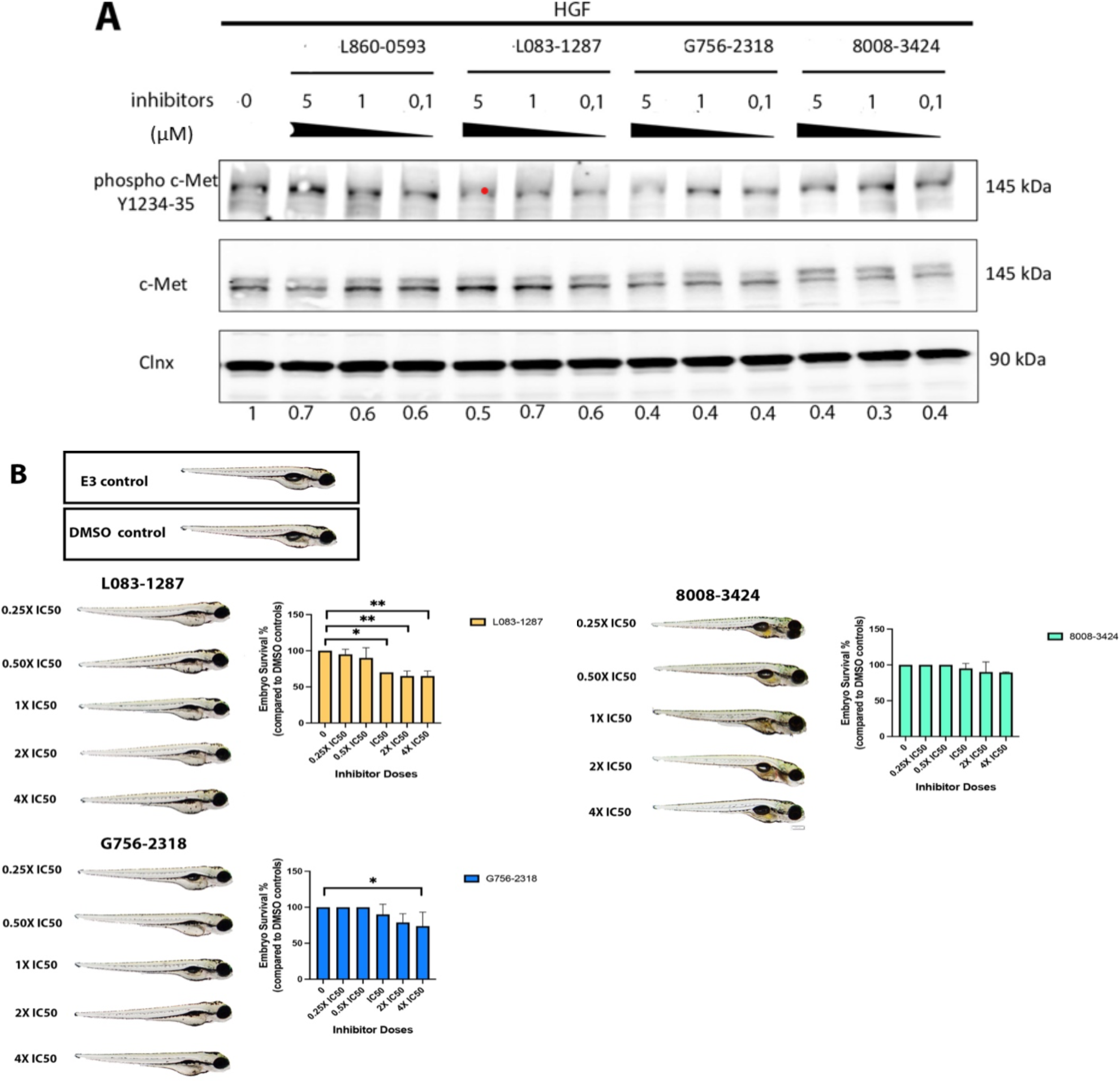
Effects of c-Met Inhibitors on c-Met Activation and Dose-Dependent Toxic Effects in Zebrafish. **A.** HuH7 cells were serum-starved and stimulated with HGF (10 ng/mL) for 2 hours, followed by treatment with c-Met inhibitors at concentrations of 0.1 μM, 1 μM, and 5 μM for 48 hours. DMSO was included as control for the HGF group. Cell lysates were analyzed via western blotting, by probing for c-Met (Y1234-35), total c-Met, and Calnexin. **B.** Representative images of 5-day post-fertilization (dpf) zebrafish larvae after 72 hours of treatment with c-Met inhibitors are shown. Zebrafish embryos were incubated with c-Met inhibitors in doses proportional to the IC_50_ values determined in MTT assay. Toxicity assays were performed for the inhibitors L083-1287 (65, 130, 260, 520, 1040 nM), G756-2318 (1419, 2839, 5678, 11356, 22712 nM), and 8008-3424 (1599, 3198, 6397, 12794, 25588 nM). E3 medium and E3 medium with DMSO served as controls. Zebrafish embryos were exposed to the treatment media starting at 2 dpf, with inhibitors being renewed every 24 hours. The experiment was concluded at 5 dpf, the embryos were fixed, analyzed under a stereomicroscope using a 4X objective, and imaged. Each experiment was repeated in 2 replicates (n=20/group). Scale bar = 200 μm.

### 3.6 Assessing c-MET Inhibitor Toxicity in Zebrafish Embryos

The zebrafish (Danio rerio) serves as a robust biological model due to its high genetic and functional homology with the human c-Met receptor, particularly within its highly conserved kinase and signaling domains. [62] Given the essential role of c-Met in normal development and growth, we further determined potential toxicity effects of selected inhibitors on zebrafish embryos. Embryos were grown in E3 medium containing DMSO or inhibitors at concentrations derived from their IC_50_ values (Figure 5A) at 2 dpf, and monitored until 5 dpf. The zebrafish were routinely checked, dead embryos were recorded and removed from the media. At the end of the experiment they were fixed, analyzed under the microscope and imaged. (Figure 6B) A survival analysis revealed that exposure to G756-2318 at 22.7 μM led to a reduction in survival rates. Although L083-1287 emerged as one of the most active candidates in the c-MET inhibition assays, its evaluation in zebrafish embryos revealed a concentration-dependent reduction in survival at 0.26, 0.52, and 1.04 μM. This was accompanied by developmental phenotypes, including cardiac and vascular alterations, abnormal body posture, and swim bladder defects, which were more apparent at higher inhibitor concentrations (Fig. 6B). These results indicate that L083-1287 retains strong pharmacological activity but requires further lead optimization to improve its developmental tolerability and therapeutic window. Thus, rather than excluding L083-1287 from further consideration, the zebrafish findings define a clear optimization path: maintaining potent c-MET pathway inhibition while reducing concentration-associated developmental liabilities.

In summary, these findings emphasize the importance of balancing efficacy and safety in the development of c-MET inhibitors. Among the tested hit compounds, L083-1287 emerged as the most potent, necessitating further investigations to optimize its clinical application while minimizing potential developmental toxicity.

### 3.7 L083-1287 and 8008-3424 directly binds to c-Met

To validate direct target engagement, we used surface plasmon resonance (SPR; Biacore), a sensitive, label-free biophysical method that enables real-time quantification of molecular binding events. The ECD of recombinant human c-Met was immobilized on a CM5 sensor chip and used as the analyte-binding surface. The recombinant protein was expressed in HEK293 cells. Therefore, the protein purified by affinity chromatography is expected to have proper posttranslational modifications and proper tertiary structure. L083-1287 and 8008-3424 compounds were subsequently injected over the c-Met-ECD coated chip surface and binding interactions were monitored in real time. (Figure 7) We observed a dose-dependent reproducible interaction between the recombinant c-Met-ECD and L083-1287 and 8008-3424. The K_D_ values were calculated to be 27.1 μM and 33.1 μM for L083-1287 and 8008-3424, respectively. Collectively, these data provide direct experimental evidence that both small molecules bind the extracellular domain of c-Met, supporting the proposed mechanism of action and confirming c-Met-ECD as a bona fide molecular target.

**Figure 7.**
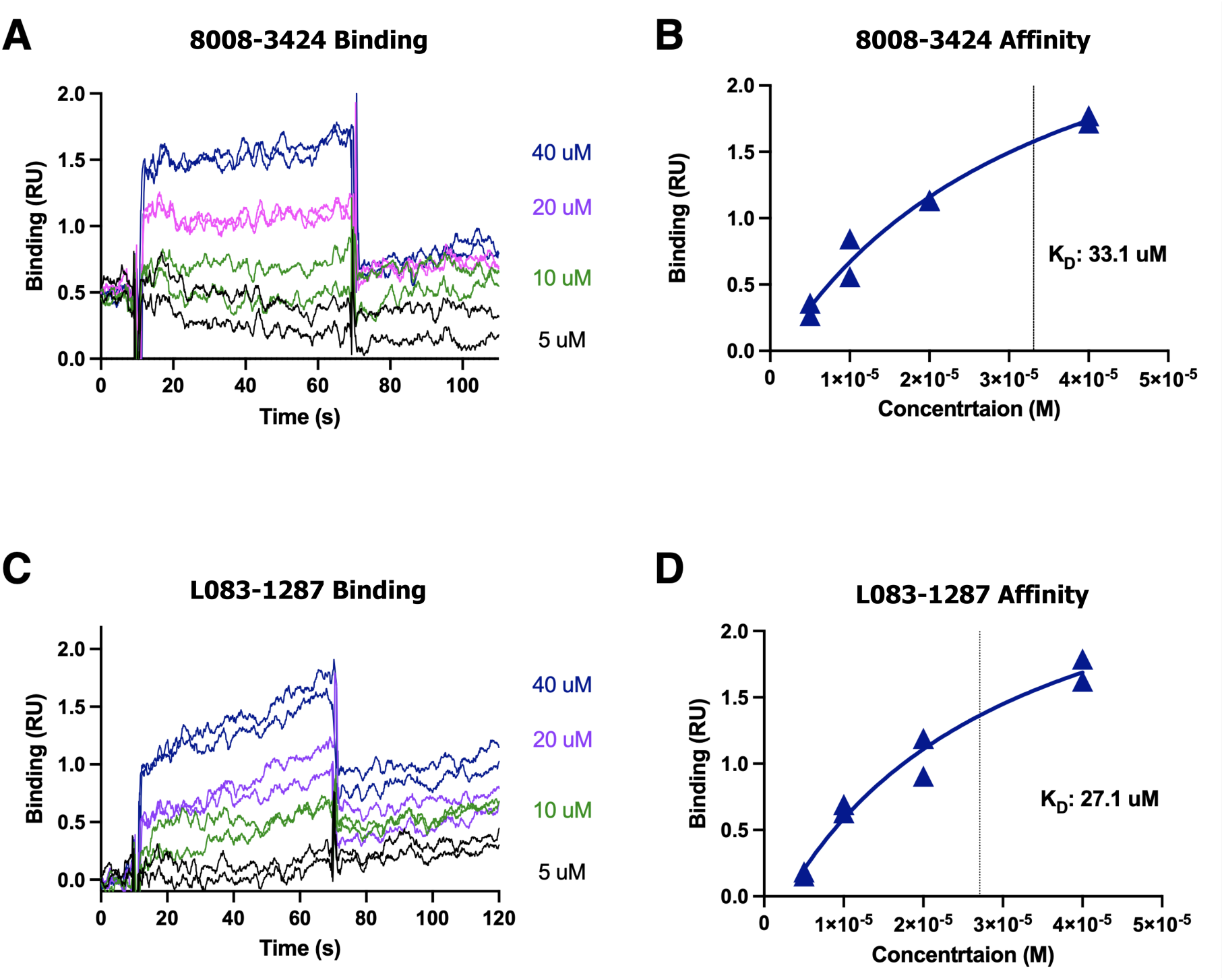
Direct binding of 8008-3424 and L083-1287 to recombinant c-Met protein. ECD of human c-Met (aa25-932) protein was immobilized on a CM5. Small molecules were injected over the protein coated surface at 4 different doses in duplicates and the binding was measured by SPR in a Biacore T-200 instrument. Binding curves for 60 second injections are presented in panels A and C. Binding values 5 sec before the injection end in panels A and C, were plotted against compound concentrations in panel B and D, respectively, to calculate the steady state binding affinity (K_D_).

### 3.8 Identification of Analogs of Determined Lead Compounds: First Lead Compound Optimization Cycle

Encouraged by the initial activity results of 8008-3424 and L083-1287 against c-MET, we next undertook a first-round lead-optimization campaign to explore whether structurally related analogs could deliver improved binding characteristics and broader drug-like potential. Although both compounds emerged as promising starting points, our goal was to expand the accessible chemical space around these scaffolds and identify analogs that might support subsequent optimization of both potency and downstream pharmaceutical properties. Thus, we performed large-scale similarity searches with SwissSimilarity against the ZINC drug-like library, encompassing approximately 9 million commercially available compounds. For each parent ligand, approximately 400 analog candidates were retrieved. To preserve chemical diversity and avoid biasing the search toward narrowly defined chemotypes, we did not apply restrictive pre-docking filters. Instead, all candidates were advanced into a hierarchical computational pipeline comprising Glide/XP docking, MM/GBSA rescoring, and replicate MD-based validation. The top 60 analogs, selected based on favorable MM/GBSA energies, were each subjected to five independent 10 ns MD simulations to evaluate the reproducibility and stability of protein–ligand interactions. Trajectory-based free-energy analyses were then used to benchmark each analog against the parent compounds. Analogs that consistently exceeded 8008-3424 and L083-1287 in terms of predicted binding energetics and conformational stability were advanced to longer (200 ns) simulations for deeper characterization. This iterative procedure enabled prioritization of compounds that not only bound favorably in static models but also maintained stable engagement under dynamic conditions. More broadly, this first optimization cycle was designed to move beyond simple hit identification and toward the establishment of a more actionable lead series, recognizing that effective c-MET inhibitors will ultimately require a coordinated optimization of potency, binding stability, selectivity, and safety-related properties. In this extended refinement stage, selected molecules underwent triplicate 200 ns MD simulations, followed by a second round of MM/GBSA free energy calculations. These long trajectories afforded deeper insight into conformational adaptability and the persistence of key molecular interactions within the c-MET binding pocket. The results of this multi-tiered screening pipeline are summarized in Table 2, which highlights the most promising analogs. Several candidates demonstrated superior binding energetics and sustained structural stability relative to the reference compounds, underscoring their potential as next-generation c-MET inhibitors. These findings validate the robustness of our integrated workflow-combining similarity-driven searches, high-precision docking, and MD-based free energy evaluation, as an effective strategy for the discovery of therapeutic leads.

**Table 2.**
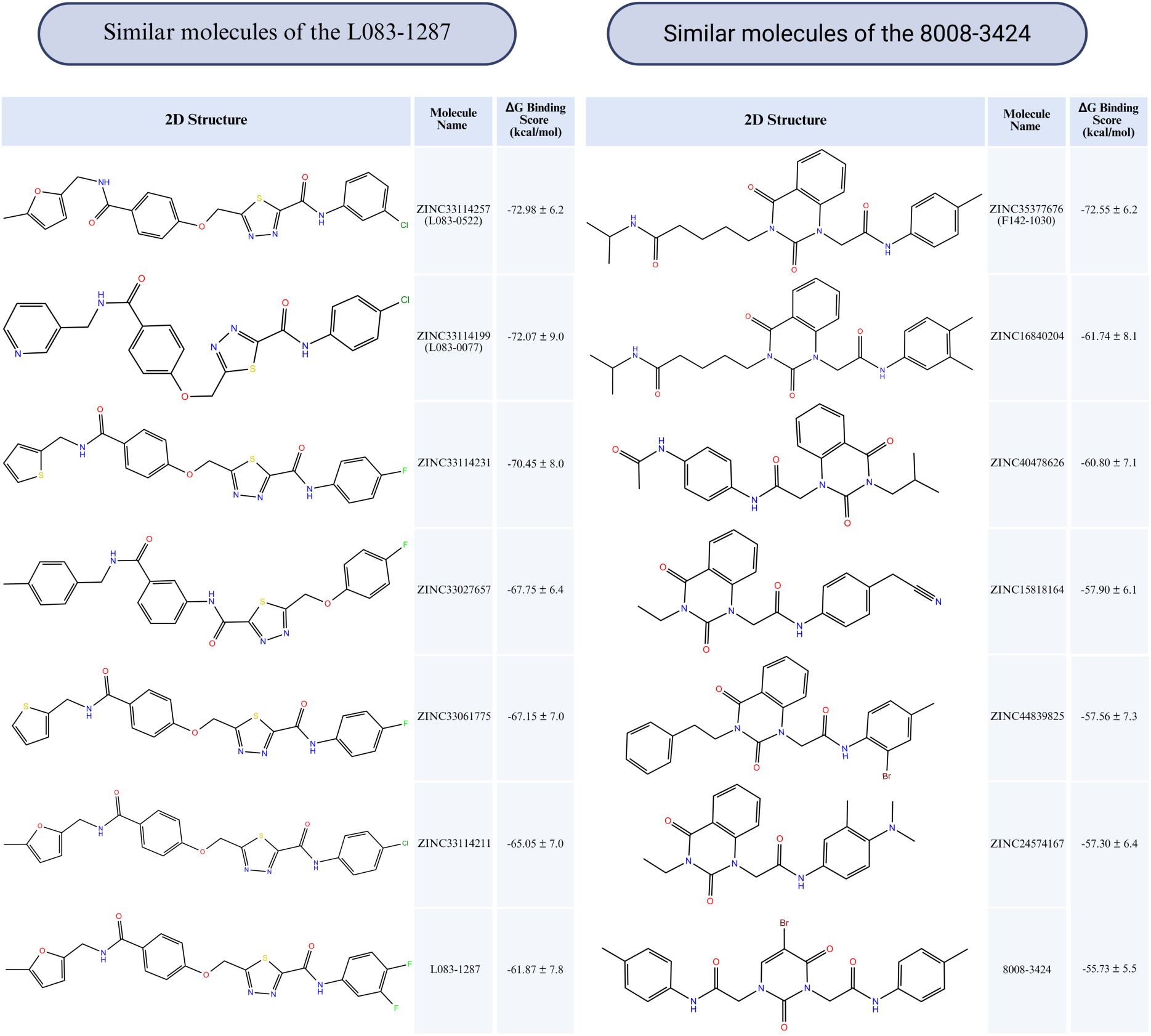
Post-experimental validation revealed that the molecules L083-1287 (targeting c-MET) and 8008-3424 (targeting the c-MET/HGF interface) effectively inhibit the c-MET/HGF interaction via distinct mechanisms of action. These compounds exhibited both low toxicity profiles and strong binding affinities in preliminary *in vitro* and *in silico* assays. To expand the chemical landscape and discover structurally related analogs with potentially enhanced pharmacological properties, a comprehensive similarity analysis was conducted using the ZINC drug-like library.

Similar molecules of the L083-1287
| 2D Structure | Molecule Name | $\Delta G$ Binding Score (kcal/mol) |
| --- | --- | --- |
| | ZINC33114257 (L083-0522) | -72.98 ± 6.2 |
| | ZINC33114199 (L083-0077) | -72.07 ± 9.0 |
| | ZINC33114231 | -70.45 ± 8.0 |
| | ZINC33027657 | -67.75 ± 6.4 |
| | ZINC33061775 | -67.15 ± 7.0 |
| | ZINC33114211 | -65.05 ± 7.0 |
| | L083-1287 | -61.87 ± 7.8 |

Similar molecules of the 8008-3424
| 2D Structure | Molecule Name | $\Delta G$ Binding Score (kcal/mol) |
| --- | --- | --- |
| | ZINC35377676 (F142-1030) | -72.55 ± 6.2 |
| | ZINC16840204 | -61.74 ± 8.1 |
| | ZINC40478626 | -60.80 ± 7.1 |
| | ZINC15818164 | -57.90 ± 6.1 |
| | ZINC44839825 | -57.56 ± 7.3 |
| | ZINC24574167 | -57.30 ± 6.4 |
| | 8008-3424 | -55.73 ± 5.5 |

Furthermore, sMD simulations were performed for L083-1287 and its six analogs ZINC33114211, ZINC33114231, ZINC33114257 (L083-0522), ZINC33027657, ZINC33061775, and ZINC33114199 (L083-0077), which were selected based on their favorable predicted binding affinities and close structural resemblance to the parent compounds. (Figures S10) Similarly sMD simulations were also conducted for 8008-3424 and its 9 analogs. (Figure S11) These simulations were intended to evaluate the mechanical stability of ligand binding to c-MET under externally applied force, thereby providing an additional level of discrimination beyond conventional docking and equilibrium MD analyses. In this setting, sMD makes it possible to assess how effectively each ligand remains bound within the active site when the complex is mechanically perturbed, offering insight into the relative robustness of protein–ligand interactions under stress. Analysis of the sMD showed clear differences in binding behavior among the six analogs. Notably, ZINC33114211, ZINC33114231, and ZINC33114257 exhibited substantially lower binding energies than the reference ligand, L083-1287, indicating stronger and more stable interactions with the c-MET binding pocket. These findings suggest that these three analogs may bind the receptor more effectively than the parent compounds and therefore warrant further consideration as promising candidates for c-MET inhibitor optimization. In contrast, ZINC33027657, ZINC33061775, and ZINC33114199 displayed binding energies generally comparable to those of L083-1287, suggesting similar overall interaction strength but no clear improvement in binding stability or resistance to applied force. Overall, these results highlight a subset of L083-1287-derived analogs with superior performance in sMD simulations and support their prioritization for longer-timescale simulations and experimental validation.

Similarly, sMD simulations were conducted for 8008-3424 and nine structurally related analogs selected from the c-MET/HGF-complex 200-ns MD screening workflow. (Figure S11) These simulations were designed to assess the mechanical stability of ligand binding under externally applied force, thereby providing an additional discriminatory layer beyond docking and equilibrium MD analyses. In this context, the sMD force–time profiles revealed clear differences in dissociation behavior across the series, indicating that closely related analogs do not exhibit equivalent resistance to forced unbinding. The reference compound, 8008-3424, displayed an earlier rupture event and a lower maximum pulling force than several of its analogs, consistent with only moderate mechanical stability within the c-MET/HGF-associated binding environment. Notably, ZINC15190707, ZINC44839825, and ZINC40478626 showed the most favorable sMD profiles, reaching the highest peak pulling forces and maintaining force-bearing trajectories for the longest simulation times before dissociation. These features indicate a markedly greater resistance to forced unbinding than the parent compound and therefore suggest stronger and more persistent ligand engagement under perturbation. A second group, comprising ZINC24574167 and ZINC15818164, also outperformed 8008-3424, as evidenced by their higher rupture forces and later detachment times, although their performance remained below that of the top three analogs. In addition, ZINC33027657, ZINC16840204, and ZINC35377676 demonstrated modest improvement over the reference structure, showing somewhat elevated force maxima and prolonged retention relative to 8008-3424, but without the pronounced mechanical resilience observed for the best-performing compounds. By contrast, ZINC33330913 exhibited the weakest profile in the series, with an earlier decline in force and premature dissociation, indicating lower stability than the parent scaffold.

Taken together, these results identify discrete subsets of analogs from both chemotypes with superior performance in sMD simulations. Within the L083-1287 series, ZINC33114211, ZINC33114231, and ZINC33114257 emerged as the most promising candidates, whereas within the 8008-3424 series, ZINC15190707, ZINC44839825, and ZINC40478626 showed the clearest improvement over the parent scaffold, with ZINC24574167 and ZINC15818164 representing additional favorable analogs. Overall, the sMD data indicate that mechanical stability under nonequilibrium force provides an informative secondary filter for ligand prioritization, complementing equilibrium binding analyses and highlighting compounds that may be better suited for longer-timescale simulations, rigorous free-energy evaluation, and experimental validation.

### 3.9 *In vitro* characterization of optimized lead compounds

To assess whether lead optimization improved the anti-survival activity of the selected c-MET-targeting chemotypes, optimized compounds (L083-0522, F142-0205, L083-0077) together with parent compound (L083-1287) were evaluated in HuH7 cells using the same MTT-based dose–response assay described above. Cell viability was measured at 24, 48 and 72 h, and IC_50_ values were calculated by non-linear regression of log-transformed compound concentrations. (Figure 8) Among the tested compounds, parent compound L083-1287 showed the strongest activity, with consistently submicromolar IC_50_ values at all time points (0.246 µM, 0.146 µM and 0.222 µM at 24, 48 and 72 h, respectively), indicating both rapid and sustained suppression of HuH7 cell survival. By contrast, L083-0522 displayed intermediate but stable activity, with IC_50_ values remaining close to 8 µM throughout treatment (8.042 µM, 8.325 µM and 8.504 µM). L083-0077 was also found effective overall and showed reduced potency over time, with the IC_50_ increasing from 8.849 µM at 24 h to 19.944 µM and 18.671 µM at 48 and 72 h, respectively. Evaluation of these compounds in non-MET-expressing SNU-398 cells (CMVGFP) and their MET-overexpressing counterpart (METGFP) was performed to assess the impact of MET overexpression on drug response. The parent compound L083-1287 exhibited the strongest overall activity; however, its potency was reduced in METGFP cells, as reflected by higher IC50 values (0.268 µM and 0.108 µM at 48 and 72 h, respectively), compared to the CMVGFP control (0.186 µM and 0.061 µM at 48 and 72 h, respectively) (Figure 8B). In contrast, L083-0522 demonstrated a different trend, with METGFP cells showing lower IC50 values (9.349 µM, 9.442 µM, and 10.202 µM at 24, 48, and 72 h, respectively), particularly at later time points, compared to CMVGFP control (9.631 µM, 13.182 µM, and 23.115 µM at 24, 48, and 72 h, respectively) (Figure 8C).

**Figure 8.**
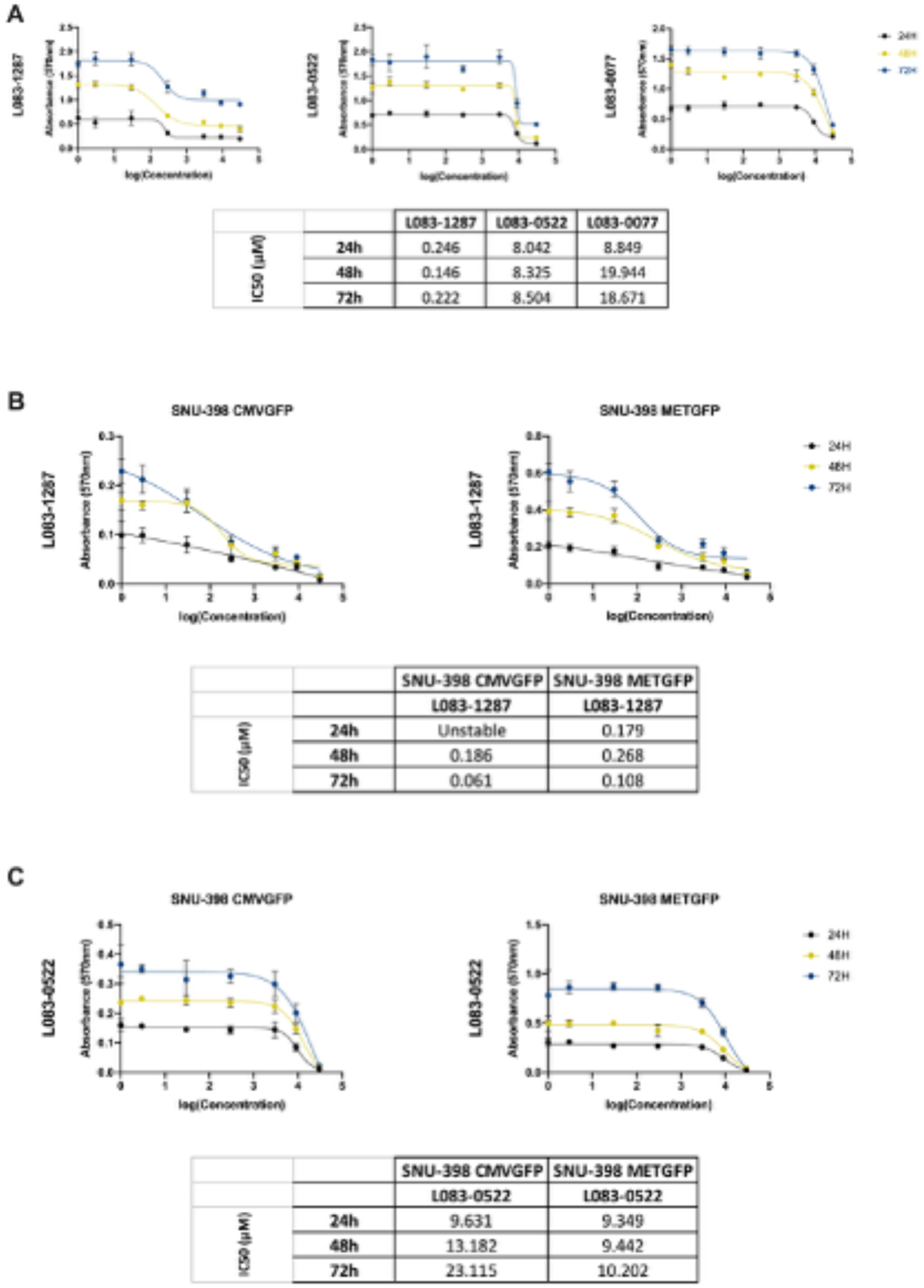
The Effect of Optimized c-Met Inhibitors on Cell Survival in Different Cell Lines. MTT analysis of HuH7 cells when incubated with parent (L083-1287) and optimized c-Met inhibitors (L083-0522, L083-0077) for 24, 48 and 72 hours **(A)**, and SNU-398 CMVGFP and METGFP cells when incubated with parent **(B)** and optimized c-Met inhibitor **(C)** in concentrations ranging from 3, 30, 300, 3000, 9000, and 30000 nM. DMSO served as control.

Absorbance at 570 nm was measured in Varioskan Plate Reader. The inhibitory concentration 50 (IC_50_) values were determined using GraphPad Prism v9.5.1. Inhibitor concentrations were log10-transformed, and dose-response inhibition was calculated using non-linear regression analysis.

The activity of F142-0205 was further examined under both HGF(-) and HGF(+) conditions to determine whether ligand-dependent c-MET activation influenced drug response (Figure 9). In the absence of HGF, F142-0205 exhibited IC_50_ values of 8.128 µM, 9.161 µM and 10.07 µM at 24, 48 and 72 h, whereas under HGF stimulation these values increased to 18.19 µM, 14.43 µM and 15 µM. This rightward shift in the dose–response curves indicates that HGF-mediated pathway activation partially attenuates the inhibitory effect of F142-0205.

**Figure 9.**
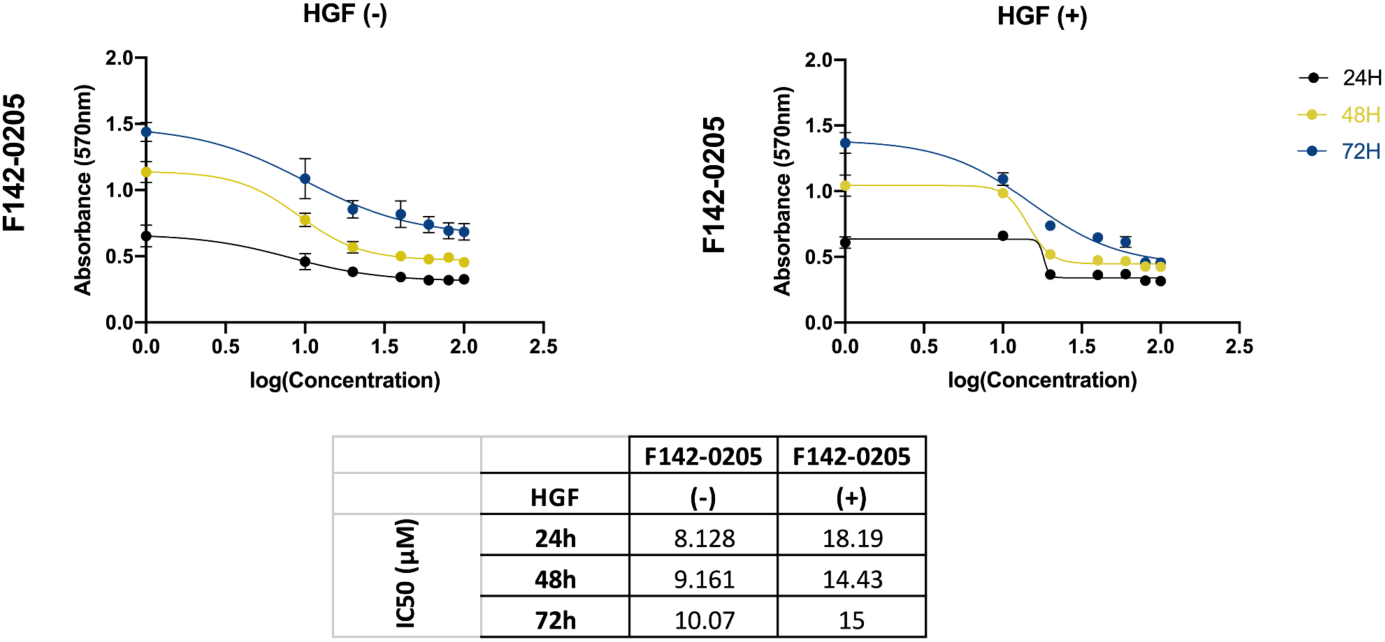
The Effect of Optimized c-Met Inhibitor on Cell Survival under HGF stimulation. MTT analysis of HuH7 cells when incubated with optimized c-Met inhibitor F142-0205 for 24, 48 and 72 hours in concentrations ranging from 3, 30, 300, 3000, 9000, and 30000 nM. DMSO served as control and cells were treated with 10 ng/ml HGF. Absorbance at 570 nm was measured in Varioskan Plate Reader. The IC_50_ values were determined using GraphPad Prism v9.5.1. Inhibitor concentrations were log10-transformed, and dose-response inhibition was calculated using non-linear regression analysis.

Taken together, these findings show that lead optimization did not uniformly enhance the anti-survival activity of this c-MET-targeting scaffold in HuH7 or SNU-398 cells. Although the optimized derivatives retained measurable inhibitory effects, none surpassed the parent compound L083-1287, which remained the most potent molecule across all time points and consistently displayed submicromolar activity in both non-MET-expressing and MET-overexpressing cells. L083-0522 and L083-0077 showed moderate but stable profiles. In parallel, the reduced activity of F142-0205 under HGF stimulation indicates that ligand-dependent activation of the HGF/c-MET axis can partially blunt compound efficacy. Overall, these results identify L083-1287 as the most promising candidate within this series and suggest that future optimization efforts should prioritize preservation of its potency while improving robustness under HGF-driven conditions.

### 3.10 L083-0077 directly binds to c-Met-ECD

To determine the binding affinity of optimized compounds for c-Met-ECD, we performed SPR analysis. In this assay, the surface of a Biacore sensor chip was coated with c-Met-ECD protein, and optimized compounds (i.e., L083-0077) were injected as the analyte at four different concentrations. (Figure 10) The binding profile of L083-077 differed from those observed for parent compounds L083-1287 and 8008-3424. For L083-1287 and 8008-3424, both the association and dissociation phases were relatively rapid (Figure 10), which limited the analysis to calculation of the equilibrium dissociation constant (K_D_) only. In contrast, L083-077 exhibited slower association and slower dissociation kinetics, allowing calculation of not only the binding affinity (K_D_), but also the association rate constant (k_a_) and dissociation rate constant (k_d_). The K_D_ value for the interaction between L083-077 and c-Met-ECD was determined to be 17.4 µM. These findings indicate that L083-077 binds directly to c-Met-ECD and displays a slightly improved binding affinity compared with the previously tested compounds.

**Figure 10.**
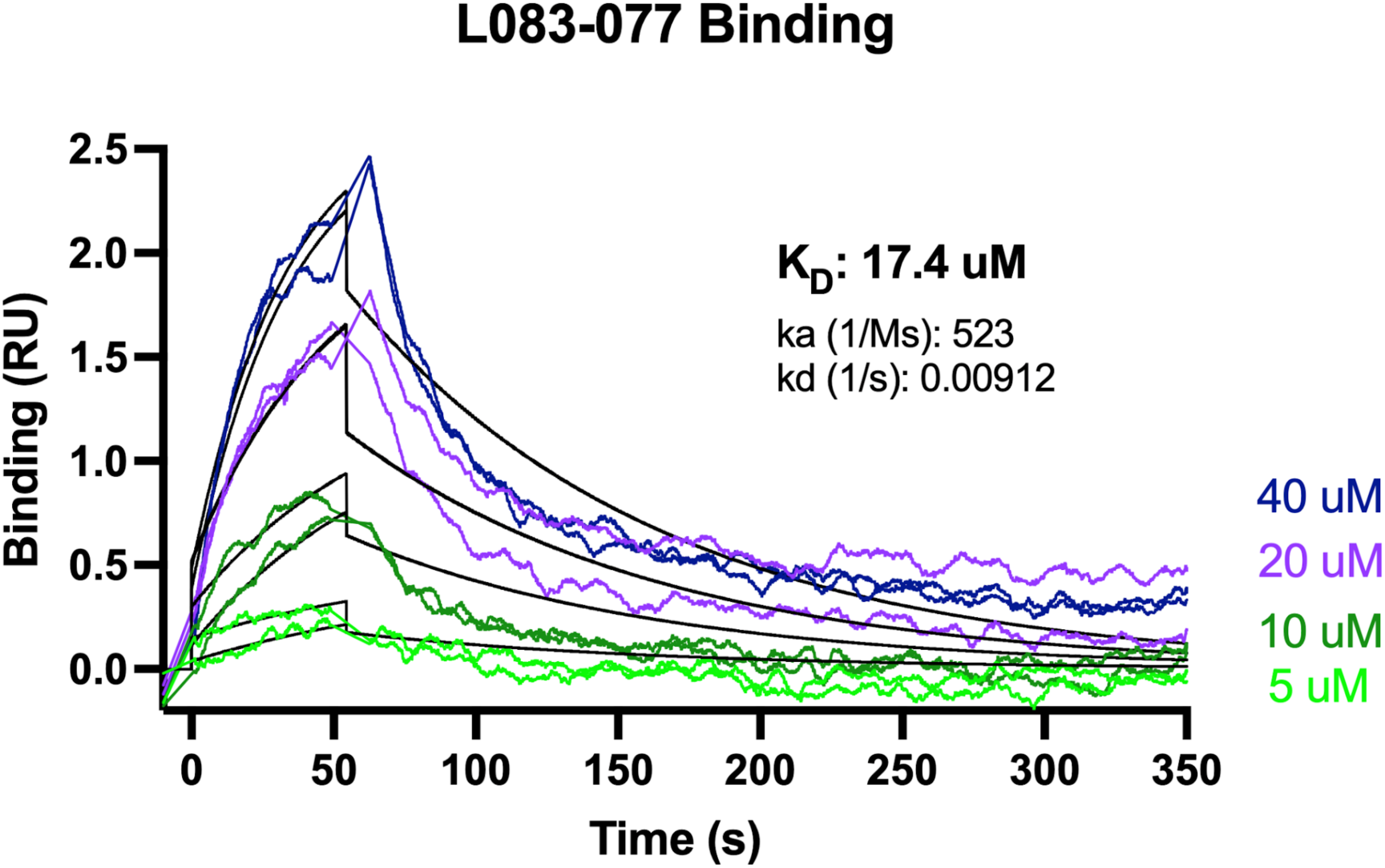
Direct binding of L083-077 to recombinant c-Met protein. ECD of human c-Met (aa25-932) protein was immobilized on a CM5 chip. L083-077 was injected overt the protein coated surface at 4 different doses in duplicates and the binding was measured by SPR in a Biacore T-200 instrument. The compound was injected for 60 seconds, followed by data collection for 300 seconds for the dissociation phase. Colored lines show the actual binding curves. Black lines indicate the 1:1 binding model that was used to calculate binding kinetics values.

### 3.11 Structural Interaction Dynamics of L083-0077

The structural interaction dynamics (SID) analysis of the selected lead compound (L083-0077) indicated relatively stable binding mode accompanied by progressive conformational adaptation over the course of the 200-ns MD simulations. (Figure 11) Throughout the trajectory, the protein backbone remained comparatively well restrained, with RMSD values largely confined to 2–6 Å, consistent with preservation of the global fold. By contrast, the ligand displayed a broader positional deviation, with RMSD values fluctuating between 5 and 9 Å. Importantly, these ligand fluctuations are unlikely to reflect simple thermal noise or nonspecific drift within the binding region. Rather, they are consistent with a directed conformational transition from a more extended initial arrangement toward a partially compacted or folded state, in which the ligand-bound complex is associated with a progressive inward movement of the IPT2 domain toward the central interfacial region. This compaction-like behavior suggests activation of a local structural rearrangement process analogous to that described in related receptor systems, where domain reorientation contributes to stabilization of metastable binding conformations.

**Figure 11.**
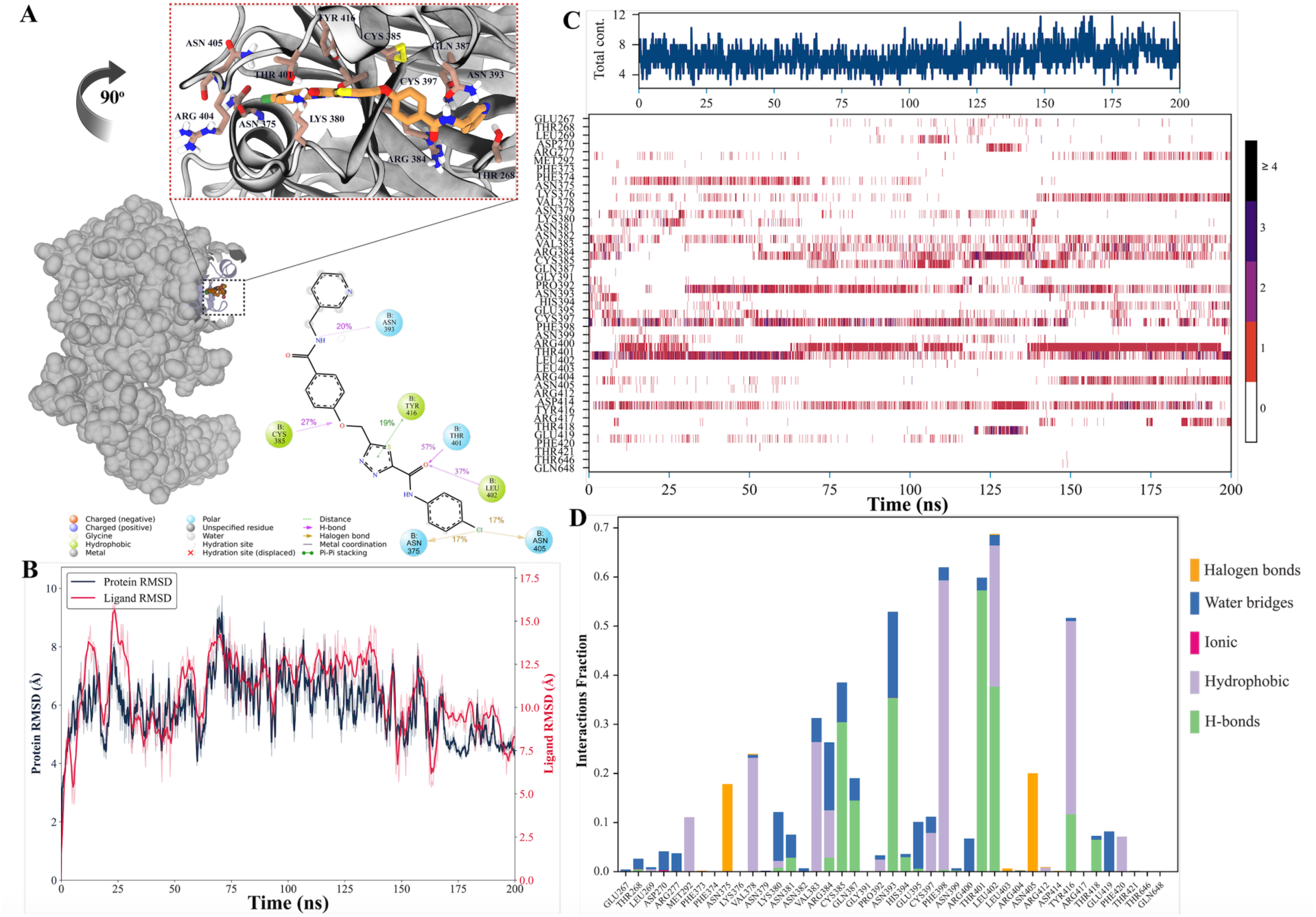
(A) Three-dimensional and two-dimensional interaction maps describing the binding mode of the selected hit compound (L083-0077) in the c-MET active site. (B) Time-resolved residue contact profile for the c-MET-hit ligand (L083-0077) complex. Simulation time is reported on the x-axis (ns), and c-MET residue IDs are shown on the y-axis; horizontal bars quantify the number of contacts formed between each residue and the ligand. (C) RMSD traces extracted from the MD trajectory. Protein RMSD was calculated over backbone atoms, whereas ligand RMSD was computed over all ligand atoms. Time (ns) is plotted on the x-axis and RMSD values are reported in Å. (D) Residue-level interaction occupancy of L083-0077 throughout the simulation. Residues are listed on the x-axis and the y-axis reports interaction fraction, defined as the proportion of MD frames in which a given residue maintains contact with the ligand.

Temporal interaction profiling further showed that the ligand did not remain statically anchored to a single contact network but instead engaged in a dynamic and continuously reorganizing set of interactions. Across the simulation, transient and discontinuous contacts were observed with Asn405, Tyr404, Cys397, Asn393, Phe398, Lys380, and Asp275. Notably, the canonical hotspot residues Phe398 and Cys397 exhibited only weak and intermittent interactions, indicating that the compound did not achieve sustained anchoring within the principal hotspot region of the pocket. This observation is consistent with a binding mode of intermediate affinity, in which cavity residence is maintained, but hotspot capture remains incomplete. In contrast, more persistent hydrogen-bonding interactions were detected with Arg404 and Asn405, suggesting that polar contacts near the upper portion of the binding environment contributed more substantially to ligand retention. In addition, Asn275, positioned closer to the lower IPT2-proximal surface, formed short-lived yet repeatedly recurring hydrogen bonds. Although individually transient, these interactions likely provided cumulative local stabilization during the domain-directed compaction process and may have helped accommodate the evolving geometry of the ligand-bound state.

The 2D interaction map and contact histogram (Figure 11D) further supported this interpretation by demonstrating that hydrogen-bonding interactions involving Asp275 and Asn405 occurred intermittently rather than continuously, while π–π stacking interaction with Phe398 was observed only sporadically. The intermittent character of these interactions indicates that the binding mode was not rigid, but instead remained structurally plastic, with the ligand adapting to ongoing rearrangements in the surrounding protein environment. The temporal organization of the contact network is therefore highly informative: As the IPT2 domain gradually approached the ligand-occupied region, earlier contacts were weakened or replaced by alternative local interactions, giving rise to periodic changes in ligand orientation and corresponding RMSD oscillations. These fluctuations should therefore not be interpreted as evidence of global destabilization or incipient ligand dissociation, but rather as a signature of conformational accommodation within a dynamically remodeling binding interface.

Taken together, these findings indicate that the compound remains resident within the binding cavity over the full simulation timescale but does so through a binding regime characterized by incomplete hotspot engagement and conformational adaptability. The observed interaction pattern suggests that ligand retention is supported less by persistent occupation of the canonical aromatic hotspot and more by a shifting ensemble of polar and transient contacts that accompany the progressive approach of the IPT2 domain. Accordingly, the RMSD oscillations observed in the simulation are best understood as reflecting a biologically meaningful, fold-associated rearrangement, specifically, the movement of the IPT2 domain toward the ligand-binding interface, rather than a simple loss of complex stability. This behavior may represent an early transition toward a more compact, activation-like receptor conformation, while also highlighting the need for further ligand optimization to improve sustained engagement of key hotspot residues and thereby reinforce binding stability.

### 3.12 Second Lead compound optimization Cycle

Based on the integrated experimental and computational evaluation, the L083-1287 analogue L083-0077 was prioritized as one of the most promising candidates for the next lead-optimization cycle. This selection was supported by SPR analysis confirming direct target engagement, *in vitro* activity with micromolar IC₅₀ values across 24–72 h treatment windows, and comprehensive MD simulations indicating stable binding behavior, persistent key interactions, and a favorable dynamic profile within the target binding site. Together, these orthogonal data suggest that L083-0077 retains the core pharmacological features of the parent scaffold while offering a rational starting point for further structural refinement. Therefore, L083-0077 was selected to guide the second optimization stage, with the aim of improving potency, binding stability, and the overall therapeutic window while maintaining target-directed activity. Accordingly, similarity-based virtual screening was performed around the L083-0077 scaffold, yielding 303 analogues, which underwent Glide SP docking followed by Prime MM/GBSA rescoring. Of these, 34 molecules showed equal or higher predicted affinity compared to L083-0077 and were advanced to short 25 ns MD simulations (5 replicas), coupled with MM/GBSA filtering. Eighteen compounds met the affinity threshold and proceeded to 3 replicas 200 ns MD simulations. (Table 3)

**Table 3.**
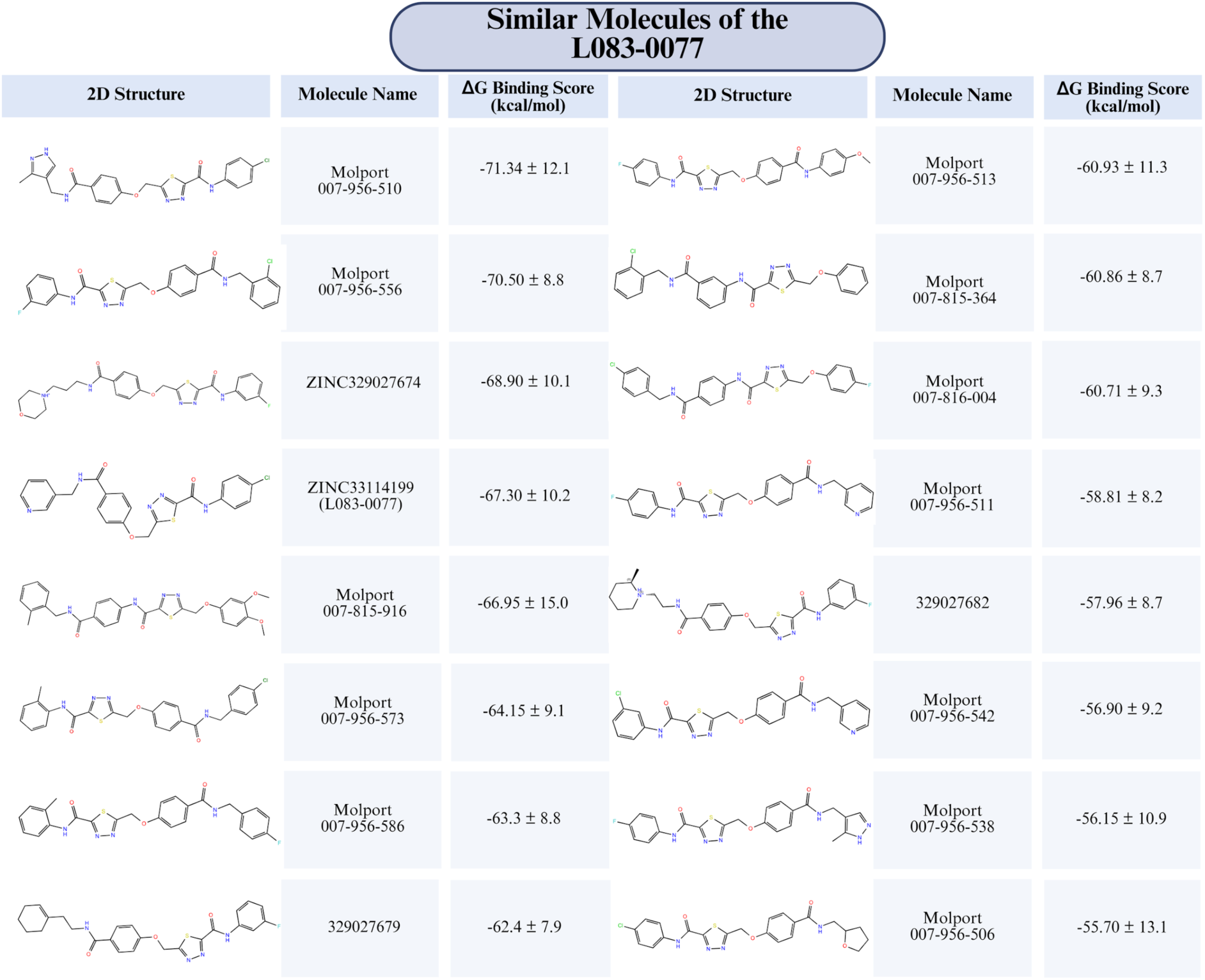
MM/GBSA binding free energies (ΔG, kcal/mol) obtained from 200 ns MD simulations of candidate molecules selected in the second lead optimization cycle. Lead compound L083-0077 was included as a parent compound to benchmark binding affinity. Compounds exhibiting lower (more negative) ΔG values than L083-0077 were considered to have enhanced binding performance and were considered for further evaluation.

| Similar Molecules of the L083-0077 |  |  |  |  |  |
| --- | --- | --- | --- | --- | --- |
| 2D Structure | Molecule Name | $\Delta G$ Binding Score (kcal/mol) | 2D Structure | Molecule Name | $\Delta G$ Binding Score (kcal/mol) |
| | Molport 007-956-510 | $-71.34 \pm 12.1$ | | Molport 007-956-513 | $-60.93 \pm 11.3$ |
| | Molport 007-956-556 | $-70.50 \pm 8.8$ | | Molport 007-815-364 | $-60.86 \pm 8.7$ |
| | ZINC329027674 | $-68.90 \pm 10.1$ | | Molport 007-816-004 | $-60.71 \pm 9.3$ |
| | ZINC33114199 (L083-0077) | $-67.30 \pm 10.2$ | | Molport 007-956-511 | $-58.81 \pm 8.2$ |
| | Molport 007-815-916 | $-66.95 \pm 15.0$ | | 329027682 | $-57.96 \pm 8.7$ |
| | Molport 007-956-573 | $-64.15 \pm 9.1$ | | Molport 007-956-542 | $-56.90 \pm 9.2$ |
| | Molport 007-956-586 | $-63.3 \pm 8.8$ | | Molport 007-956-538 | $-56.15 \pm 10.9$ |
| | 329027679 | $-62.4 \pm 7.9$ | | Molport 007-956-506 | $-55.70 \pm 13.1$ |

Pull MD analysis of the three top-scoring candidates revealed that Molport-007-956-556 and Molport-007-956-510 displayed rupture-force profiles comparable to the reference L083-0077, whereas L083-0725 exhibited a markedly weaker mechanical stability, detaching at approximately 150 kJ/mol. (Figure S12)

### 3.13 Fragment-Based Binding Free Energy Decomposition of Lead Compounds

To gain structural insight into the molecular determinants of ligand binding, we next performed fragment-resolved MM/GBSA analysis on the lead compounds. This approach enables decomposition of the overall binding free energy into contributions from individual regions of the ligand, thereby identifying which structural elements contribute most strongly to target engagement. By mapping the energetic contribution of each fragment, this analysis provides a useful framework for interpreting ligand binding mode and for guiding subsequent structure-based optimization. Fragment-based MM/GBSA analyses of L083-1287 and its analog L083-0077 showed that the overall binding free energy was not distributed uniformly across the molecule, but was instead dominated by a limited number of structurally important regions.

(Figure 12) For L083-1287, the central phenyl linker made the largest contribution to complex stabilization (−19.99 kcal/mol, normalized score by number of non-hydrogen atoms: -2.49 kcal/mol), identifying this fragment as a major determinant of ligand binding. (Figure 12, top) Likewise, the para-difluoro-substituted phenyl ring contributed strongly (−17.56 kcal/mol, normalized score: -2.49 kcal/mol), indicating that this aromatic region also plays a critical role in target recognition. Together, these pronounced contributions suggest that the binding mode of L083-1287 is primarily supported by favorable hydrophobic packing and aromatic π-mediated interactions within the receptor pocket. In contrast, the peripheral regions of the molecule contributed less strongly to the calculated binding energy. The 1,2,4-thiadiazole ring (−13.72 kcal/mol, normalized score: -2.74 kcal/mol) and acetamide linkage (−10.36 kcal/mol, normalized score: -3.45 kcal/mol) provided intermediate stabilization, whereas the terminal lactone moiety (−9.20 kcal/mol) and methyl-furan ring (−9.25 kcal/mol; normalized score: - 1.53 kcal/mol) made comparatively smaller contributions. These data indicate that, although the peripheral fragments still participate in ligand stabilization, they are less critical for maintaining high-affinity binding than the central aromatic framework. Overall, the MM/GBSA decomposition analysis indicates that the central aromatic framework of L083-1287 is the principal driver of binding stability, whereas the normalized scores highlight the acetamide and thiadiazole fragments as compact but efficient contributors. These findings suggest that future lead optimization should preserve the central phenyl/difluorophenyl pharmacophoric core while prioritizing modification of the less efficient peripheral regions, particularly the methyl-furan/lactone side of the molecule, to improve binding efficiency, potency, and overall drug-like properties.

**Figure 12.**
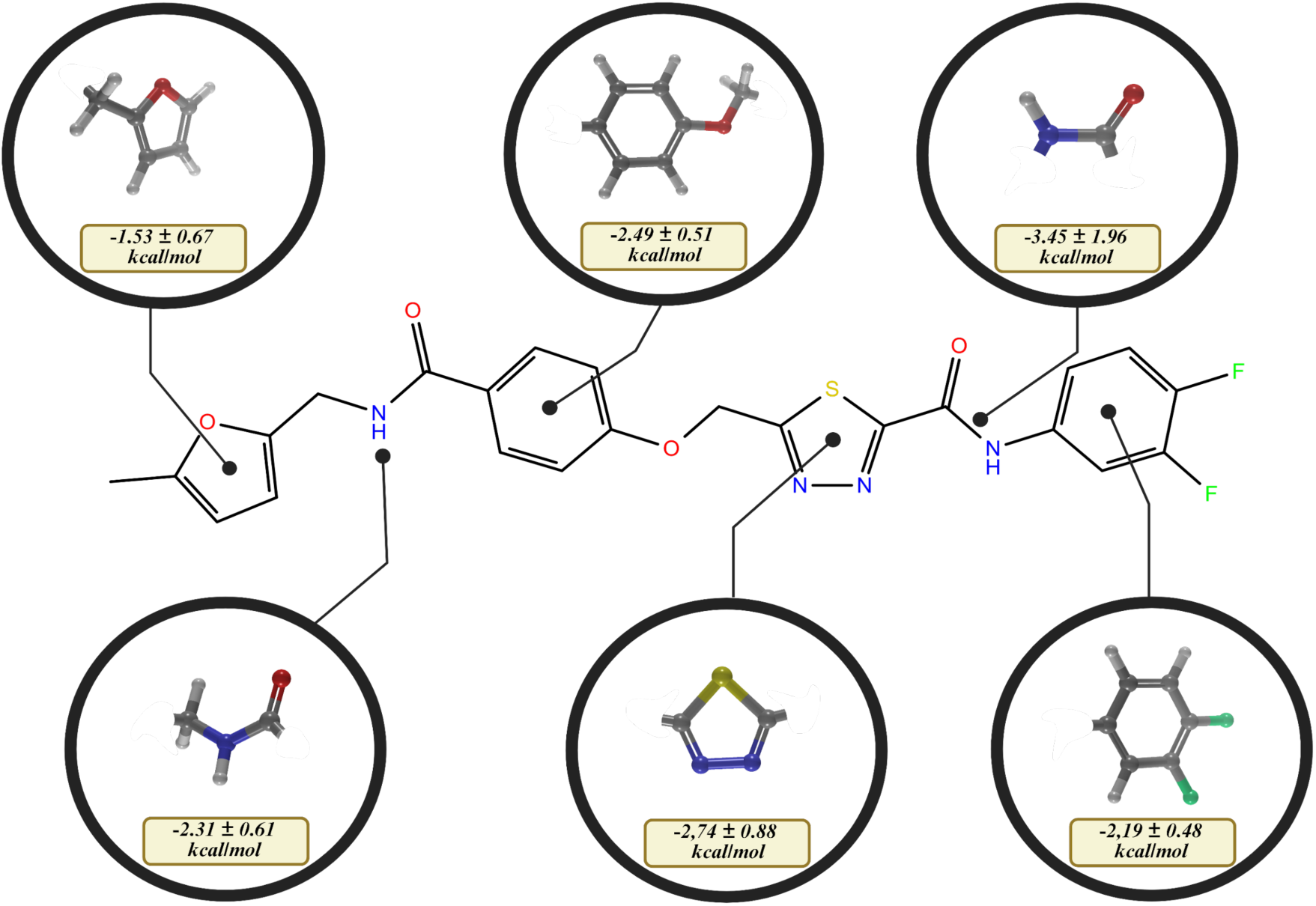
Fragment-based MM/GBSA binding free-energy contributions of lead compounds L083-1287 calculated from 200 ns MD simulations (3 replicas).

A similar fragment-based analysis of the second lead compound, L083-0077, also revealed a highly uneven energetic distribution, with binding dominated by a small subset of aromatic and heteroaromatic moieties. (Figure S13) In this case, the chloro-substituted phenyl ring (−22.60 kcal/mol), the ether-linked isopropoxy phenyl core (−22.79 kcal/mol), and the pyridinyl ring (−22.57 kcal/mol) emerged as the principal contributors to binding. The magnitude of these contributions suggests that the affinity of L083-0077 is driven largely by extensive aromatic contacts, potentially supported by hydrophobic interactions, π-stacking, and chlorine-mediated contacts within the binding pocket. By comparison, the 1,2,4-thiadiazole ring showed a more moderate contribution (−15.10 kcal/mol), while the amide linkage (−11.31 kcal/mol) and terminal methyl-amide fragment (−9.76 kcal/mol) contributed less strongly. As observed for L083-1287, this pattern indicates that the outer polar fragments of L083-0077 play a more supportive role, whereas the central aromatic scaffold serves as the principal energetic driver of target engagement.

Because MM/GBSA fragment energies are influenced not only by the quality of intermolecular contacts but also by fragment size, direct comparison of raw energy values may overestimate the contribution of larger moieties. To better distinguish intrinsic interaction efficiency from simple size effects, the fragment energies were therefore normalized to the number of non-hydrogen atoms, providing a ligand-efficiency-like metric for each structural element. A useful complementary interpretation emerges when the fragment contributions are normalized to the number of non-hydrogen atoms in each moiety, providing a ligand-efficiency-like view of the MM/GBSA data. In this size-normalized framework, the largest absolute contributors are not always the most efficient interaction units. For L083-1287, the central phenyl linker remains highly efficient at −2.49 kcal/mol per heavy atom, confirming that this ring is not only a major absolute contributor but also a dense energetic hotspot. By contrast, the para-difluoro-substituted phenyl ring, although strongly favorable in absolute terms, contributes more modestly on a per-atom basis (−2.19 kcal/mol per heavy atom), suggesting that part of its strong total contribution reflects its larger size. Notably, the 1,2,4-thiadiazole ring shows comparatively strong normalized efficiency (−2.74 kcal/mol), indicating that this smaller heteroaromatic unit makes disproportionately favorable contacts relative to its size. A similar trend is observed for L083-0077, where the pyridinyl ring and chloro-phenyl fragments emerge as the most efficient motifs on a normalized basis (−3.76 kcal/mol), followed by the 1,2,4-thiadiazole ring (−3.02 kcal/mol). In contrast, the larger ether-linked isopropoxy phenyl core, despite its strong absolute MM/GBSA contribution, appears less dominant when corrected for fragment size. Taken together, this normalized analysis suggests that while the larger aromatic cores account for much of the total binding energy through extensive surface complementarity, smaller heteroaromatic fragments may represent the most energetically efficient binding elements. This distinction is important for optimization, as it indicates that improvements in ligand efficiency may be achieved not only by preserving the major hydrophobic anchor regions but also by prioritizing compact fragments that deliver stronger binding per atom added.

Importantly, comparison of the parent compound L083-1287 with its optimized analog L083-0077 indicates that L083-0077 largely follows the pharmacophore optimization strategy suggested by the fragment-based analysis of L083-1287, while also revealing a distinction between target engagement and cellular activity. The decomposition of L083-1287 identified the central aromatic framework as the principal binding determinant and suggested that peripheral regions, particularly the methyl-furan/lactone side of the molecule, could be modified to improve binding efficiency and drug-like properties without disrupting the key pharmacophoric core. Consistent with this rationale, L083-0077 preserves a dominant aromatic/heteroaromatic binding architecture and redistributes the major energetic contributions toward the chloro-substituted phenyl ring, ether-linked isopropoxy phenyl core, and pyridinyl moiety, which together support stronger and more efficient direct engagement with the c-MET EC domain. This interpretation is further supported by the SPR results, where L083-0077 displays relatively a more favorable K_D_ value than L083-1287, indicating improved biochemical binding affinity. However, the MTT assays show that L083-1287 has a better IC_50_ profile than L083-0077, suggesting that enhanced target binding does not directly translate into superior cellular potency. This difference may reflect additional cell-based determinants such as permeability, intracellular exposure, solubility, metabolic stability, or context-dependent effects on receptor inhibition. Therefore, L083-0077 can be considered a pharmacophore- and binding-mode-optimized derivative of L083-1287, but not yet a fully cell-potency-optimized compound. These findings support the use of L083-0077 as a structurally improved scaffold for further optimization aimed at retaining its superior direct c-MET binding while improving its cellular efficacy.

Taken together, these analyses show that binding of both lead compounds is governed primarily by a limited number of core aromatic fragments rather than by uniform contributions across the entire scaffold. For L083-1287, the phenyl linker and para-difluoro phenyl group appear to define the main binding anchor, whereas in L083-0077 the dominant energetic contributions arise from the chloro-phenyl, isopropoxy phenyl core, and pyridinyl moieties. From a medicinal chemistry perspective, these findings can be used for future optimization efforts. Such as peripheral heterocyclic and amide-containing substituents may provide greater room for chemical diversification, potentially enabling improvements in selectivity, solubility, or pharmacokinetic properties without causing a major loss in target binding. Overall, the fragment-resolved MM/GBSA data define the structural hotspots that underlie ligand recognition and provide a rational basis for the next round of lead optimization.

### 3.14 Diffusion-Based Ligand Efficiency Assessment with Reference to Collision Theory

In addition to binding affinity and interaction energetics, the dynamic mobility of small-molecule inhibitors is an important parameter that can influence target engagement under biologically relevant conditions. According to collision theory, productive ligand–protein interactions depend not only on the strength of binding once contact is established, but also on how frequently ligand molecules can diffuse through the surrounding medium and encounter their target. For this reason, evaluating ligand diffusion behavior provides complementary information to static binding analyses, particularly in molecularly crowded and protein-rich environments where translational mobility may influence the probability of effective molecular collisions. We therefore analyzed the diffusion properties of the experimentally tested inhibitor candidates to assess their relative mobility and to estimate how efficiently they may sample the surrounding environment prior to binding. (Figure 13)

**Figure 13.**
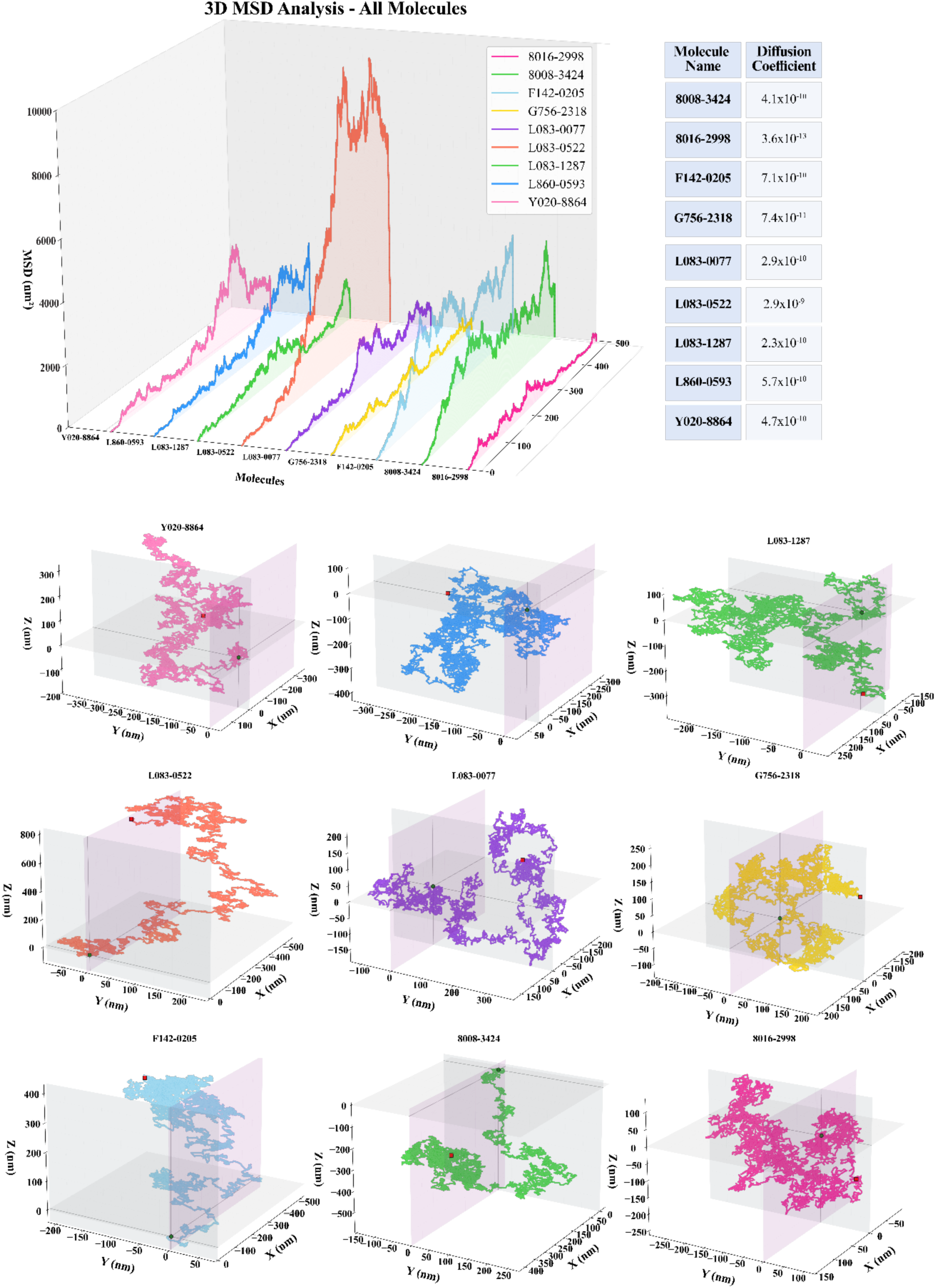
Diffusion coefficient profiles of candidate lead inhibitors over 500 ns MD simulations. (top) The 3D comparison chart compares the ability of molecules to move within chemical space in the presence of an unlimited solvent during 500 ns simulations: Molecules are classified with colors. Table lists diffusion coefficient of the molecules. (bottom) Separately plotted figures for each molecule of the MSD calculations.

To this end, mean square displacement (MSD) profiles were extracted from 500 ns MD simulations, and diffusion-related behavior was compared across the compound set. The MSD curves revealed clear heterogeneity among the tested molecules, indicating that the compounds do not move through the simulated environment with equivalent efficiency. Molecules displaying higher diffusion coefficients, in the range of approximately 4.0 × 10⁻¹⁰ to 8.0 × 10⁻¹⁰ m²/s, showed broader and more steeply rising MSD profiles, consistent with greater translational freedom and faster spatial exploration. Such behavior suggests that these compounds may achieve a higher frequency of random encounters with surrounding macromolecules, including the target receptor, which could favor initial collision events in accordance with diffusion-controlled interaction models. L083-0077 and L083-1287 showed comparatively restricted mobility, with diffusion-related values of approximately 2.9 × 10⁻¹⁰ and 2.3 × 10⁻¹⁰, respectively. Their shallower diffusion coefficient profiles indicate slower translational motion relative to the more mobile candidates. From the perspective of collision theory, this reduced diffusional behavior may decrease the frequency of ligand–target encounters over a given time window, particularly in crowded biological settings where molecular movement is already constrained by solvent organization, macromolecular obstacles, and transient nonspecific interactions. At the same time, lower diffusion is not necessarily disadvantageous in absolute terms, as reduced mobility may also reflect physicochemical features such as increased molecular compactness, stronger solvent interactions, or conformational stability, all of which can influence ligand behavior in solution. These findings therefore suggest that the candidate inhibitors differ not only in their binding characteristics but also in their capacity to dynamically access the target. In this context, diffusion analysis adds an important kinetic dimension to ligand evaluation: whereas MM/GBSA and docking-based approaches primarily describe the favorability of the bound state, MSD-derived diffusion profiles provide insight into how efficiently a molecule may reach the target in the first place. Accordingly, compounds with higher diffusional mobility may benefit from more frequent target collisions, while compounds with lower mobility may rely more heavily on stronger or more persistent binding once engagement occurs. Together, these results underscore that ligand efficiency should be considered as a combined function of both transport properties and binding energetics, and that optimal inhibitor design may require balancing sufficient molecular mobility with strong and selective target recognition.

### 3.15 Ligand-based similarity-guided prediction of specificity through off-target affinity mapping

To evaluate the potential off-target profiles of the two lead compounds (L083-1287 and L083-0077), similarity ensemble analysis (SEA) was employed as a ligand-based chemoinformatic approach to identify proteins with statistically significant similarity to known ligand sets. This analysis generated probability-based association scores that enabled the prioritization of putative secondary targets. For L083-1287, SEA indicated potential off-target interactions with PTPRE, SIRT3, SIRT2, FFAR1 (GPR40), and HSPB1. (Figure S14A) In parallel, L083-0077 was predicted to interact with NAMPT, PTPRE, SIRT3, SIRT2, and FFAR1 (GPR40). (Figure S14B) These results suggest that, although the two compounds share part of their predicted off-target landscape, each molecule also exhibits a distinct target-association profile, reflecting subtle differences in their underlying chemotypes. To further interrogate the plausibility of these predicted interactions at the structural level, each shortlisted off-target protein was matched with its corresponding experimentally resolved three-dimensional structures. These structures served as reference systems for subsequent computational validation. The predicted binding behavior of both lead compounds was then examined through triplicate 20 ns MD simulations, allowing assessment of ligand accommodation and stability within the target binding pockets under dynamic conditions. In addition, MM/GBSA binding free-energy calculations were performed to estimate the relative strength of compound–target interactions across the candidate off-target panel.

MM/GBSA binding free-energy calculations further substantiated the off-target interaction potential of both lead compounds across the selected protein panel. For L083-1287, the estimated binding free energies were −63.55 kcal/mol for GPR40 (FFAR1), −61.36 kcal/mol for NAMPT, −76.64 kcal/mol for SIRT2, and −68.10 kcal/mol for SIRT3. In comparison, L083-0077 showed consistently more favourable binding free-energy values against the same targets, with calculated MM/GBSA values of −78.10 kcal/mol for GPR40 (FFAR1), −71.61 kcal/mol for NAMPT, −82.37 kcal/mol for SIRT2, and −73.30 kcal/mol for SIRT3. These results indicate that both compounds are capable of forming energetically stable complexes with all four evaluated off-target proteins, with L083-0077 demonstrating a generally stronger predicted binding profile than L083-1287. When benchmarked against the respective co-crystallized reference ligands of these targets, both lead molecules displayed binding free-energy profiles that were broadly comparable. Specifically, the reference ligands 2YB for GPR40 (−87.35 kcal/mol), 2RM for NAMPT (−59.89 kcal/mol), KZU for SIRT2 (−68.10 kcal/mol), and SR7 for SIRT3 (−68.19 kcal/mol) provided a basis for contextualizing the calculated affinities. Relative to these controls, both lead compounds exhibited similar or modestly improved predicted binding energetics across the investigated targets, particularly for NAMPT, SIRT2, and SIRT3. Overall, these findings reinforce the molecular dynamics observations and support the notion that both L083-1287 and L083-0077 possess a notable capacity to engage multiple off-target proteins with appreciable affinity. (Table S2)

Collectively, these structure-based analyses supported the SEA predictions and showed that both L083-1287 and L083-0077 maintained favorable binding across all five evaluated off-target proteins, with consistently high binding scores obtained from the post-simulation energy analyses. (Table S2) These findings indicate that the two lead compounds may possess measurable polypharmacological potential beyond their primary intended targets, underscoring the importance of incorporating off-target profiling into the early-stage assessment of compound specificity, safety, and mechanistic selectivity.

### 3.16 hERG liability prediction and interaction profiling

In light of the developmental phenotypes observed in the zebrafish embryo assay, including cardiac- and vascular-associated alterations, we performed an exploratory *in silico* evaluation of the human ether-à-go-go-related gene (hERG) channel liability. To assess the potential cardiotoxicity risk associated with the lead compounds L083-1287 and L083-0077, we investigated their propensity to inhibit the hERG potassium channel, a major antitarget in drug discovery because of its established association with QT-interval prolongation and drug-induced arrhythmia. *In silico* hERG liability was first evaluated using the Pred-hERG platform (https://predherg.labmol.com.br), which integrates QSAR-based classification and regression models trained on experimentally curated datasets. The binary classification model identified both compounds as potential hERG blockers, with predicted probabilities of 77.59% for L083-1287 and 56.96% for L083-0077. By contrast, the consensus weighted classifier did not assign either compound to a strong blocker category, whereas the weak multiclass model suggested that both ligands may display only weak hERG-blocking behaviour. These classification outputs were consistent with the regression-derived IC₅₀ estimates, which placed both compounds within the low micromolar range, with predicted pIC₅₀ values of 5.6 (IC_50_ = 2.52 μM) for L083-1287 and 5.4 (IC_50_ = 3.75 μM) for L083-0077. (Figure S15) These predicted hERG values should be interpreted together with the cell-based activity data to avoid overestimating cardiac liability. Although both L083-1287 and L083-0077 were predicted to have hERG-associated IC₅₀ values in the micromolar range, their cellular potency profiles differed substantially. For L083-1287, the MTT-derived IC₅₀ values ranged from 0.015 to 0.26 µM, which are well below the predicted hERG IC₅₀ value of 2.52 µM. This separation suggests a relatively favorable activity window, indicating that L083-1287 may exert its cellular effects at concentrations lower than those predicted to be associated with hERG inhibition. In contrast, L083-0077 showed weaker cell-based potency, with MTT IC₅₀ values ranging from 8.85 to 18.67 µM, which are above its predicted hERG IC₅₀ value of 3.75 µM. Therefore, despite the improved target-binding affinity of L083-0077, its lower cellular potency results in a less favorable predicted hERG safety margin compared with L083-1287.

Fragment-level interpretation of the regression models provided additional insight into the structural basis of the predicted hERG activity. For L083-1287, the principal contributions to the micromolar inhibition profile originated from the piperidine-containing region, in which the pyridine ring and the para-chloro-substituted phenyl moiety emerged as the dominant features associated with predicted hERG engagement. In contrast, the contribution to activity declined around the thiazole fragment and the central phenyl ring located along the fourth rotational branch, indicating a less prominent role for these regions in channel recognition. The highest localized predicted potency for this ligand was observed in the pyridine region. For L083-0077, elevated pIC₅₀ contributions were mapped primarily to the terminal phenyl ring substituted at the meta and para positions with fluorine atoms, as well as to the central phenyl ring, suggesting that aromatic and halogen-substituted regions are key determinants of the predicted interaction pattern with the hERG channel. (Figure S15)

To further interrogate these predictions at the structural level, both lead compounds were subjected to MD simulations and MM/GBSA binding free-energy calculations using the experimentally resolved Astemizole–hERG complex (PDB ID: 8ZYO) as a reference system.

Across three independent simulation replicates, the average binding free energy for Astemizole was calculated as −100.57 kcal mol⁻¹, consistent with its well-established high-affinity hERG blockade. Under the same conditions, L083-0077 yielded an average binding free energy of −99.57 kcal mol⁻¹, indicating a binding strength closely comparable to that of Astemizole, whereas L083-1287 displayed a somewhat weaker but still substantial interaction profile, with an average value of −84.70 kcal mol⁻¹. These data support the prediction that both compounds can engage the hERG channel with appreciable affinity, while also suggesting that L083-0077 poses the greater hERG liability of the two leads on the basis of its overall energetic similarity to the reference blocker. (Figure S15)

Detailed contact mapping showed that all three ligands engaged residues classically implicated in hERG blockade, including Thr623, Ser624, Val625, Tyr652, Ala653, and Phe656, distributed across channel subunits A–D. As expected, Astemizole exhibited the most extensive contact network, dominated by persistent interactions with Tyr652 and Phe656, which are recognized as critical aromatic residues for blocker stabilization within the channel cavity. L083-0077 displayed the next most pronounced interaction pattern and strong recurrent engagement of Tyr652 across all four subunits, together with substantial interactions involving Phe656 in chains B and C. In comparison, L083-1287 retained hydrogen-bonding interactions with Ser624 and Thr623, but showed a reduced capacity to establish π-cation interactions with Tyr652 relative to Astemizole. This diminished aromatic cationic stabilization likely contributes to its comparatively weaker overall binding energy. (Figure S15)

To characterize the dynamic similarity of ligand–residue contact patterns over the course of simulation, principal component analysis (PCA) was performed on the residue contact trajectories. (Figure S15) This analysis revealed that the interaction profile of L083-1287 diverged most strongly on the B and C subunits, indicating a distinct binding mode relative to the reference ligand. By contrast, L083-0077 clustered closely with Astemizole, consistent with a highly similar pattern of residue engagement and a comparable energetic profile. Notably, interactions involving the A subunit differed substantially between both lead compounds and Astemizole, whereas those associated with the D subunit remained relatively conserved across all ligands, suggesting that some elements of hERG binding are preserved despite ligand-dependent differences in orientation and contact preference.

Analysis of the protonation states provided a plausible mechanistic explanation for these interaction differences. Astemizole was predicted to undergo protonation at its piperidine nitrogen, thereby enabling a stabilizing π-cation interaction with Tyr652, an interaction frequently associated with high-affinity hERG blockers. In contrast, the corresponding amine functionalities of L083-1287 and L083-0077 remained predominantly unprotonated under the analysed conditions, which explains the absence of an equivalent π-cation interaction network. This difference in ionization behaviour likely underlies the modest reduction in binding affinity observed for the lead compounds relative to Astemizole, particularly in the case of L083-1287.

### 3.17 pH-dependent ligand-binding stability under tumour-microenvironment-relevant conditions

Because the proposed inhibitors target the ECD of c-MET, their binding behavior should be evaluated under conditions that more closely reflect the tumor microenvironment rather than only under neutral physiological pH. Solid tumors frequently exhibit EC acidification as a consequence of altered metabolism, hypoxia, and impaired perfusion, and such pH shifts can alter the protonation states of both ligand functional groups and receptor-side residues. These changes may influence hydrogen-bonding patterns, electrostatic contacts, aromatic interactions, and overall complex stability, thereby affecting whether a compound can maintain productive target engagement in a disease-relevant environment. Therefore, pH-dependent ligand-binding stability analysis was performed as a complementary mechanistic assessment to determine whether the lead compounds preserve their interaction profiles with the c-MET extracellular domain under tumor-microenvironment-relevant acidic conditions. This analysis provides an additional criterion for lead prioritization by distinguishing compounds whose predicted binding remains stable despite pH variation from those whose target engagement may be more sensitive to extracellular acidification. To assess whether the identified ligands retain target engagement under conditions that better reflect the biochemical landscape of solid tumours, the first set of candidate molecules was evaluated at physiological pH (7.0) and under acidic conditions (pH 6.0 and pH 5.5) representative of the tumour microenvironment. Across the compound set, acidification generally led to a reduction in binding free energy, indicating that ligand binding is, in most cases, sensitive to pH-dependent changes in the protonation states of both the ligand and the binding-site residues. Such energetic weakening is consistent with a partial disruption of the electrostatic and hydrogen-bonding network required for stable complex formation under acidic stress.

Among the compounds examined, L083-1287 showed the most robust pH-resilient binding profile. At pH 7.0, the ligand maintained highly favorable binding free energies of −70.4 kcal mol⁻¹ at 50 ns and −65.15 kcal mol⁻¹ at 200 ns. Importantly, this energetic stability was largely preserved under acidic conditions, with ΔG values of −62.96 kcal mol⁻¹ at pH 6.0 and −65.12 kcal mol⁻¹ at pH 5.5. (Table S3) The limited magnitude of these changes suggests that the interaction network formed by L083-1287 is comparatively resistant to protonation-induced perturbations, supporting the view that this ligand may retain productive binding even in the acidic extracellular milieu characteristic of tumours. By contrast, other compounds were markedly more vulnerable to pH-driven destabilization. 8008-3424 and L860-0593, in particular, exhibited pronounced reductions in binding free energy, with losses approaching ∼20 kcal mol⁻¹, indicative of substantially weakened complex stability upon acidification. (Table S3) This behaviour suggests that their binding modes rely more strongly on interactions that are disrupted or attenuated at lower pH. G756-2318 retained relatively strong binding under acidic conditions, pointing to an intermediate level of pH tolerance. In contrast, Y020-8864 displayed the weakest overall profile and the highest degree of pH sensitivity, indicating that its target engagement is strongly compromised under tumour-relevant acidic conditions. (Table S3) Upon examination of the pH-dependent changes applied to the initial conformations, it was observed that the protonation states of all protein–ligand complexes varied across pH 5.5, 6, and 7. (Tables S4-11 and Supplementary Excel File 1)

Collectively, these findings identify extracellular pH as a critical determinant of ligand-binding stability and highlight the importance of evaluating candidate molecules under microenvironment-relevant conditions rather than relying exclusively on neutral-pH simulations. From a translational perspective, ligands such as L083-1287, which preserve favourable binding energetics across a physiologically relevant pH range, may offer a greater likelihood of maintaining pharmacological activity in vivo within acidic tumour niches.

### 3.18 Mechanistic validation of the intrinsic regulatory dynamics of c-MET

To understand how extracellularly binding small molecules inhibit c-MET, it was necessary to go beyond static structural models and examine the intrinsic conformational behavior of the ectodomain. Given the pronounced flexibility of the SEMA–PSI–IPT architecture and the likely regulatory role of disulfide-constrained hinge motions, mechanistic analysis was required to determine whether c-MET samples pre-existing inactive-like conformations and whether ligand binding stabilizes these states. This approach therefore provided the conceptual bridge between compound binding and receptor inhibition, allowing us to interpret lead-compound activity in the context of the native dynamic regulatory mechanism of c-MET.

Metadynamics and well-tempered metadynamics analyses revealed that the c-MET ectodomain possesses substantial intrinsic conformational plasticity, driven by multiple hinge elements and regulatory loops whose local motions propagate in a cascade-like manner to generate dominant global rearrangements. Rather than behaving as an unconstrained flexible assembly, however, the ectodomain appeared to operate within a mechanically regulated framework in which extensive mobility was channeled and limited by disulfide-bonded cystine networks. These data indicate that disulfide-linked structural elements are not merely passive stabilizers of fold integrity, but active determinants of the conformational landscape accessible to c-MET. The corresponding structural definitions and mechanistic representations are summarized in Figures S16 and S17. A first prominent feature of this framework was the stabilizing role of the PSI domain and adjacent cystine-rich elements. On the PSI-facing side, the disulfide pairs Cys526–Cys561, Cys529–Cys545 and Cys520–Cys538 formed, together with hydrophobic loops on the SEMA side, a compact hydrophobic core that prevented structural disintegration under steric stress while still permitting controlled rotational motion. (Figure S16E) A second constraint was provided by the R26-L segment, which behaved as a robotic arm-like structural limiter (Figure S16E) and restricted excessive SEMA–PSI–IPT1 rotation through the Cys26–Cys584 linkage. (Figures S17) This region additionally contributed a hydrophobic pocket-like environment under high-amplitude hinge motion, thereby reducing the risk of collapse. Within these constraints, the SEMA domain underwent a clockwise rotation of approximately 70° relative to the starting state; beyond this range, the PSI domain and R26-L region imposed substantial energetic barriers that suppressed further motion. (Figure 14C) Free-energy surface analysis further indicated that an additional clockwise rotational component of approximately 40° could be sampled before the system became trapped by high-energy barriers, consistent with an overall angular displacement of approximately 110° for the SEMA–PSI–IPT1 module. (Figures S18 and S19) By contrast, the IPT1–IPT2 junction behaved as a distinct rotational center with much less apparent restriction.

**Figure 14.**
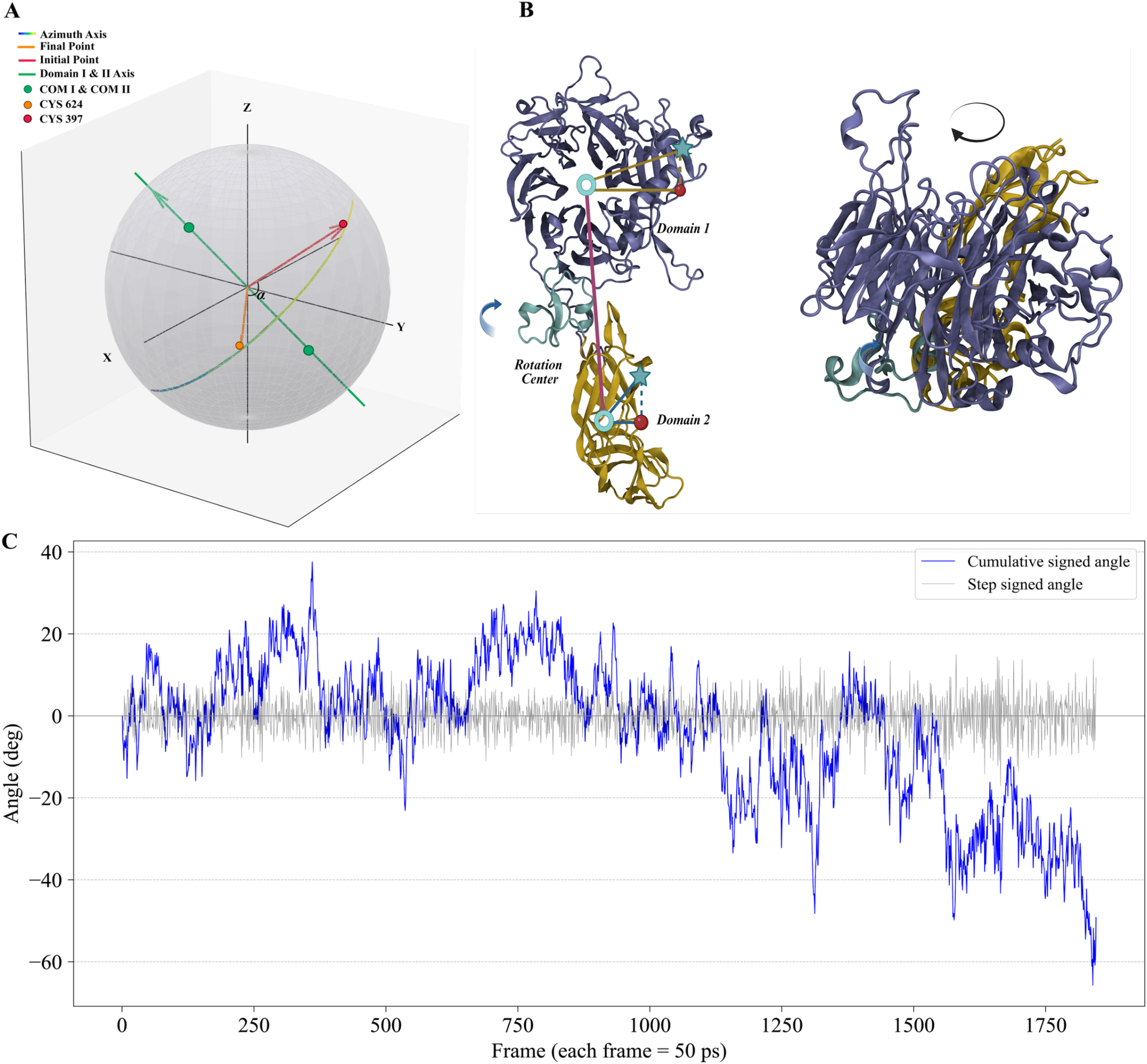
Time-dependent rotational dynamics were analyzed using representative metadynamics simulation trajectories. **A)** A simplified Bloch sphere representation was used to describe the rotational behavior between Domain 1 and Domain 2. The red vector was defined between the center of mass of Domain 1 (COM1) and a structurally stable reference point located near the domain interface (Cys397). The orange vector was defined from the center of mass of Domain 2 (IPT1–IPT2 combined, COM2), selected based on positional stability. The green axis represents the line connecting COM1 and COM2. The rotation axis (azimuth) is visualized using a gradient color transition. **B)** A three-dimensional structural representation is provided to facilitate interpretation of the rotational motion. The PSI domain, acting as the rotational center, is highlighted in cyan. The centers of mass of Domain 1 and Domain 2 are shown as cyan annuli. Red stars indicate cysteine residues, while red lines represent vector projections. A top-down view of the protein is also included to illustrate the rotational orientation and direction. **C)** The rotation of the system was quantified relative to the centers of mass using a signed angular metric. Frame-by-frame fluctuations are shown in gray, while the cumulative signed rotation angle is overlaid to capture the net rotational behavior throughout the trajectory. The y-axis represents the α angle defined in panel (A), corresponding to the rotational displacement, and the x-axis represents the simulation time. Each frame corresponds to a 50 ps interval.

The simulations further showed that motion across the SEMA domain is not localized to a single structural element, but is redistributed through coupled communication among Interfaces I (Figure S16B), II (Figure S16D) and III (Figure S16C). In particular, the 367–377 helix (Figure S16A) emerged as a key motion-transfer element that was responsive to structural changes originating from both Interfaces I and III and tilted by approximately 60° during the transition. (Figure 14C) This rearrangement was accompanied by compaction between Interfaces I and II and by pronounced fluctuations in the Lys306–Glu616 distance, which ranged from 72 Å to 1.6 Å to, consistent with repeated barrier crossing and exchange between local free-energy wells. (Figure 15A) Mechanistically, Asp270 in Interface I engaged the SDRG-L270 (Figure S16F) segment of Interface II, while SDRG-L358 (Figure S16F) cooperated with SDRG-L270 to form a hydrophobic core. (Figure 15B, 15C) Engagement between RG-L306 in Interface III and RG-L616 increased hinge-like motion, which could then be transmitted back towards Interface I through angular changes in the 367–377 helix. (Figures 15B and 15C) During the first quarter of the simulation, before major barrier crossing, the distances of SDRG-L358 and SDRG-L270 from the Interface I reference residue HIS388 were negatively correlated with the RG-L306–RG-L616 distance. (Figure 15C) Although this correlation weakened during the second and third quarters as the system crossed successive barriers, it re-emerged during the final quarter, supporting a recurrent and dynamically regulated mechanism of interfacial coupling.

**Figure 15.**
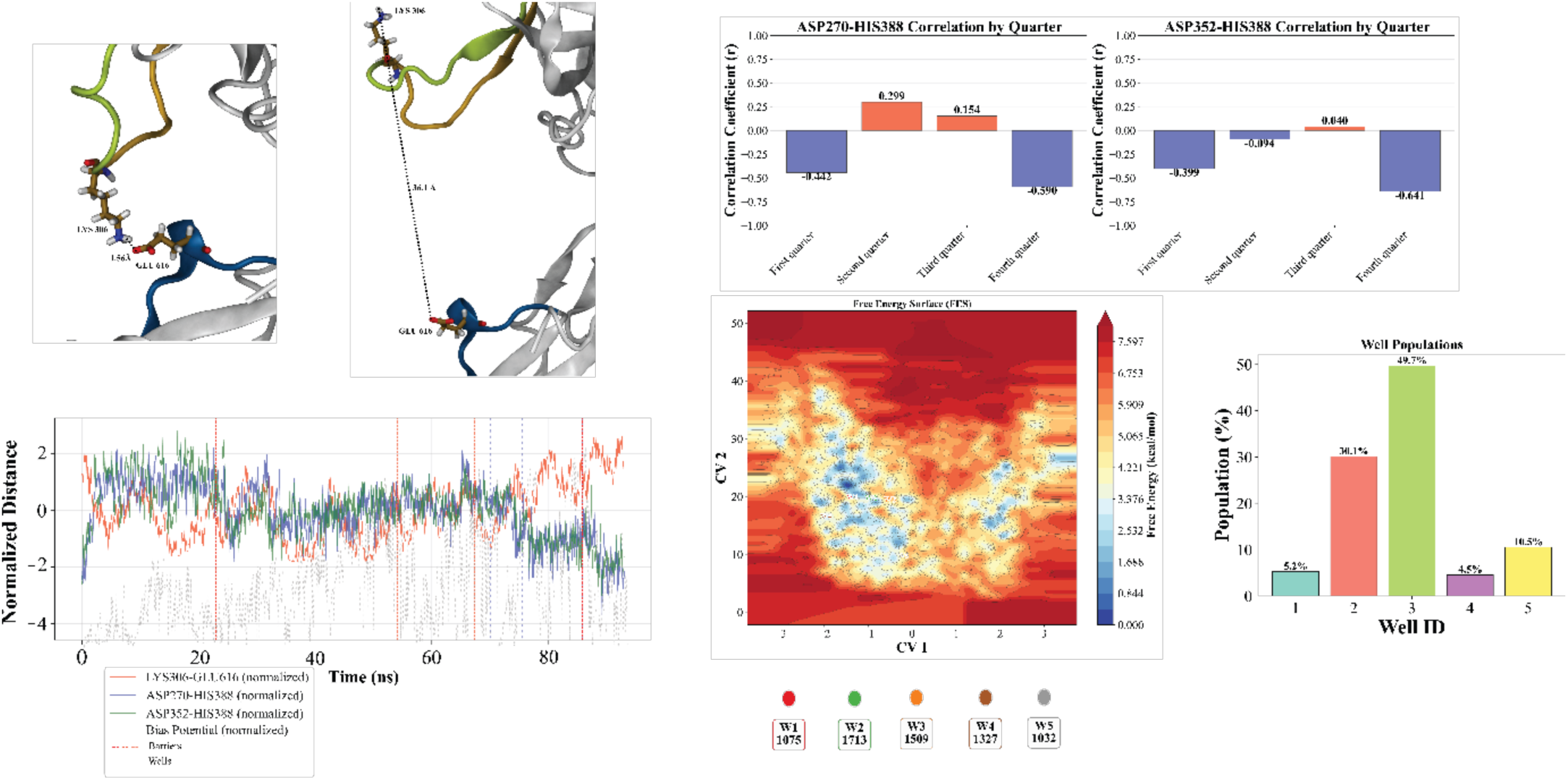
Evidence for the sequential angular hinged-rotation mechanism. **(A)** Structural snapshots illustrating the angular hinge motion responsible for coordinated SEMA-PSI-IPT domain contraction, driven by the Regulatory Loops RG-L 614-618 and RG-L 298-312. The dominant residues GLU616 (RG-L 614-618) and LYS306 (RG-L 298-312) exhibit large distance fluctuations across the simulation: (i) first frame and (ii) frame 332. **(B)** Time evolution of normalized distances during the 93.2 ns single-walker metadynamics simulation. The x-axis indicates simulation time (ns), and the y-axis shows normalized distance variations. The orange trace represents the LYS306-GLU616 distance. The blue trace corresponds to the distance between the dominant residue ASP270 (SDRG-L 265-276) and the reference residue HIE388, whereas the green trace represents the distance between ASP352 (SDRG-L 346-361) and HIE388. Red-orange dashed boxes mark energy barrier-crossing regions, while purple dashed boxes indicate free-energy wells (local minima). **(C)** Correlation analysis between loop-distance variations and hinge motion. Bar plots report the correlation between ASP270-HIE388 and ASP352-HIE388 distances with the LYS306-GLU616 distance, evaluated across simulation quarters (x-axis). The y-axis denotes the correlation coefficient (r). **(D)** Metadynamics-derived free-energy surface (FES) with annotated wells and representative states. Identified wells are color-coded and mapped to corresponding frame indices. The colormap indicates free-energy levels, with blue representing lower energy and red representing higher energy. **(E)** Population analysis of sampled wells across the metadynamics trajectory. The x-axis lists well identities, and the y-axis reports the population fraction (%).

Frustration analysis provided an additional layer of mechanistic support by showing that disulfide-bonded cystine regions and neighboring regulatory loops display correlated behavior and cluster around pivot-like highly frustrated regions. Two principal pathways were identified as candidate conduits for hinge-like motion across the SEMA domain. The first originated from Interface I, corresponding to the ligand-binding region, and extended through Helix-366 towards a central pivot while incorporating portions of Interfaces II and III. A second, complementary pathway approached the same pivot from the opposite direction, indicating that the conformational transition is organized through convergent rather than purely linear communication routes. On the far side of this pivot, the PSI domain, which contains four disulfide bonds and several tryptophan residues, also appeared as a localized highly frustrated region, further supporting its role as an integrated mechanical regulator rather than an isolated stabilizing appendage. Within Interfaces I–III, the cystine pairs Cys282–Cys409, Cys385–Cys397 and Cys298–Cys363 were repeatedly embedded in these pathways and are therefore likely to function as motion-transducing elements within the ectodomain. (Figure S20)

The free-energy landscape was dominated by a limited number of recurrent conformational states, comprising one global minimum, one global minimum-like state, and three local minima. (Figures 15D, 15E) The corresponding well populations were W1, 5.3% (frame 1075), W2, 34.1% (frame 1713), W3, 49.7% (frame 1509), W4, 4.5% (frame 1327), and W5, 10% (frame 1032). (Figure 15E) Time-resolved reconstruction of the free-energy surface at 23.3, 46.6, 69.8 and 93.1 ns showed progressive sharpening of the dominant low-energy basin and increasing consistency of the overall landscape topology. (Figure S21) In the second half of the simulation in particular, both the position and width of the principal low-energy region were largely preserved, indicating convergence towards a specific conformational basin rather than unrestricted exploration of structurally unrelated states.

Collectively, these results support a model in which c-MET samples a mechanically constrained, sequential hinge-rotation mechanism in which disulfide-linked stabilizers, frustrated pivot regions and interfacial loop networks cooperate to direct the ectodomain towards a defined intrinsic inhibitory conformational regime.

### 3.19 Inhibitor Binding Recapitulates the Native Regulatory Cascade and Mimics a Rapid Intrinsic Inhibition Mechanism

Comparison of holo and apo metadynamics trajectories showed that inhibitor binding does not abolish the intrinsic dynamic program of the c-MET ectodomain, but instead can preserve and redirect it within the same regulatory framework. (Figure S20) In the inhibitor-bound state, the dominant dynamic features observed in apo c-MET were largely retained, most notably the fluctuations along the RG-L306/RG-L616 axis, indicating that ligand engagement does not override the native long-range motion network. (Figure 15A) Rather, binding of the optimized inhibitor L083-0077 appeared to bias this pre-existing conformational landscape towards a controlled inhibitory regime. In particular, the pyridine moiety of L083-0077 formed a specific interaction with the SDRG-L270 structural element through Asp270, thereby stabilizing the regulatory circuitry centered on Interface II and promoting coupling to the RG-L306/RG-L616 hinge axis. (Figures 15A and 15C) These findings suggest that the compound acts not by imposing an artificial conformational constraint, but by functionally hijacking an intrinsic loop-mediated cascade that is already encoded within the ectodomain architecture.

Across extensive apo-state metadynamics simulations, c-MET consistently sampled a dominant angular hinge motion, confirming that this behaviour represents a native property of the receptor rather than an inhibitor-induced artifact. Among trajectories showing pronounced rotational behaviour, three principal motion regimes were observed (Figure S22): (i) clockwise rotation of the SEMA–PSI–IPT1 region accompanied by anticlockwise rotation of IPT1–IPT2 (Figure S22A); (ii) clockwise rotation of both SEMA–PSI–IPT1 and IPT1–IPT2 (Figure S22B); and (iii) clockwise rotation restricted primarily to the SEMA–PSI–IPT1 region (Figure S22C). These patterns were reproduced across three independent simulations, and the resulting trajectories were mapped onto the final free-energy surface on a frame-by-frame basis. (Figure S23) Together, these data indicate that the c-MET ectodomain contains three major rotational centers, supporting a model in which global rearrangement emerges from coordinated but non-equivalent hinge and rotation axes.

Integration of these results with structural comparisons from the literature further supports the existence of a defined rotational hotspot around Gly517 and Gly519 at the SEMA–PSI–IPT1 junction. Whereas previously reported X-ray structures suggest an angular displacement of approximately 60° in this region, the present simulations indicate that conformational changes of up to approximately 110° may be accessible under the sampled conditions. (Figure S22) At the same time, the data suggest that the apparently “unlimited-like” cyclic behaviour described in non-tethered experimental settings is more likely to arise from motion at the IPT1–IPT2 axis rather than unrestricted rotation of the entire SEMA–PSI–IPT1 unit. This interpretation is consistent with the observed restraining influence of the R26-L region and PSI-associated disulfide-rich elements identified in the apo mechanistic analysis. Because the metadynamics systems were not membrane-tethered and were based on the 7MO7 ectodomain construct lacking IPT3–IPT4, the absolute amplitude of motion should be interpreted cautiously.

Metadynamics analyses revealed that the sequential angular hinge and rotational behaviours of c-MET differ substantially between the holo and apo conformational states. In unbiased classical simulations, rotational motion was not evident in the apo ensemble, whereas the holo state displayed a detectable, albeit limited, degree of rotation. Under metadynamics, however, this relationship was inverted. When literature-derived collective variables were applied and conformational sampling was expanded through multiple single-walker trajectories, the apo system underwent extensive sampling of both clockwise and counterclockwise rotational transitions. (Figures S22 and S23) These data suggest that the apo ectodomain possesses a broader intrinsic capacity for rotational rearrangement than is captured under conventional simulation conditions. (Figure S19)

In contrast, the holo state exhibited substantial ligand fluctuations during metadynamics, and only minimal rotational motion was observed. This reduced sampling of large-scale rotation in the presence of ligand suggests that ligand binding modifies the way in which conformational strain is distributed across the receptor. Rather than uniformly stabilizing the structure, ligand engagement may introduce an alternative balance of local stabilization and destabilization, thereby redirecting energy flow within the ectodomain. This interpretation is supported by the distinct frustration signatures obtained for the two states. (Figure S20) The observed differences are likely to arise from the interplay between ligand entropy and the imposed collective-variable framework, which can favour energy dissipation in some regions while amplifying it in others. Accordingly, the holo and apo conformations appear to populate different dynamic and energetic landscapes, as reflected in their representative frustration profiles.

From the inhibitor perspective, the two lead molecules differed sharply in their relationship to this native dynamic program. The current simulations did not indicate that L083-1287 strongly recapitulates the endogenous conformational cycle, as the angular hinge behaviour and the rotational features associated with the apo mechanism were not clearly observed. By contrast, L083-0077 reproducibly displayed angular hinge motion across multiple independent short-and long-timescale all-atom simulations, although clear rotational sampling was not observed under conventional MD conditions. This likely reflects a limitation of accessible sampling rather than a mechanistic incompatibility, because the compound nonetheless preserved the key loop-coupling features of the intrinsic regulatory network. Taken together, these observations position L083-0077 as a mechanistically privileged extracellular c-MET ligand that most closely reproduces the native inhibitory cascade of the receptor. In this sense, L083-0077 can be regarded as a positive-control-like small molecule for the c-MET ectodomain, providing a basis for future mechanism-guided screening strategies that prioritize compounds capable not only of binding the ectodomain, but also of recapitulating its intrinsic inhibitory dynamics.

## 4. Discussion

This study identifies the EC c-MET ectodomain as a tractable target for small-molecule inhibition and, in doing so, expands the conceptual space of c-MET-directed therapy beyond kinase blockade. Whereas most clinically advanced c-MET inhibitors act at the intracellular ATP-binding cleft, our data supports an upstream strategy in which ligand recognition and receptor activation are intercepted at the ectodomain itself. This is significant for two reasons. First, it shows that a receptor surface long viewed as a difficult protein–protein interaction landscape can, in fact, support stable and selective small-molecule engagement. Secondly, it suggests that EC c-MET inhibition is not merely a steric alternative to kinase inhibition, but a mechanistically distinct mode of control linked to the native conformational logic of the receptor. Across large-scale virtual screening, multistage MD, biochemical target-engagement assays, cellular validation, and SPR analyses we converge on the conclusion that EC c-MET is druggable and that productive inhibition is associated with stabilization of a pre-existing inhibitory program encoded within the ectodomain architecture.

Although EC targeting of MET has already been explored using biologic agents, the present small-molecule strategy provides a mechanistically distinct and potentially complementary alternative. Antibody-based approaches such as onartuzumab and emibetuzumab were designed to interfere with MET activation at the EC level, either by blocking HGF-dependent receptor activation or by promoting MET internalization and degradation; however, their clinical development has highlighted challenges related to patient selection, tumor heterogeneity, and translation of extracellular target engagement into durable clinical benefit. [63] For example, onartuzumab did not improve clinical outcomes in the phase III MET-Lung trial in MET-positive NSCLC, while emibetuzumab showed biologically relevant MET-directed activity but remained limited by the typical constraints of antibody-based therapeutics. [63] More recently, amivantamab has validated the extracellular MET/EGFR axis as a clinically actionable surface-targeted strategy, but it remains a large bispecific antibody whose activity also involves EGFR co-targeting and immune-effector mechanisms rather than selective small-molecule modulation of MET ectodomain conformational regulation. [64] In this context, the compounds described in this study are not intended to replace antibody therapeutics directly, but rather to expand the modality space for EC MET inhibition by offering chemically tunable, low-molecular-weight scaffolds that may provide advantages in tissue penetration, synthetic optimization, and fine control of hotspot-directed binding. The identification of L083-1287 and L083-0077 as EC-domain binders therefore supports a complementary therapeutic concept, such that small molecules capable of engaging MET regulatory surfaces and modulating receptor activation upstream of the kinase domain, while potentially avoiding some of the developability limitations associated with biologic MET inhibitors.

An important advance of the present work is that it moves the field from static hotspot mapping to dynamic mechanistic interpretation. The initial screening campaign was built around literature-supported interface residues, including the HGF-side hotspot set and c-MET residues such as Asn393 and Phe398, but the most informative outcome was not the recovery of binders to those residues alone. Rather, subsequent simulations, metadynamics, and holo–apo comparisons indicate that these sites are embedded in a larger interfacial communication network spanning Interfaces I, II and III of the ectodomain. Within that framework, Helix 367–377, the SDRG-associated loops around ASP270, the RG-L306/RG-L616 hinge axis, the PSI domain, and the disulfide-rich SEMA–PSI–IPT junction emerge as coupled mechanical elements that channel long-range conformational change. The implication is that EC inhibition of c-MET is not adequately described as local occupancy of a ligand-binding pocket. Instead, the most effective compounds appear to exploit a distributed inhibitory cascade in which local contacts are translated into a regulated global contraction of the receptor.

This mechanistic perspective helps explain why the three principal compounds emerging from the study should not be interpreted as equivalent leads. Early in the workflow, L083-1287, 8008-3424 and later L083-0077 all appeared compelling by docking, MM/GBSA and short-timescale simulation criteria. However, the subsequent layers of validation reveal a meaningful divergence between apparent affinity, mechanical stability, direct ectodomain engagement and mechanistic fidelity. L083-1287 is the clearest example of strong biological activity at the cellular level. It displayed submicromolar IC_50_ values in HuH7 cells across all tested time points, strongly reduced phospho-c-MET, and substantially suppressed long-term colony formation. These properties establish that the scaffold can produce robust anti-survival effects in an HGF-stimulated HCC setting. At the same time, its ectodomain binding by SPR was more modest than its cellular potency would predict, and its developmental liability in zebrafish was more pronounced than that of the other early hits tested. Taken together, these data raise the possibility that the exceptional cellular activity of L083-1287 may not derive exclusively from its intended EC c-MET interaction, but may also reflect additional pharmacology.

By contrast, L083-0077 emerges as the mechanistically most persuasive ectodomain ligand, even though it is not the most potent compound in the simple cell-viability readout. Lead optimization around the L083-1287 series did not improve cytotoxic potency in HuH7 cells; indeed, L083-0077 remained in the single- to low-double-digit micromolar range in that assay. Yet, SPR showed improved direct binding to recombinant c-MET ectodomain, with a lower K_D_ than either L083-1287 or 8008-3424 and with slower association and dissociation behavior, consistent with a more persistent target-engagement mode. More importantly, the simulation data place L083-0077 in a different mechanistic category. Its interaction network is centered on the literature-supported hotspots Asn393 and Phe398, but extends toward Asp270 and the surrounding regulatory elements in a way that appears to couple local binding to the broader inhibitory cascade. Fragment-based decomposition further supports this view by showing that the pyridyl and aromatic terminal elements make dominant contributions to binding, consistent with a scaffold organized to combine anchoring, conformational adaptability and long-range influence on receptor motion. In other words, L083-0077 may be less cytotoxic than L083-1287 in a single HCC line while still being the better EC probe compound.

The behavior of 8008-3424 is also informative, as it highlights the importance of iterative computational and experimental refinement during EC inhibitor discovery. In short-timescale simulations and sMD, 8008-3424 emerged as a promising candidate, engaging both c-MET-and HGF-side residues, showing substantial resistance to forced dissociation, and supporting a plausible interface-bridging mode of inhibition. Its direct binding to the c-MET ectodomain by SPR further reinforced its relevance as an active scaffold. In the longer validation simulations, however, this binding mode appeared less persistent over extended timescales, suggesting that the interaction may be more transient or context-dependent than initially appreciated. Rather than diminishing the impact of the early computational results, this progression underscores the complementary nature of multistage modeling approaches: early docking, short MD, and sMD are highly effective for identifying promising interaction patterns, whereas longer simulations can further refine prioritization by distinguishing ligands with sustained binding behavior from those with more dynamic modes of engagement. In this regard, 8008-3424 remains a biologically and mechanistically valuable scaffold, and its trajectory through the workflow highlights the strength of the tiered computational strategy used here for extracellular PPI-directed discovery.

One of the most interesting outcomes of the study is that extracellular c-MET inhibition appears to be most effective when it preserves, rather than overrides, the receptor’s endogenous dynamic grammar. Holo–apo comparisons indicate that c-MET intrinsically samples hinge-like and rotational motions, with major communication routes converging on a frustrated pivot architecture shaped by disulfide-linked regions and the PSI domain. In this setting, L083-0077 does not behave like an exogenous clamp that freezes the receptor into an artificial conformation. Instead, it appears to bias the native landscape towards a controlled inhibitory regime by stabilizing the same loop-coupling framework used by the apo receptor during its intrinsic motion cycle. This distinction matters. A ligand that suppresses signaling by co-opting an endogenous inhibitory trajectory may, at least in principle, be less likely to trigger maladaptive distortions, aggregation-prone states or compensatory escape routes than a ligand that inhibits through non-physiological deformation. Although this conclusion remains to be tested experimentally, it provides a rationale for mechanism-guided scoring schemes that go beyond energy ranking alone.

The tumor-microenvironment analyses further refine lead selection by showing that pH resilience is not a generic property of this chemical series. Acidification weakened binding for several compounds, particularly 8008-3424 and L860-0593, indicating that protonation-sensitive electrostatic interactions can sharply erode EC target engagement under tumor-relevant conditions. L083-1287 showed the clearest quantitative resilience in this assay set, maintaining favorable binding energies at pH 6.0 and 5.5. In the broader workflow, L083-0077 also remained compatible with the prioritized mechanistic framework under later validation. Together these results underscore an often underappreciated point: for EC oncology targets, persistence of binding under acidic conditions may be as important as nominal affinity at neutral pH. Future optimization should therefore treat pH robustness as a lead-defining parameter, not merely as a supplementary characterization.

The safety and selectivity data provides useful early filters, on one hand, ligand-based similarity scores were low and diffusion/orientation analyses argued against straightforward competition with known high-affinity off-target ligands. On the other hand, structure-based profiling suggested that both L083-1287 and L083-0077 can form energetically favorable complexes with several secondary targets, including NAMPT, SIRT2, SIRT3 and FFAR1. Likewise, the hERG analysis simulations placed L083-0077 in particular close to a known blocker-like energetic and contact pattern, even if other analyses suggested reduced π-cation stabilization and possibly weaker functional liability than canonical blockers.

Future studies are required to fully elucidate the off-target effects and broader biological impact of the proposed inhibitors. Although this study is limited by the use of a single cell line and a single autophosphorylation site, the data presented have profound value in underscoring c-Met activation kinetics. In addition, the zebrafish data are valuable as an early whole-organism liability readout, but because they are phenotypic and were performed only with selected early hits, they should be discussed as prioritization data rather than definitive on-target toxicology. Finally, the amplitude of the proposed conformational cycle should be interpreted with caution because of simulation time limitations. However, none of these limitations undermine the central conclusion, but they do define the boundary between what has been established here and what remains to be demonstrated.

Overall, this study provides more than a lead-identification campaign. It also offers a mechanistic framework for how small molecules can engage a dynamic receptor ectodomain and exploit its intrinsic inhibitory circuitry. By combining large-scale screening with long-timescale dynamics, steered dissociation, fragment-resolved energetics, tumor-microenvironment stress testing and mechanistic trajectory analysis, the work outlines a general route for converting apparently undruggable EC interfaces into tractable discovery problems. Within that framework, L083-1287 is best viewed as a potent biologically active scaffold that reveals the therapeutic promise and developmental liabilities of EC c-MET inhibition, whereas its analog L083-0077 is the more mechanistically privileged ectodomain ligand and the strongest foundation for future optimization.

## 5. Conclusions

This study establishes the EC c-MET ectodomain as a tractable target for small-molecule intervention and supports EC HGF–c-MET blockade as a mechanistically distinct alternative to conventional kinase-domain inhibition. By integrating large-scale virtual screening, multistage MD simulations, sMD analyses, free-energy profiling, and experimental validation, we show that small molecules can productively engage the c-MET EC region and suppress HGF-driven receptor activity in HCC models. These findings expand the therapeutic landscape of c-MET inhibition from intracellular catalytic blockade to upstream control of ligand-dependent receptor activation.

A central outcome of this work is that EC c-MET inhibition is not explained solely by static occupancy of a surface hotspot. Instead, our data support a broader inhibitory framework in which ligand binding is coupled to an intrinsic EC regulatory network spanning multiple c-MET ectodomain interfaces. Within this landscape, the computational analyses were not only effective for hit discovery, but also essential for resolving how different compounds interact with the receptor over time, under mechanical stress, and under tumor-relevant acidic conditions. In this sense, the study demonstrates the value of tiered computational modeling as both a discovery engine and a mechanistic tool for EC protein–protein interface targeting.

Among the compounds identified, L083-1287, 8008-3424, and L083-0077 represent distinct stages of lead evolution. L083-1287 emerged as a potent biologically active scaffold with strong effects on c-MET phosphorylation, cell viability, and clonogenic growth, thereby providing clear proof that EC directed small molecules can suppress this pathway in HCC cells. 8008-3424 further supported the tractability of the interface by showing direct ectodomain binding and early evidence of interface-bridging behavior, even though its longer-timescale binding profile suggested a more dynamic and less persistent interaction mode. In this context, L083-1287 is best interpreted as a biologically potent first-generation scaffold that establishes both the promise and the constraints of EC c-MET targeting: it demonstrates that pharmacological inhibition at the ectodomain is achievable and functionally relevant, yet it also exposes developmental liabilities that will need to be addressed during optimization. L083-0077, in contrast, represents the more mechanistically mature analog of parent compound L083-1287, with a mode of ectodomain engagement that maps more convincingly onto the proposed EC inhibitory cascade, thereby positioning it as the more compelling platform for future medicinal-chemistry refinement.

Overall, this work establishes the c-MET ectodomain as a tractable target for small-molecule intervention and provides a mechanistic foundation for EC c-MET inhibition as a distinct therapeutic strategy. Rather than simply delivering active compounds, the study shows that ligand engagement at the ectodomain can be understood in the context of a broader EC regulatory network that shapes receptor activation. More broadly, our findings demonstrate that dynamic, conformationally complex receptor–ligand interfaces, long considered difficult to target with small molecules, can be interrogated in a systematic and mechanistically informative manner through an integrated computational–experimental framework. These results position EC c-MET targeting as a promising route toward selective, upstream, and potentially resistance-limiting inhibitors, while also providing a transferable blueprint for small-molecule modulation of other disease-relevant extracellular protein–protein interactions.

