## Supporting Material for "First-in-Class Small Molecules Inhibit c-MET Activity Through Self-Regulatory Elements in Its Extracellular Domain"

Serdar Durdağı[1,2,7,8,*]

[1]Laboratory for Innovative Drugs (Lab4IND), Computational Drug Design Center (HİTMER), Bahçeşehir University, 34734, İstanbul, Türkiye; [2]Computational Biology and Molecular Simulations Laboratory, Department of Biophysics, School of Medicine, Bahçeşehir University, 34734, İstanbul, Türkiye; [3]Izmir Biomedicine and Genome Center, Izmir, 35340, Türkiye; [4] Department of Oncology, Georgetown University, Washington, DC., USA; [5] Galen Research Center, Izmir Tinaztepe University, Buca, Izmir, Türkiye; [6] Department of Medical Biology and Genetics, Faculty of Medicine, Izmir Tinaztepe University, 35340, Buca, Izmir, Türkiye; [7]Molecular Therapy Laboratory, Department of Pharmaceutical Chemistry, School of Pharmacy, Bahçeşehir University, 34353, İstanbul, Türkiye; [8]Quantitative System Biology Lab, Faculty of Medicine, Biruni University, İstanbul, Türkiye.

* (NA); (SD)

**Table S1.** Targeted binding surfaces, GBSA , docking and binary scores of promising potential drug molecules from three different binding modes (Only c-MET, HGF only, c-MET/HGF complex)

| Target and database | Molecule name | ΔG binding score (kcal/mol) | Docking Score (kcal/mol) | Binary Qsar Score |
|---|---|---|---|---|
| MET ChemDiv Interface I | 8016-2998 | -54.4831 | -4.01 | 0.79 |
| HGF-MET ChemDiv Interface I | Y043-7353 | -52.7152 | -4.3 | 0.62 |
| HGF-MET ChemDiv Interface I | Y020-8864 | -50.0091 | -3.73 | 0.64 |
| HGF-MET ChemDiv Interface I | 8008-8176 | -49.3189 | -3.93 | 0.78 |
| MET Enamine Interface I | 341750 | -48.9972 | -4.06 | 0.65 |
| HGF-MET ChemDiv Interface I | G290-0199 | -45.5067 | -3.6 | 0.65 |
| MET Enamine Interface I | 211437 | -44.8058 | -4.03 | 0.76 |
| HGF ChemDiv Interface I | Y043-5500 | -43.5347 | -5.57 | 0.69 |
| MET Enamine Interface I | 453691 | -43.0312 | -3.99 | 0.80 |
| MET ChemDiv Interface I | F499-0938 | -42.992 | -3.93 | 0.71 |
| HGF-MET Enamine Interface I | 134801 | -41.6471 | -5.04 | 0.76 |
| MET Enamine Interface I | 20834 | -41.5117 | -4.53 | 0.61 |
| HGF-MET Enamine Interface I | 421221 | -38.6964 | -6.18 | 0.63 |
| HGF-MET Enamine Interface I | 379402 | -37.6371 | -5.49 | 0.77 |
| HGF-MET Enamine Interface I | 319316 | -36.0645 | -4.85 | 0.63 |
| HGF-MET Enamine Interface I | 232031 | -34.5801 | -4.88 | 0.62 |
| HGF ChemDiv Interface I | 8005-9360 | -33.8589 | -7.52 | 0.74 |
| HGF-MET Enamine Interface I | 296157 | -32.0768 | -4.96 | 0.58 |
| MET Enamine Interface I | 31378 | -30.4191 | -4.15 | 0.69 |
| HGF Enamine Interface I | 310740 | -30.3748 | -4.77 | 0.52 |
| HGF ChemDiv Interface I | Y021-1697 | -27.446 | -6.05 | 0.61 |
| HGF Enamine Interface I | 276391 | -26.5734 | -6.63 | 0.54 |
| HGF ChemDiv Interface I | C327-0261 | -24.3193 | -5.54 | 0.83 |
| HGF ChemDiv Interface I | 8756-0066 | -23.3006 | -6.66 | 0.61 |
| HGF Enamine Interface I | 124791 | -17.8784 | -4.84 | 0.77 |
| HGF Enamine Interface I | 400803 | -17.3593 | -5.07 | 0.58 |
| HGF Enamine Interface I | 376489 | -16.062 | -4.44 | 0.74 |

**Table S2: Comparative MM/GBSA binding free energy results of lead compounds (L083-1287 and L083-0077) and reference ligands across predicted off-targets.**

| Molecule | $\Delta G_{avg}$ | $\sigma\Delta G$ | $\Delta G_{max}$ | $\Delta G_{min}$ | $\Delta G_{NS,avg}$ | $\sigma\Delta G_{NS}$ | $\Delta G_{NS,max}$ | $\Delta G_{NS,min}$ | Target | PDB Code |
|---|---|---|---|---|---|---|---|---|---|---|
| 2YB | -87.35 | 4.99 | -72.72 | -98.52 | -92.98 | 4.95 | -79.66 | -103.05 | GPR40 | 8EJC |
| L083-0077 ***(L)*** | -78.10 | 5.35 | -66.18 | -89.96 | -81.86 | 5.26 | -70.32 | -93.80 | GPR40 | 8EJC |
| L083-1287 ***(L)*** | -63.55 | 3.62 | -54.32 | -73.13 | -66.80 | 3.65 | -56.40 | -76.52 | GPR40 | 8EJC |
| L083-0077 ***(L)*** | -71.61 | 5.77 | -58.40 | -83.94 | -76.13 | 6.06 | -61.34 | -88.62 | NAMPT | 7PPG |
| L083-1287 ***(L)*** | -61.36 | 6.26 | -45.29 | -76.32 | -64.93 | 6.38 | -50.12 | -80.97 | NAMPT | 7PPG |
| 2RM | -59.89 | 4.61 | -48.52 | -71.86 | -63.19 | 4.94 | -51.03 | -75.57 | NAMPT | 7PPG |
| 1XD | -58.64 | 4.13 | -48.73 | -69.09 | -61.87 | 4.32 | -51.39 | -72.10 | NAMPT | 7PPG |
| 2P1 | -54.78 | 3.90 | -44.71 | -64.84 | -58.21 | 4.23 | -47.49 | -68.32 | NAMPT | 7PPG |
| DGB | -51.86 | 4.49 | -39.03 | -62.39 | -56.68 | 4.72 | -43.90 | -67.67 | NAMPT | 7PPG |
| 7YX | -42.82 | 6.48 | -29.87 | -62.44 | -46.74 | 6.75 | -33.25 | -68.28 | NAMPT | 7PPG |
| IS1 | -38.88 | 7.50 | -20.06 | -57.53 | -44.62 | 7.74 | -25.70 | -63.36 | NAMPT | 7PPG |
| AQ1 | -23.20 | 3.96 | -14.40 | -33.90 | -24.67 | 4.43 | -15.94 | -37.39 | NAMPT | 7PPG |
| L083-0077 ***(L)*** | -82.37 | 8.35 | -59.88 | -98.00 | -86.04 | 8.31 | -63.74 | -101.99 | SIRT2 | 5MAR |
| L083-1287 ***(L)*** | -76.64 | 8.65 | -55.07 | -99.00 | -81.30 | 8.82 | -58.38 | -104.24 | SIRT2 | 5MAR |
| KZU | -68.10 | 7.25 | -50.80 | -85.92 | -73.62 | 7.24 | -58.21 | -91.95 | SIRT2 | 5MAR |
| 8K9 | -58.73 | 6.71 | -41.19 | -74.36 | -62.48 | 7.04 | -44.93 | -78.30 | SIRT2 | 5MAR |
| 7KE | -56.48 | 5.15 | -45.87 | -68.32 | -58.49 | 5.31 | -47.79 | -71.45 | SIRT2 | 5MAR |
| A2X | -53.60 | 7.36 | -37.50 | -71.49 | -57.06 | 7.34 | -40.35 | -75.84 | SIRT2 | 5MAR |
| 8NO | -51.85 | 5.40 | -35.42 | -66.84 | -55.03 | 5.57 | -37.47 | -70.96 | SIRT2 | 5MAR |
| A2I | -50.09 | 7.53 | -34.86 | -68.53 | -53.69 | 7.92 | -37.53 | -75.96 | SIRT2 | 5MAR |
| L083-0077 ***(L)*** | -73.30 | 7.49 | -56.22 | -90.50 | -76.67 | 7.42 | -60.44 | -93.53 | SIRT3 | 4BN5 |
| SR7 | -68.19 | 5.30 | -55.37 | -81.31 | -71.13 | 5.15 | -58.61 | -84.49 | SIRT3 | 4BN5 |
| L083-1287 ***(L)*** | -68.10 | 8.32 | -47.73 | -89.14 | -71.68 | 8.55 | -50.46 | -93.37 | SIRT3 | 4BN5 |

**Table S3:** Binding free energy values of the first set of candidate molecules under different tumor-relevant pH conditions. Binding scores were calculated from MD simulations of 50 ns at pH 7.0, pH 6.0, and pH 5.5, as well as extended 200 ns simulations at pH 7.0. Values represent MM/GBSA binding free energies derived from three replicates for each ligand-protein complex.

| 2D Structures | Target and Database | Molecule Name | ΔG Binding Score pH 7 50 ns (kcal/mol) | ΔG Binding Score pH 6 50 ns (kcal/mol) | ΔG Binding Score pH 5.5 50 ns (kcal/mol) | ΔG Binding Score pH 7 200 ns (kcal/mol) |
|---|---|---|---|---|---|---|
| 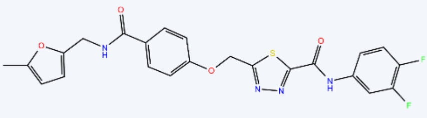 | MET ChemDiv | L083-1287 | -70,4 | -62.96 | -65.12 | -65.15 |
| 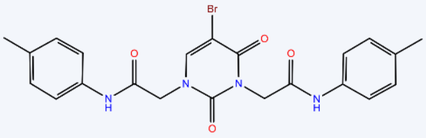 | HGF-MET ChemDiv | 8008-3424 | -60.80 | -47,15 | --57.54 | -42.08 |
| 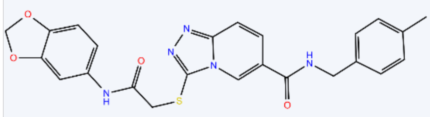 | MET ChemDiv | L860-0593 | -59,52 | -54.13 | -48.54 | -42.08 |
| 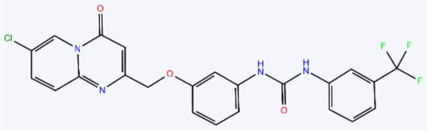 | MET ChemDiv | G756-2318 | -55,81 | -65.43 | -59.54 | -53.20 |
| 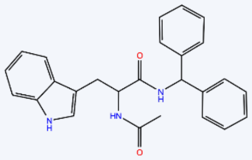 | HGF-MET ChemDiv | 8016-2998 | -54.48 | -57.84 | -56.35 | -57.31 |
| 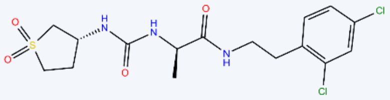 | MET ChemDiv | Y043-7353 | -52.71 | -61.44 | 48.53 | -66,45 |
| 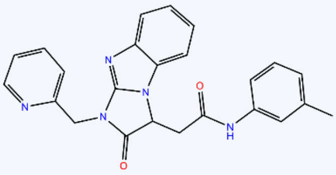 | MET ChemDiv | Y020-8864 | -50.00 | -49.75 | -54.65 | -37.92 |

**Table S4.** Protonation state differences of selected residues at varying pH levels used to generate the initial MD structures of the c-MET–HGF complex with 8008-3424.

| HGF | | | | | | | |
|---|---|---|---|---|---|---|---|
| **Residue** | **pH 7** | **pH 6** | **pH 5.5** | **Residue** | **pH 7** | **pH 6** | **pH 5.5** |
| A:158 | HIE | HIP | HIP | A:225 | HIE | HIP | HIP |
| A:257 | ASP | ASH | ASH | A:277 | HIS | HIP | HIP |
| A:335 | HIS | HIP | HIP | A:387 | HIE | HIP | HIP |
| A:423 | HIS | HIS | HIP | A:425 | HIS | HIP | HIP |
| A:551 | HIE | HIP | HIP | A:554 | HIS | HIP | HIP |
| A:633 | HIS | HIP | HIP | A:670 | GLU | GLU | GLH |
| A:682 | HIE | HIS | HIP | A:717 | HIS | HIE | HIE |

| c-MET | | | | | | | |
|---|---|---|---|---|---|---|---|
| **Residue** | **pH 7** | **pH 6** | **pH 5.5** | **Residue** | **pH 7** | **pH 6** | **pH 5.5** |
| B:61 | HIS | HIS | HIP | B:123 | ASP | ASH | ASH |
| B:148 | HIE | HIP | HIP | B:150 | HIE | HIE | HIP |
| B:159 | HIE | HIE | HIP | B:212 | HIS | HIP | HIP |
| B:289 | HIS | HIE | HIE | B:394 | HIE | HIE | HIP |
| B:396 | HIS | HIE | HIS | B:484 | HIE | HIE | HIP |
| B:489 | GLU | GLH | GLH | B:542 | HIS | HIP | HIP |
| B:633 | HIE | HIE | HIP | B:644 | HIE | HIE | HIP |
| B:689 | HIS | HIS | HIP | | | | |

**Table S5.** pH-dependent protonation state assignments of selected residues defining the initial MD conformations of c-MET bound to L860-0593.

| c-MET | | | | | | | |
|---|---|---|---|---|---|---|---|
| **Residue** | **pH 7** | **pH 6** | **pH 5.5** | **Residue** | **pH 7** | **pH 6** | **pH 5.5** |
| B:61 | HIS | HIS | HIP | B:123 | ASP | ASH | ASH |
| B:148 | HIE | HIP | HIP | B:150 | HIE | HIE | HIP |
| B:159 | HIE | HIE | HIP | B:212 | HIS | HIP | HIP |
| B:267 | GLU | GLU | GLH | B:289 | HIS | HIE | HIE |
| B:394 | HIE | HIE | HIP | B:396 | HIS | HIE | HIS |
| B:484 | HIE | HIE | HIP | B:489 | GLU | GLH | GLH |
| B:542 | HIS | HIP | HIP | B:633 | HIE | HIE | HIP |
| B:644 | HIE | HIE | HIP | B:689 | HIS | HIS | HIP |

**Table S6.** Assignment of residue protonation states under different pH conditions for the initial conformations of c-MET MD simulations with G756-2318

| c-MET | | | | | | | |
|---|---|---|---|---|---|---|---|
| **Residue** | **pH 7** | **pH 6** | **pH 5.5** | **Residue** | **pH 7** | **pH 6** | **pH 5.5** |
| B:61 | HIS | HIS | HIP | B:123 | ASP | ASH | ASH |
| B:148 | HIE | HIP | HIP | B:150 | HIE | HIE | HIP |
| B:159 | HIE | HIE | HIP | B:212 | HIS | HIP | HIP |
| B:267 | GLU | GLU | GLH | B:289 | HIS | HIE | HIE |
| B:394 | HIE | HIE | HIP | B:396 | HIS | HIE | HIS |
| B:484 | HIE | HIE | HIP | B:489 | GLU | GLH | GLH |
| B:542 | HIS | HIP | HIP | B:633 | HIE | HIE | HIP |
| B:644 | HIE | HIE | HIP | B:689 | HIS | HIS | HIP |

**Table S7.** pH-dependent residue protonation profiles employed in the initial MD setup of c-MET bound to 8016-2998.

| c-MET | | | | | | | |
|---|---|---|---|---|---|---|---|
| **Residue** | **pH 7** | **pH 6** | **pH 5.5** | **Residue** | **pH 7** | **pH 6** | **pH 5.5** |
| B:61 | HIS | HIS | HIP | B:123 | ASP | ASH | ASH |
| B:148 | HIE | HIP | HIP | B:150 | HIE | HIE | HIP |
| B:159 | HIE | HIE | HIP | B:212 | HIS | HIP | HIP |
| B:267 | GLU | GLU | GLH | B:289 | HIS | HIE | HIE |
| B:394 | HIE | HIE | HIP | B:396 | HIS | HIE | HIS |
| B:484 | HIE | HIE | HIP | B:489 | GLU | GLH | GLH |
| B:542 | HIS | HIP | HIP | B:633 | HIE | HIE | HIP |
| B:644 | HIE | HIE | HIP | B:689 | HIS | HIS | HIP |

**Table S8.** Initial protonation state configurations of titratable residues at multiple pH values for MD simulations of c-MET bound to Y020-8864.

| HGF | | | | | | | |
|---|---|---|---|---|---|---|---|
| **Residue** | **pH 7** | **pH 6** | **pH 5.5** | **Residue** | **pH 7** | **pH 6** | **pH 5.5** |
| A:158 | HIE | HIP | HIP | A:225 | HIE | HIP | HIP |
| A:257 | ASP | ASH | ASH | A:277 | HIS | HIP | HIP |
| A:335 | HIS | HIP | HIP | A:387 | HIE | HIP | HIP |
| A:423 | HIS | HIS | HIP | A:425 | HIS | HIP | HIP |
| A:551 | HIE | HIP | HIP | A:554 | HIS | HIP | HIP |
| A:633 | HIS | HIP | HIP | A:670 | GLU | GLU | GLH |
| A:682 | HIE | HIS | HIP | A:717 | HIS | HIE | HIE |

| c-MET | | | | | | | |
|---|---|---|---|---|---|---|---|
| **Residue** | **pH 7** | **pH 6** | **pH 5.5** | **Residue** | **pH 7** | **pH 6** | **pH 5.5** |
| B:61 | HIS | HIS | HIP | B:123 | ASP | ASH | ASH |
| B:148 | HIE | HIP | HIP | B:150 | HIE | HIE | HIP |
| B:159 | HIE | HIE | HIP | B:212 | HIS | HIP | HIP |
| B:289 | HIS | HIE | HIE | B:394 | HIE | HIE | HIP |
| B:396 | HIS | HIE | HIS | B:484 | HIE | HIE | HIP |
| B:489 | GLU | GLH | GLH | B:542 | HIS | HIP | HIP |
| B:633 | HIE | HIE | HIP | B:644 | HIE | HIE | HIP |
| B:689 | HIS | HIS | HIP | | | | |

**Table S9**. pH-driven protonation state assignments of key residues in the starting conformations of MD simulations for the c-MET-HGF complex in the presence of F142-0205.

| HGF | | | | | | | |
|---|---|---|---|---|---|---|---|
| **Residue** | **pH 7** | **pH 6** | **pH 5.5** | **Residue** | **pH 7** | **pH 6** | **pH 5.5** |
| A:158 | HIE | HIP | HIP | A:217 | GLU | GLU | GLH |
| A:225 | HIE | HIP | HIP | A:257 | ASP | ASH | ASH |
| A:277 | HIS | HIP | HIP | A:335 | HIS | HIP | HIP |
| A:387 | HIE | HIP | HIP | A:423 | HIS | HIS | HIP |
| A:425 | HIS | HIP | HIP | A:551 | HIE | HIP | HIP |
| A:554 | HIS | HIP | HIP | A:633 | HIS | HIP | HIP |
| A:670 | GLU | GLU | GLH | A:682 | HIE | HIS | HIP |
| A:717 | HIS | HIE | HIE | | | | |

| c-MET | | | | | | | |
|---|---|---|---|---|---|---|---|
| **Residue** | **pH 7** | **pH 6** | **pH 5.5** | **Residue** | **pH 7** | **pH 6** | **pH 5.5** |
| B:61 | HIS | HIS | HIP | B:123 | ASP | ASH | ASH |
| B:148 | HIE | HIP | HIP | B:150 | HIE | HIE | HIP |
| B:159 | HIE | HIE | HIP | B:212 | HIS | HIP | HIP |
| B:289 | HIS | HIE | HIE | B:394 | HIE | HIE | HIP |
| B:396 | HIS | HIE | HIS | B:484 | HIE | HIE | HIP |
| B:489 | GLU | GLH | GLH | B:542 | HIS | HIP | HIP |
| B:633 | HIE | HIE | HIP | B:644 | HIE | HIE | HIP |
| B:689 | HIS | HIS | HIP | | | | |

**Table S10.** Residue protonation state assignments reflecting pH effects in the initial MD models of c-MET bound to L083-0077.

| c-MET | | | | | | | |
|---|---|---|---|---|---|---|---|
| **Residue** | **pH 7** | **pH 6** | **pH 5.5** | **Residue** | **pH 7** | **pH 6** | **pH 5.5** |
| B:61 | HIS | HIS | HIP | B:123 | ASP | ASP | ASH |
| B:148 | HIE | HIP | HIP | B:150 | HIE | HIE | HIP |
| B:159 | HIE | HIE | HIP | B:212 | HIS | HIS | HIP |
| B:267 | GLU | GLU | GLH | B:289 | HIS | HIE | HIE |
| B:394 | HIE | HIE | HIP | B:396 | HIS | HIE | HIS |
| B:484 | HIE | HIE | HIP | B:489 | GLU | GLU | GLH |
| B:542 | HIS | HIS | HIP | B:633 | HIE | HIE | HIP |
| B:644 | HIE | HIE | HIP | B:689 | HIS | HIS | HIP |

**Table S11.** Residue-specific protonation state changes defining the pH-dependent initial MD structures of c-MET in complex with L083-1287.

| c-MET | | | | | | | |
|---|---|---|---|---|---|---|---|
| **Residue** | **pH 7** | **pH 6** | **pH 5.5** | **Residue** | **pH 7** | **pH 6** | **pH 5.5** |
| B:61 | HIS | HIS | HIP | B:123 | ASP | ASH | ASH |
| B:148 | HIE | HIP | HIP | B:150 | HIE | HIE | HIP |
| B:159 | HIE | HIE | HIP | B:212 | HIS | HIP | HIP |
| B:267 | GLU | GLU | GLH | B:289 | HIS | HIE | HIE |
| B:394 | HIE | HIE | HIP | B:396 | HIS | HIE | HIS |
| B:484 | HIE | HIE | HIP | B:489 | GLU | GLH | GLH |
| B:542 | HIS | HIP | HIP | B:633 | HIE | HIE | HIP |
| B:644 | HIE | HIE | HIP | B:689 | HIS | HIS | HIP |

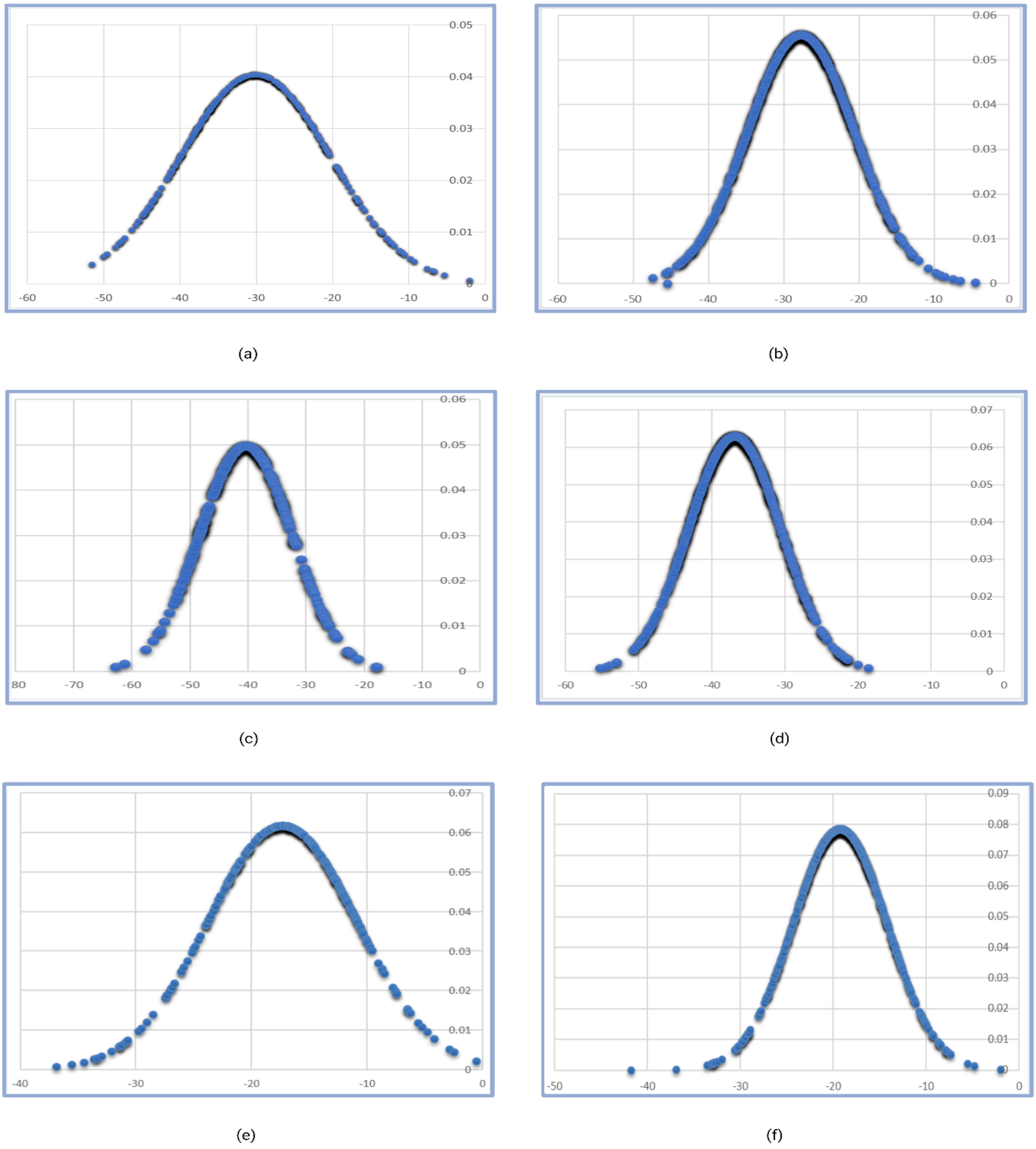


**Figure S1.** Z-scores calculation of virtual screening workflow results. (a) The graph show the normal distribution of docking results between HGF-MET complex and chemdiv library molecules (b) The graph show the Normal distribution of docking results between MET protein and chemdiv library molecules. (c) The graph show the the Normal distribution of docking results between HGF protein and chemdiv library molecules (d) The graph show the the normal distribution of docking results between HGF-MET complex and chemdiv library molecules (e) The graph show the normal distribution of docking results between MET protein and chemdiv library molecules. (f) The graph show the normal distribution of docking results between HGF protein and chemdiv library molecules.

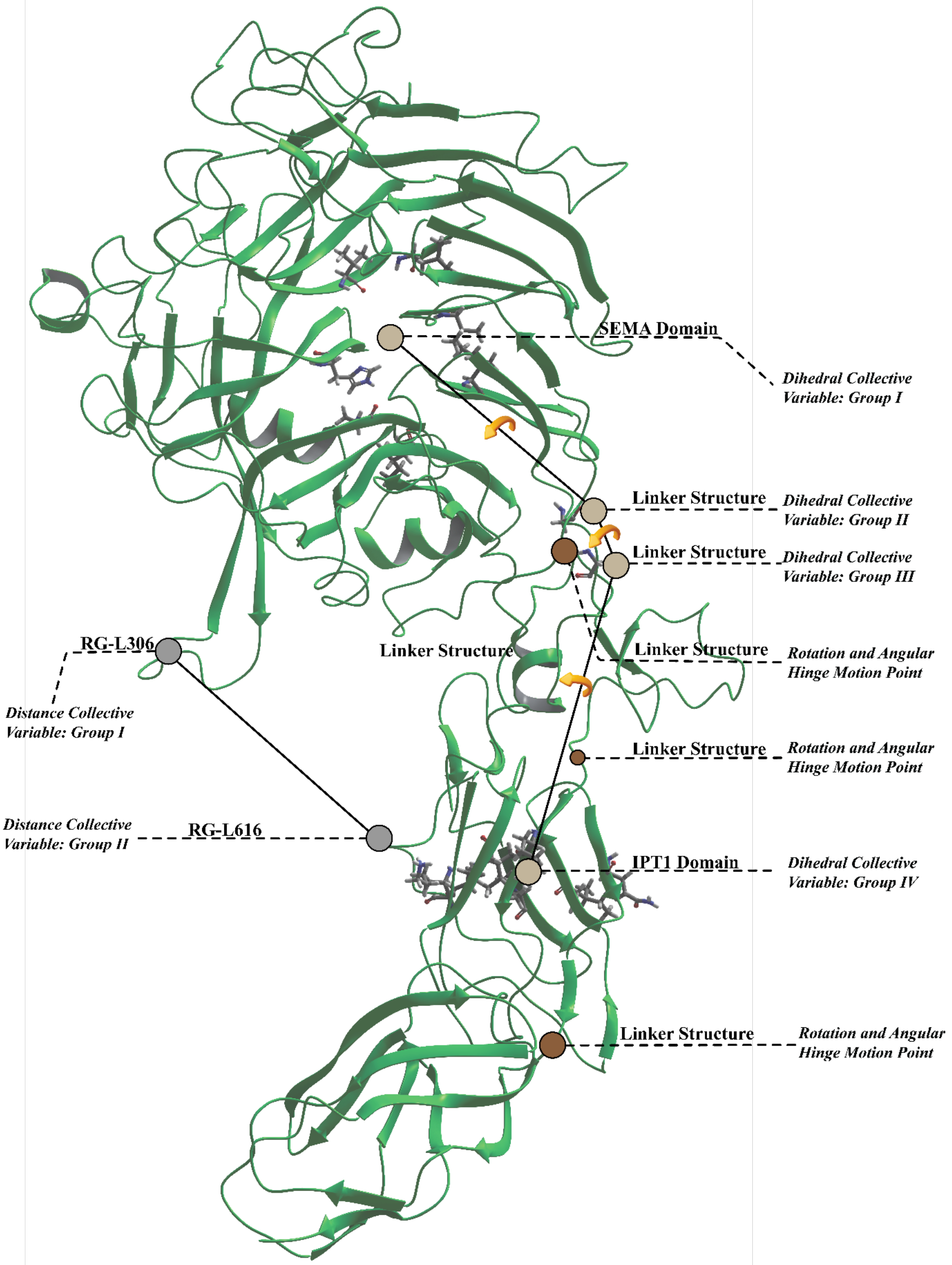


**Figure S2.** CV's of the c-MET: Distance CV's are colour coded with grey spheres, Dihedral CV's are defined with warm-beige spheres. Conformational drivers (motion points) are depicted by brown colour coded shperes. Structural information defined by using common literature identifications.

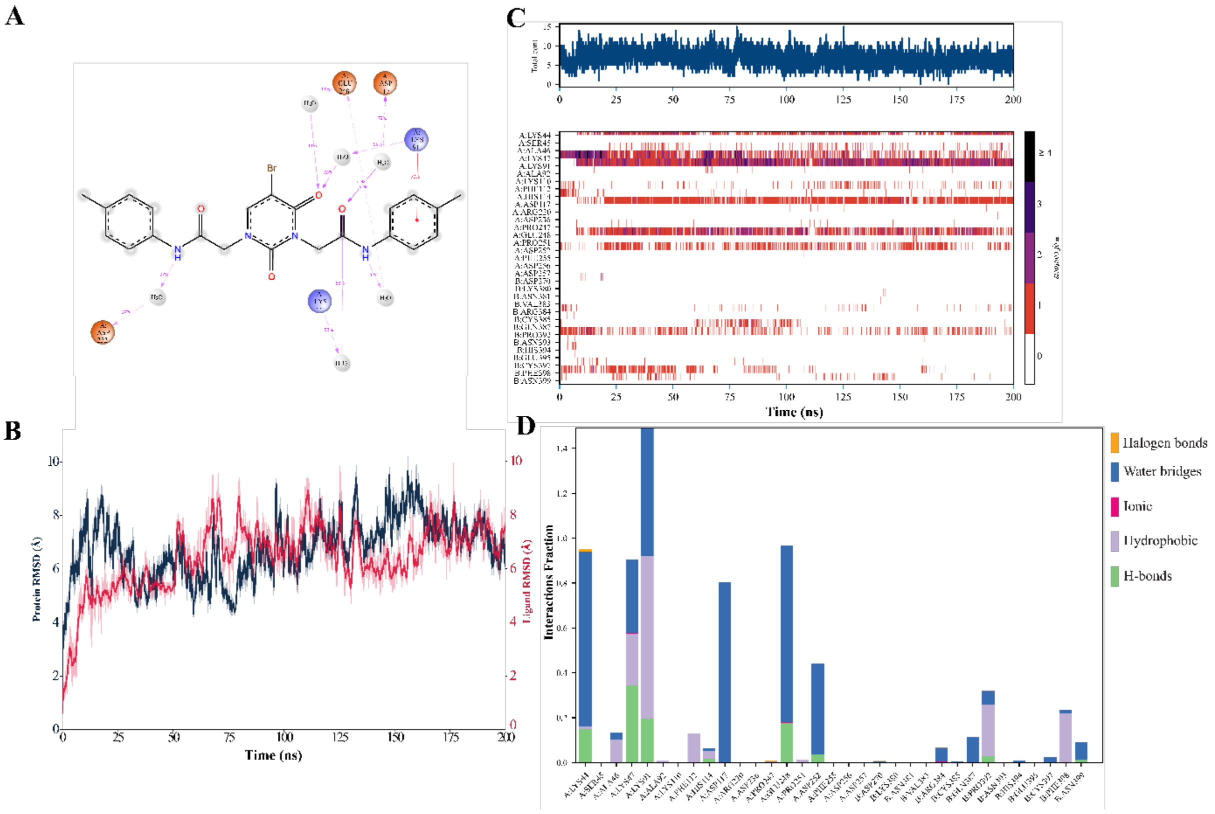


**Figure S3.** 200 ns MD simulation data of the lead compound 8008-3424. (A) Two-dimensional chemical structure of 8008-3424. Color-coded spheres denote residue categories and water molecules. (B) Time evolution of RMSD values for the protein (navy blue) and 8008-3424 (red) during the MD simulation. The y-axis reports RMSD (Å), while the x-axis denotes simulation time (ns). (C) Residue–ligand contact number map. The y-axis indicates residue indices, and the x-axis represents simulation time (ns). The color bar encodes the contact number range (0–4). The upper bar summarizes the total contacts accumulated over the simulation time. (D) Interaction fraction profile showing residue-wise contributions to ligand binding. The y-axis indicates interaction fraction, and the x-axis lists interacting residues. Interaction types are shown as: halogen bonds (orange), water bridges (dark blue), ionic interactions (pink), hydrophobic contacts (lilac), and hydrogen bonds (green).

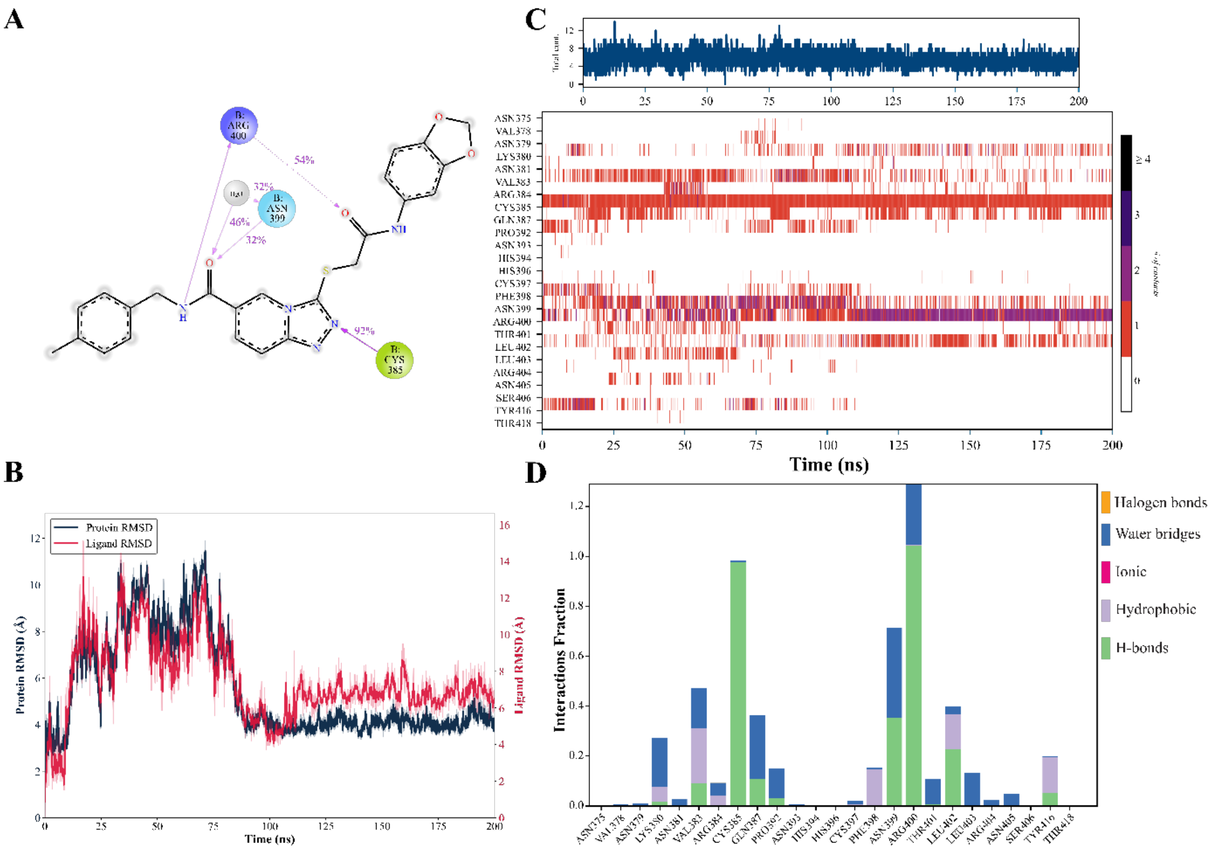

**Figure S4.** 200 ns MD simulation data of the Lead compound: L860-0593, A) 2D structure of L860-0593, Different colour coded spheres represent the residue type and water molecules. B) Protein (navy Blue) and L860-0593 (Red) RMSD values are depict y axis represent distance by Å unit. X axis represent time scale of the simulation by ns unit. C) Contact number map of the residue-ligand interaction, y axis reports the residue number and y axis corresponds time scale (ns) and also bar show contact number by using different colour respectively 0-4 range. Upper bar report total contact information during simulation time (ns). D) Fraction of the interaction plot show which residue contribute which contribution. Y axis illustrates fraction rate, x axis shows residue that interact with lead compound. Colors correspond by respectively Halogen bonds (orange), Water bridge (dark blue) , ionic (pink), hydrophobic (lilia), H bonds (green).

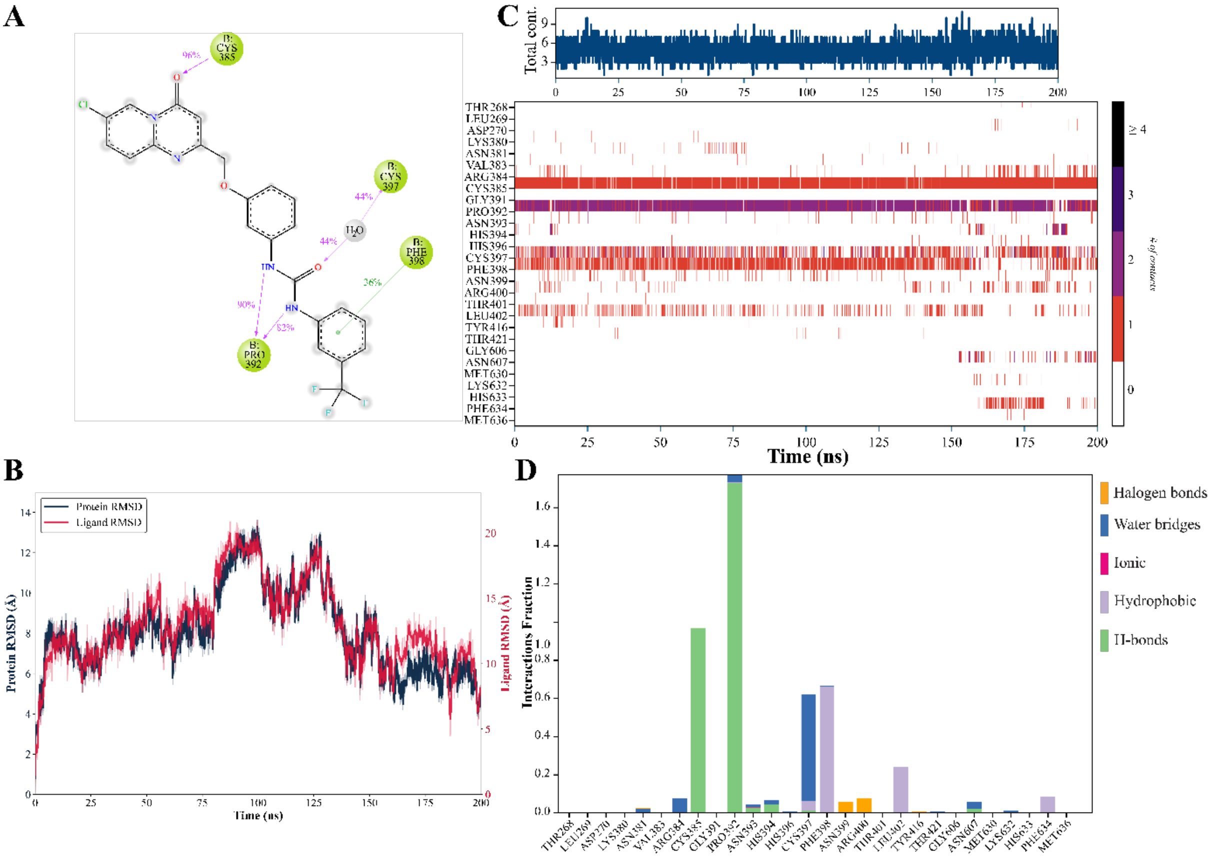

**Figure S5.** 200 ns MD simulation data of the lead compound G756-2318. (A) Two-dimensional chemical representation of G756-2318. Spheres with distinct colors indicate residue types and water molecules. (B) RMSD trajectories for the protein (navy blue) and G756-2318 (red) along the simulation. RMSD is reported in Å (y-axis) as a function of time in ns (x-axis). (C) Contact number map illustrating residue–ligand interactions across the trajectory. Residue numbers are shown on the y-axis, while simulation time (ns) is provided on the x-axis. Contact numbers are color-scaled within the 0–4 range. The top bar represents total contact frequency throughout the simulation. (D) Fractional contribution plot detailing residue-specific interaction persistence. The y-axis reports interaction fraction, and the x-axis lists interacting residues. Colors denote: halogen bonds (orange), water bridges (dark blue), ionic interactions (pink), hydrophobic interactions (lilac), and hydrogen bonds (green).

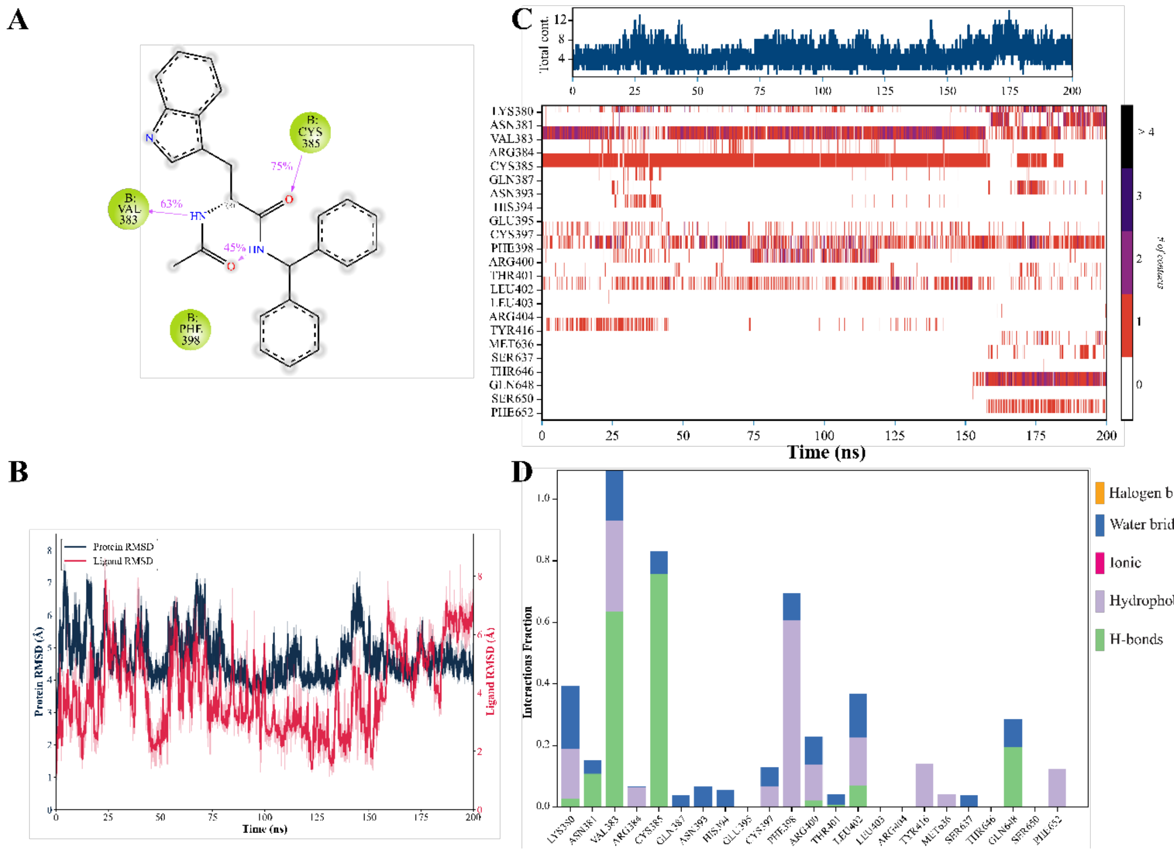


**Figure S6.** 200 ns MD simulation data of the lead compound 8016-2998. (A) Two-dimensional structure of 8016-2998. Residue classes and water molecules are represented by color-coded spheres. (B) RMSD profiles of the protein (navy blue) and 8016-2998 (red) over the simulation course. The y-axis indicates RMSD (Å), and the x-axis indicates simulation time (ns). (C) Residue–ligand contact number heatmap. The y-axis corresponds to residue indices, and the x-axis represents time (ns). The contact number scale is shown using a color legend (0–4). The upper bar summarizes total contact occurrence across the trajectory. (D) Interaction fraction analysis reporting the relative contribution of each residue to ligand binding. The y-axis indicates interaction fraction, and the x-axis lists residues involved in interactions. Interaction categories are colored as: halogen bonds (orange), water bridges (dark blue), ionic (pink), hydrophobic (lilac), and H-bonds (green).

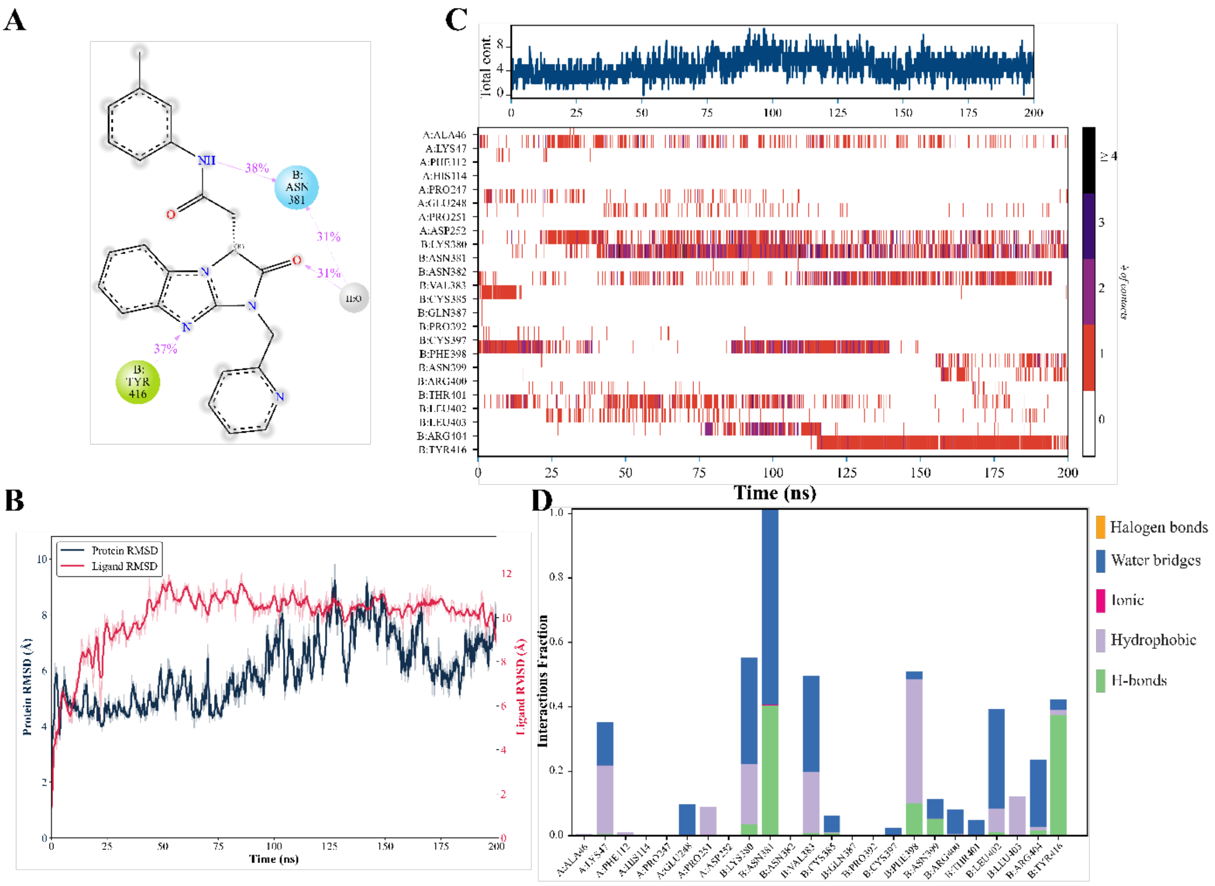


**Figure S7.** 200 ns MD simulation data of the lead compound Y020-8864. (A) Two-dimensional chemical structure of Y020-8864. Color-coded spheres represent residue types and water molecules. (B) RMSD time series of the protein (navy blue) and Y020-8864 (red) during the simulation. RMSD (Å) is plotted against time (ns). (C) Contact number map describing residue–ligand interactions along the MD trajectory. Residues are listed on the y-axis, and simulation time (ns) is shown on the x-axis. The contact numbers are encoded by a color gradient spanning 0–4. The upper bar provides the cumulative contact information over the entire simulation window. (D) Residue-level interaction fraction plot indicating the persistence and contribution of individual residues. The y-axis shows interaction fraction, while the x-axis lists interacting residues. Interaction types are encoded as: halogen bonds (orange), water bridges (dark blue), ionic interactions (pink), hydrophobic contacts (lilac), and hydrogen bonds (green).

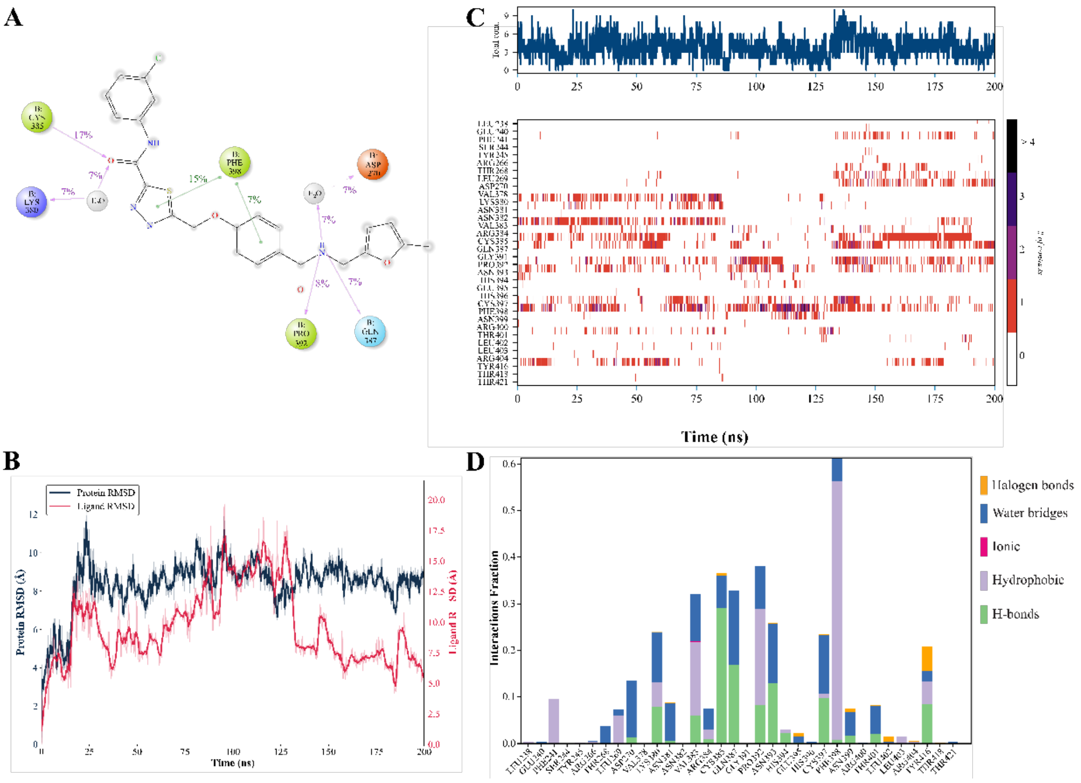


**Figure S8.** 200 ns MD simulation data of the lead compound L083-0522. (A) Two-dimensional depiction of L083-0522. Different residue categories and water molecules are indicated with color-coded spheres. (B) RMSD profiles for the protein (navy blue) and L083-0522 (red) across the simulation trajectory. The y-axis denotes RMSD (Å) and the x-axis denotes time (ns). (C) Contact number map highlighting residue–ligand contacts over time. The y-axis reports residue indices, and the x-axis represents simulation time (ns). Contact numbers are color-scaled (0–4). The upper bar summarizes total contact frequency during the simulation. (D) Interaction fraction analysis illustrating residue-specific contributions to ligand engagement. The y-axis represents interaction fraction and the x-axis lists interacting residues. Colors denote interaction types: halogen bonds (orange), water bridges (dark blue), ionic interactions (pink), hydrophobic interactions (lilac), and hydrogen bonds (green).

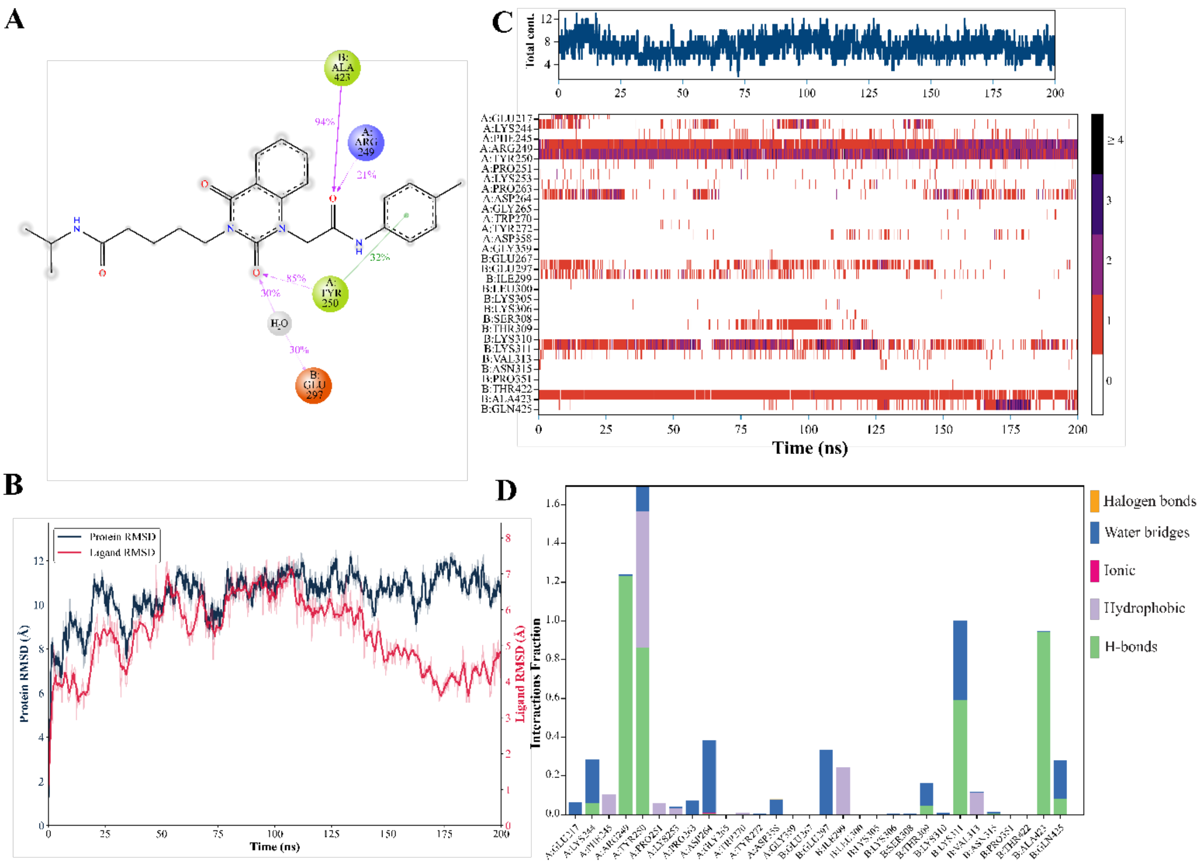



**Figure S9.** 200 ns MD simulation data of the lead compound F142-0205. (A) Two-dimensional chemical structure of F142-0205. Color-coded spheres correspond to residue classes and water molecules. (B) RMSD evolution of the protein (navy blue) and F142-0205 (red) along the MD simulation. RMSD values are shown in Å (y-axis) as a function of simulation time in ns (x-axis). (C) Residue–ligand contact number map. The y-axis indicates residue numbers, and the x-axis represents simulation time (ns). Contact counts are encoded with a color bar (0–4). The upper bar reports total contact accumulation throughout the simulation. (D) Interaction fraction plot showing the residue-wise contribution and persistence of interactions. The y-axis denotes interaction fraction, and the x-axis lists interacting residues. Colors represent: halogen bonds (orange), water bridges (dark blue), ionic interactions (pink), hydrophobic contacts (lilac), and hydrogen bonds (green).

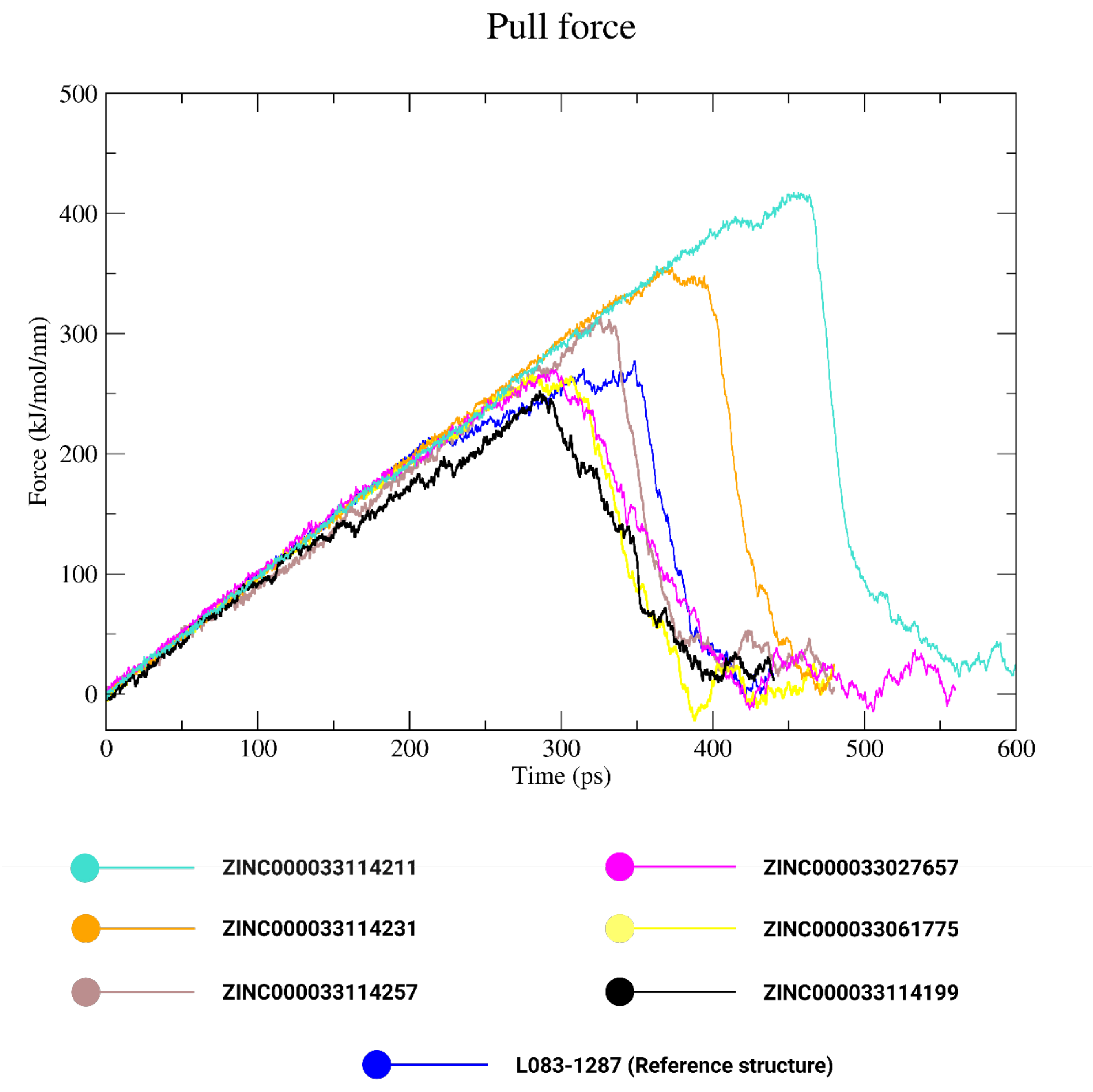


**Figure S10.** sMD simulation outcomes are presented for six compounds wtih L083-1287, which were selected based on the lowest binding scores obtained from the 50-ns MD simulation by targeting only c-MET. Each compound is colour coded with different colour. In the plot, the x-axis denotes simulation time (ps), whereas the y-axis reports the pulling force in units of $kJ \cdot mol^{-1} \cdot nm^{-1}$.

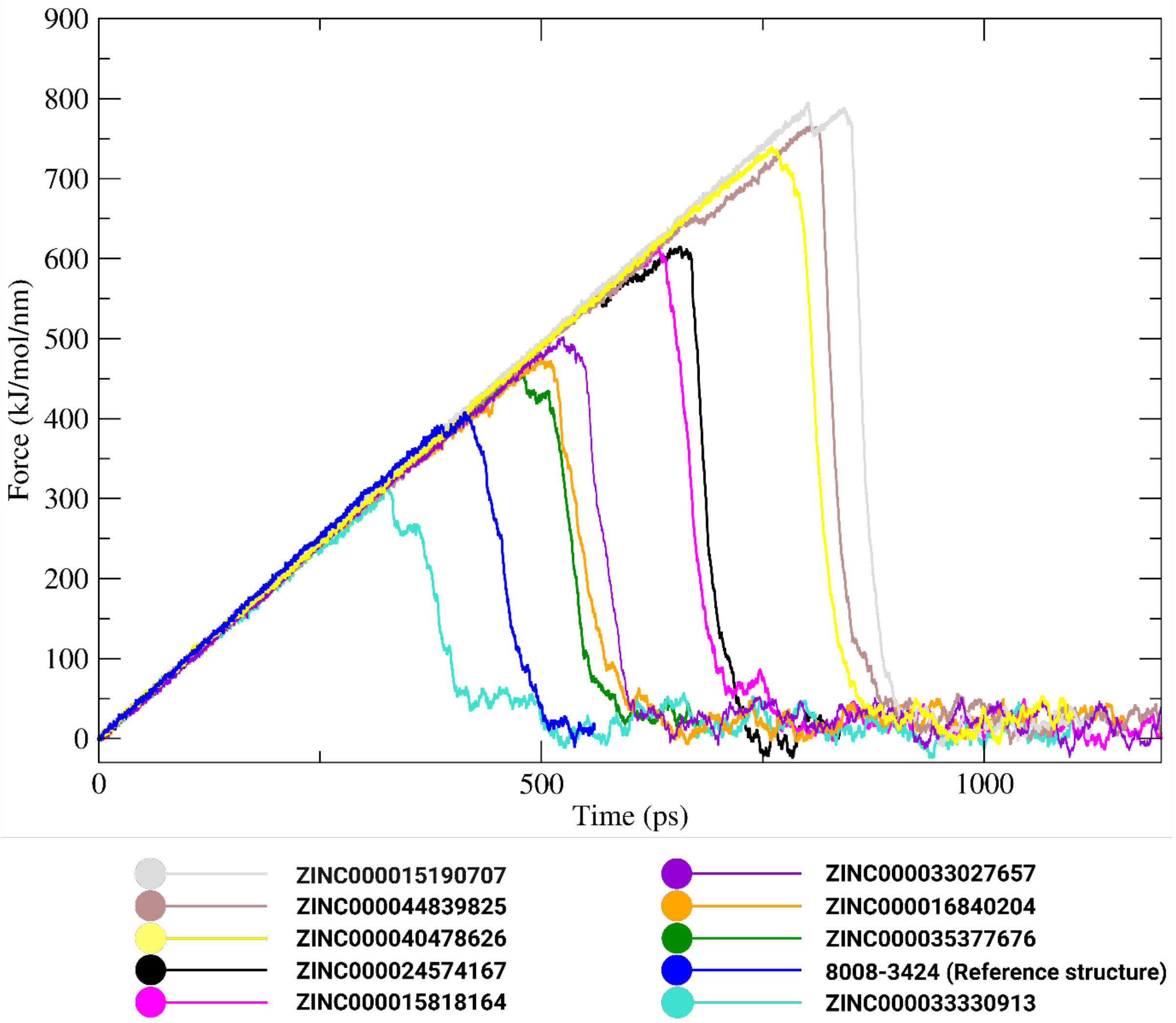


**Figure S11.** sMD simulation outcomes are presented for nine compounds with 8008-3424, which were selected based on the lowest binding scores obtained from the 50-ns MD simulation by targeting by c-MET/HGF complex. Each compound is colour coded with different colour. The x-axis depicts simulation time by unit of the ps, whereas the y-axis shows the pulling force in units of kJ·mol$^{-1}$·nm$^{-1}$.

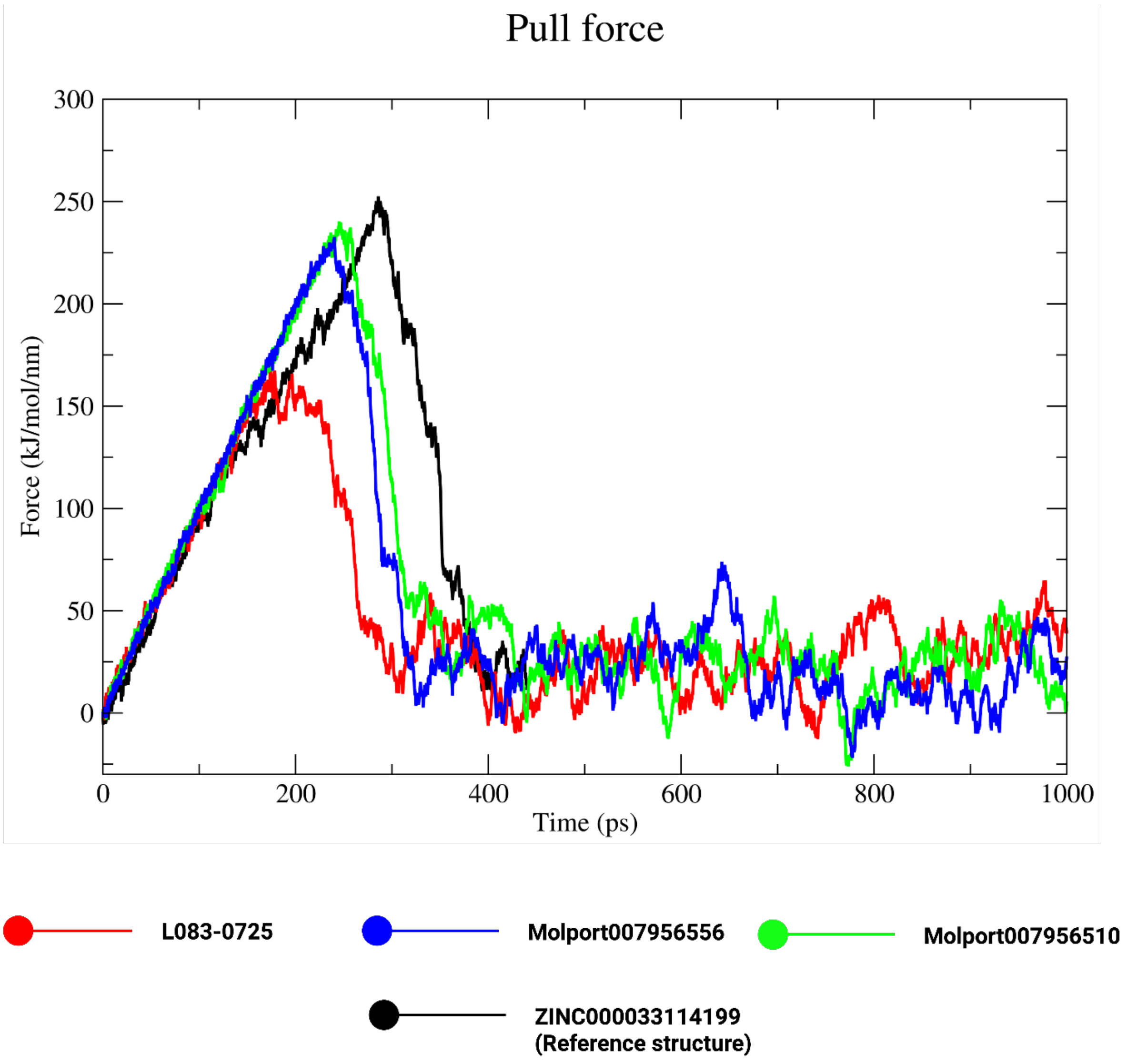


**Figure S12.** sMD simulation outcomes are presented for three compounds with L083-0077 (ZINC000033114199). Each compound is colour coded with different colour. The x-axis depicts simulation time by unit of the ps, whereas the y-axis shows the pulling force in units of $kJ \cdot mol^{-1} \cdot nm^{-1}$.

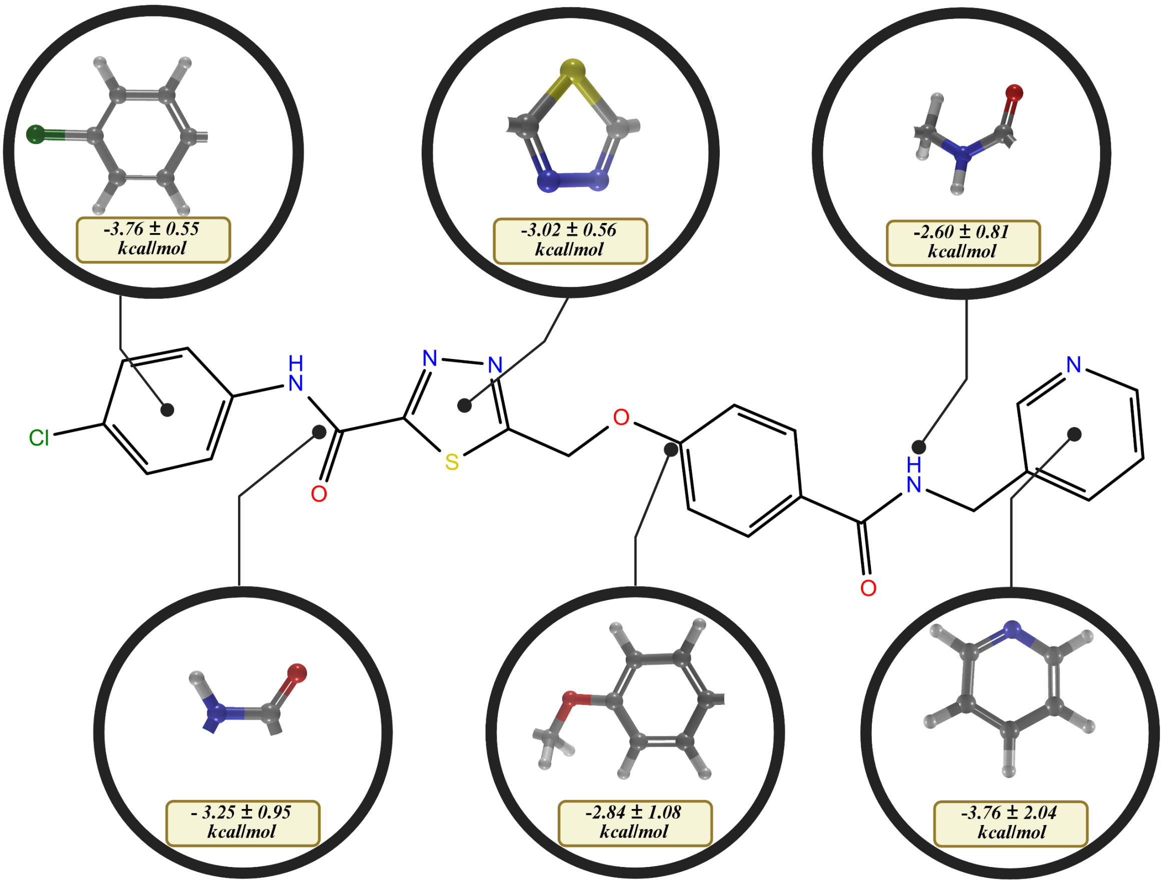


**Figure S13.** Fragment-based MM/GBSA binding free-energy contributions of lead compound L083-0077, obtained from 200 ns molecular dynamics simulations (3 replicas).

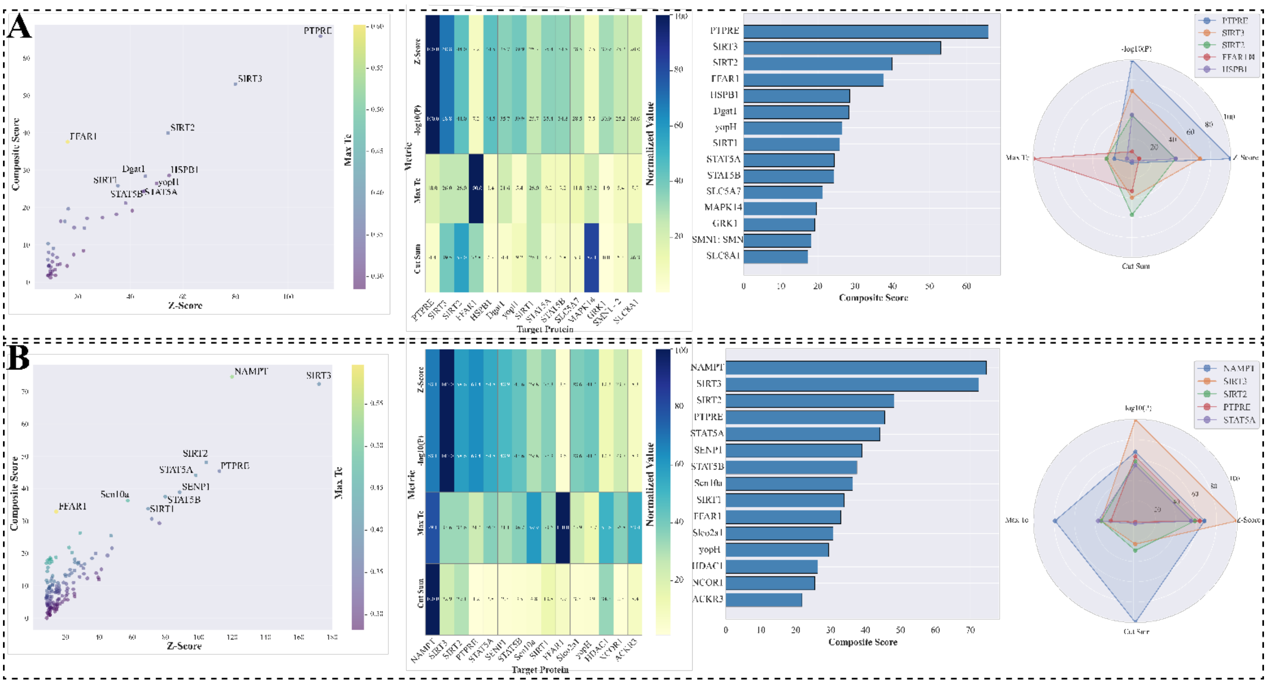


**Figure S14. Examining the off-target potential of the lead compounds.** From left to right, the first panel shows the overall off-target binding landscape as a scatter plot, where the x-axis represents the SEA Z-score and the y-axis represents the composite score, with the color scale indicating Max Tc (maximum Tanimoto coefficient). The second panel presents a heatmap of the normalized SEA metrics for the top fifteen predicted off-target proteins (x-axis: target/off-target proteins; y-axis: SEA metrics). The third panel displays the ranked composite scores for the top fifteen predicted targets (x-axis: composite score; y-axis: target/off-target proteins). The fourth panel summarizes the main SEA contributors as a radar plot, where each axis corresponds to a specific metric (log10(E-value), Z-score, Max Tc, and composite score) and the radial distance indicates the relative magnitude of each contributor. (A) Off-target profile of L083-0077. (B) Off-target profile of L083-1287.

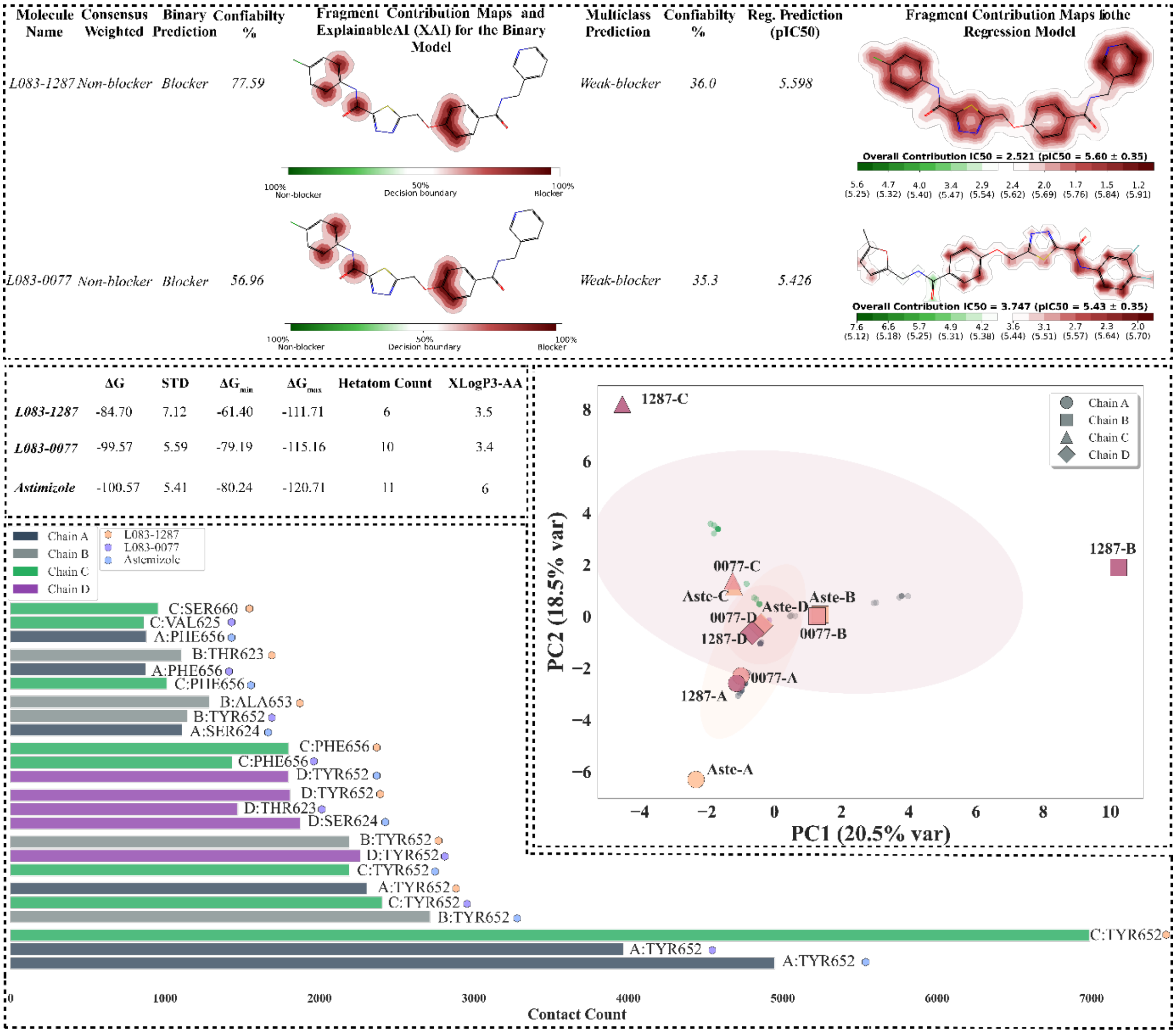


**Figure S15**. **Comparative structural and dynamic analysis of hERG-ligand complexes.** Electrostatic surface maps (top panels) illustrate the binding orientation and charge distribution differences among Astemizole, L083-0077, and L083-1287 within the hERG cavity. The bar plot (bottom left) shows the residue-specific contact frequencies across the four channel subunits (A-D), highlighting TYR652, PHE656, and SER624 as the most recurrent interaction sites. The PCA scatter plot (bottom right) represents the principal components of interaction dynamics, where Astemizole and L083-0077 cluster closely, indicating similar binding behavior, while L083-1287 exhibits distinct positioning on subunits B and C. Overall, the figure demonstrates that Astemizole establishes the highest total contact count (≈ 11 × $10^3$) and strongest π-cation/π-π interactions with TYR652 and PHE656, whereas both lead compounds maintain moderate contact patterns with reduced aromatic stacking contributions.

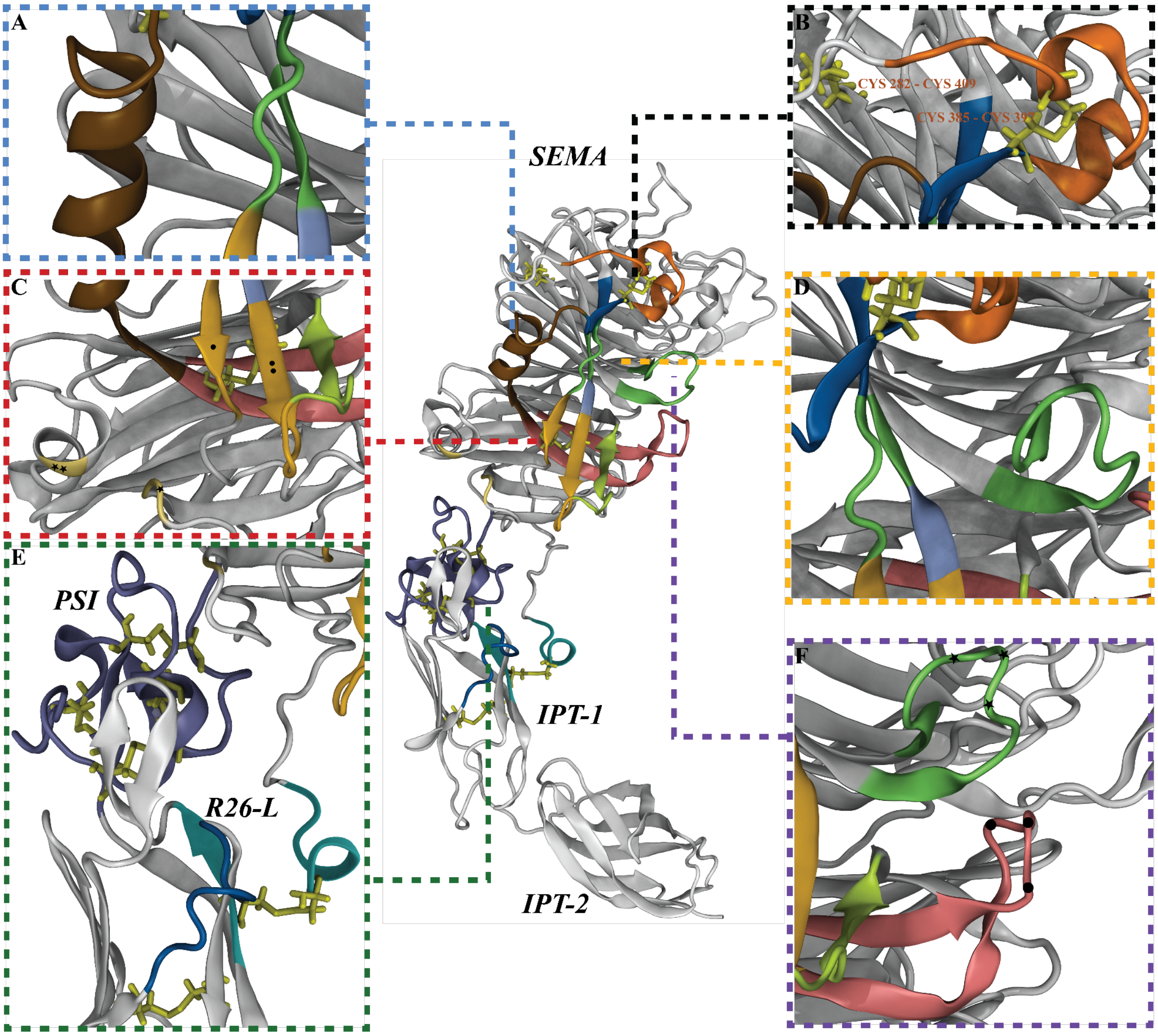


**Figure S16: Structural definition of the c-MET for mechanistic elucidation A)** The brown-colored helix mediates motion transfer between Interface I and Interface III (Helix 367-377). The auxiliary regions supporting this motion transfer include the segments preceding the helix (365-367) and following the helix (378-383), which are also shown in brown in the main figure. **B)** The region coded in metallic orange represents Interface I (386-402). In addition, the regions shown in matte navy blue the cysteine-containing β-sheet (382-385) and the β-sheet extension (418-420) function as motion-transfer elements in the sequential angular hinged-rotation mechanism. These segments also contain parts of both Interface I and Interface II, acting as shared structural junctions. The stabilizing cystines displayed in stick representation on the β-sheet involved in the Interface I-Interface II interaction are: CYS385-CYS397 disulfide bond, CYS298-CYS409 disulfide bond **C)** Interface III and auxiliary groups contributing to motion transfer; C1) When Interface III is evaluated, it can be subdivided into three distinct Interface III elements: (I) Matte gold β-sheets with different dot patterns: the single-dot β-sheet (β425-428) and the twice-dot long β-sheet, which constitute the main mechanistic regions. The ice-blue region corresponds to the shared interaction surface between Interface II and Interface III, serving as a key motion-transfer junction. (II) The pistachio-green loop/β-sheet (RG-L306) is not classified as part of the interface in the literature; however, it represents a highly critical loop for mechanistic motion transfer. (III) Metallic gold regions with different star patterns (reported in the literature as residues critical for c-MET ligand affinity): the single-star region (ARG469) and the double-star region (GLN332). This structural element is important for the solvent shield, though its mechanistic contribution is presumed to be relatively minor. C2) (I) The regulatory loop and β-sheet (346-361) are shown in matte pink. (II) The Interface I-III motion transfer region is highlighted in brown. C3) The stabilizing and guiding disulfide architecture formed by CYS298-CYS363 is illustrated in stick representation and color-coded in yellow. **D)** (I) The regions shown in green correspond to the primary interaction regions of Interface II. (II) The matte navy-blue regions represent intersection zones with Interface I (382-385 and 418-420). (III) The blue-violet region denotes a shared overlap region with Interface II (418-420). **E)** Angular hinge motion and rotation regulatory groups: (I) The PSI domain, which provides mechanical stabilizer capability, contains four yellow-coded cystine pairs: CYS526-CYS561, CYS529-CYS545, CYS520-CYS538. This domain is coded in blue-violet. (II) The regulatory sheet/helix26-loop (R26-L) structure, which serves to reduce the number of degrees of freedom, contains the CYS26-CYS584 cystine pair (shown in yellow stick representation) and is coded in pastel cyan. (III) The Regulatory Loop (RG-L) 614-618 is shown in navy blue, and the adjacent RG-L stabilizer cystine pair (CYS610-CYS624) is coded in yellow. **F)** Solvent Dependent Regulatory Loops (SDRG-L) 265-276 & 346-361 (I) The SDRG-L270 (residues 265-276), which is part of Interface II, is coded in lime green. (II) The SDRG-L352 (residues 346-361) is coded in matte pink.

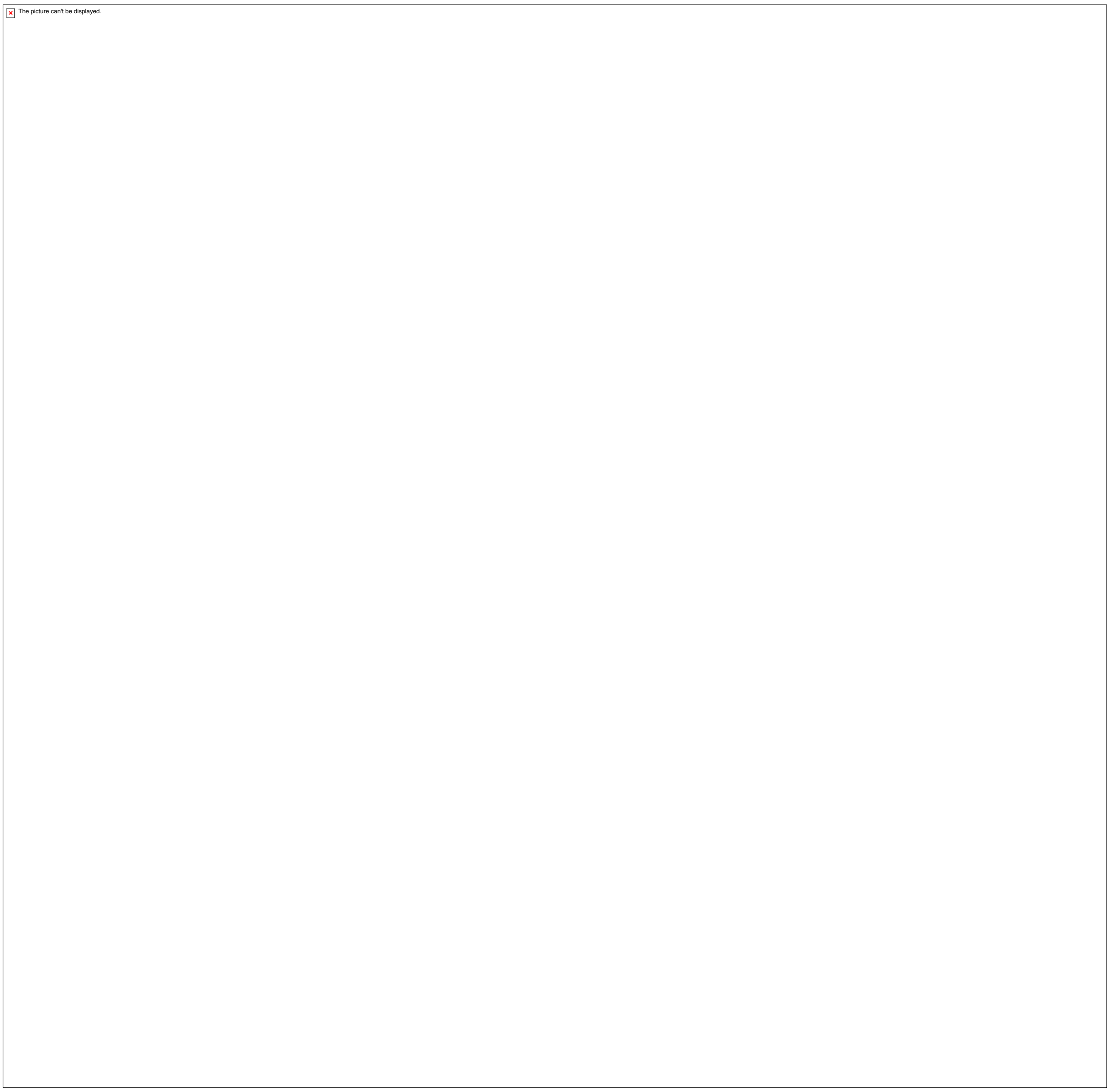


**Figure S17.** Mechanism controller Cyctine groups of the c-MET: Each domain is defined on the illustration and cyctine groups are report by their essential tasks. Cyctine groups are colour coded with matt yellow colour.

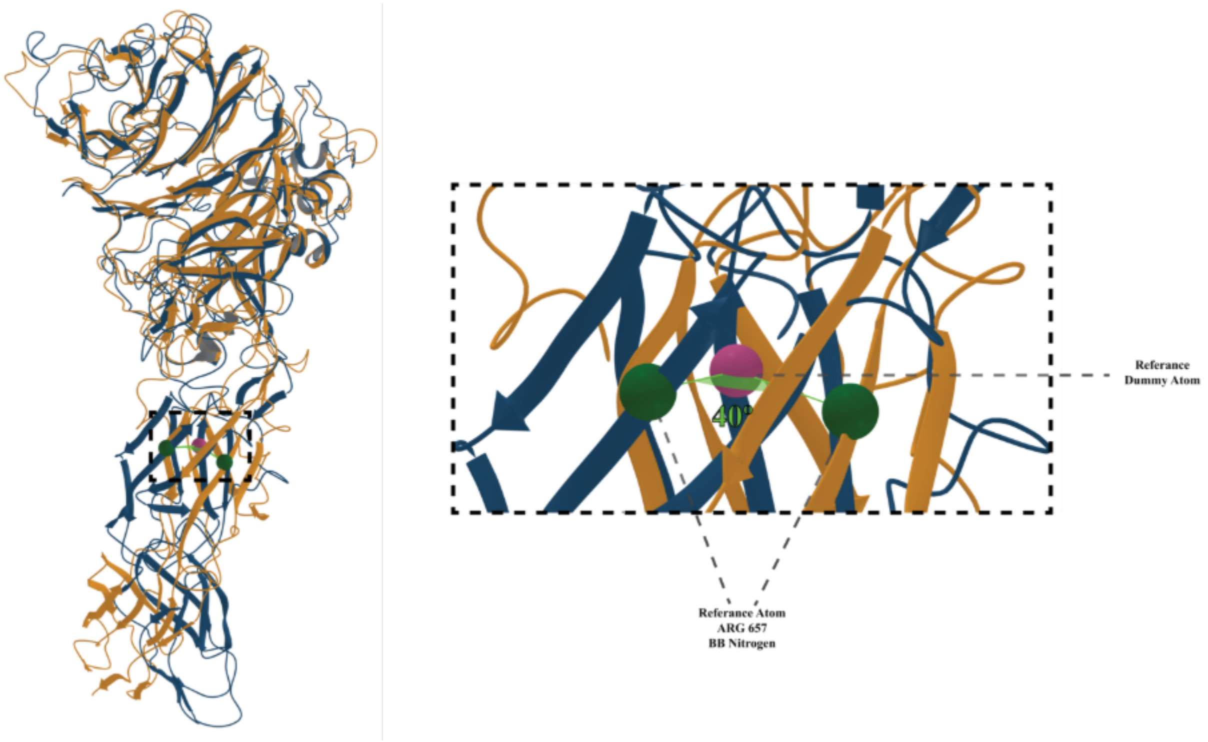


**Figure S18.** Total rotation derived from a representative simulation exhibiting the average clockwise rotational behavior. The analysis was performed by defining initial and final positions based on the IPT1 domain.

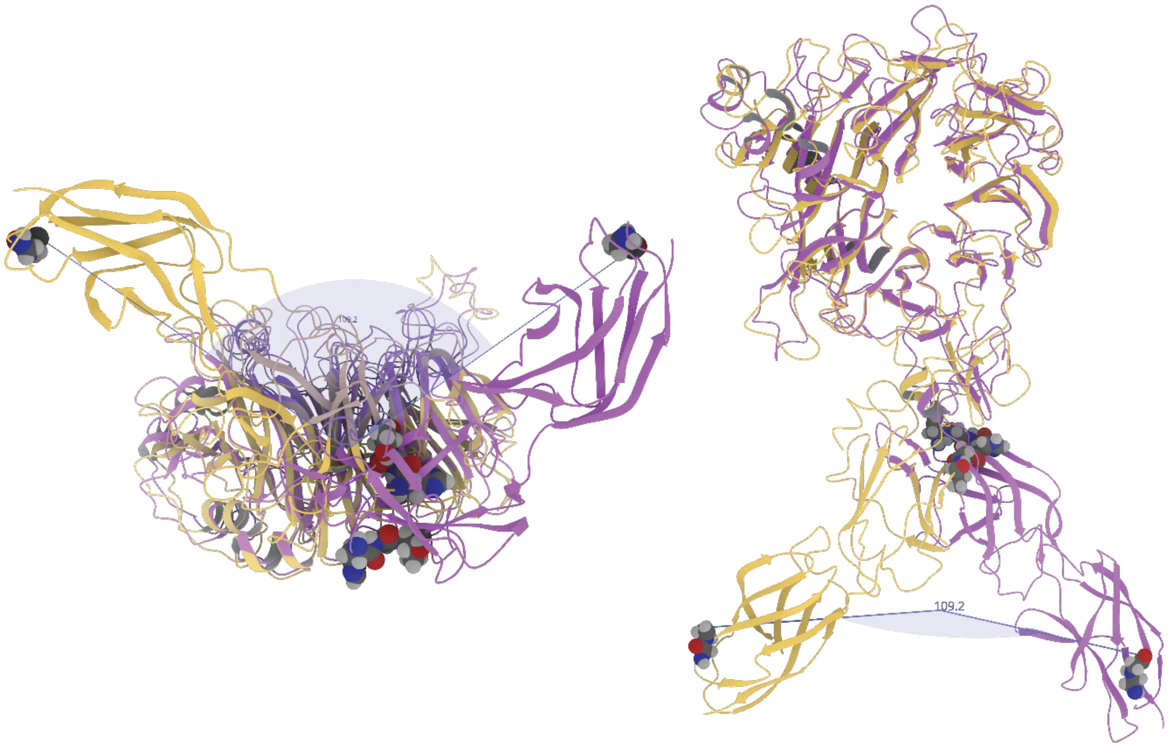


**Figure S19: Structural overlay of the 109.2° angular hinge rotation captured in metadynamics.** Superposition of conformations obtained from two independent metadynamics simulations initiated from the energy-minimized 7MO7 structure. The overlay highlights a characteristic 109.2° angular hinge rotation between the initial and rotated states (yellow and purple, respectively). The right panel shows the same overlay after an additional ~90° rotation relative to the left panel, providing an orthogonal view of the domain-scale rearrangement.

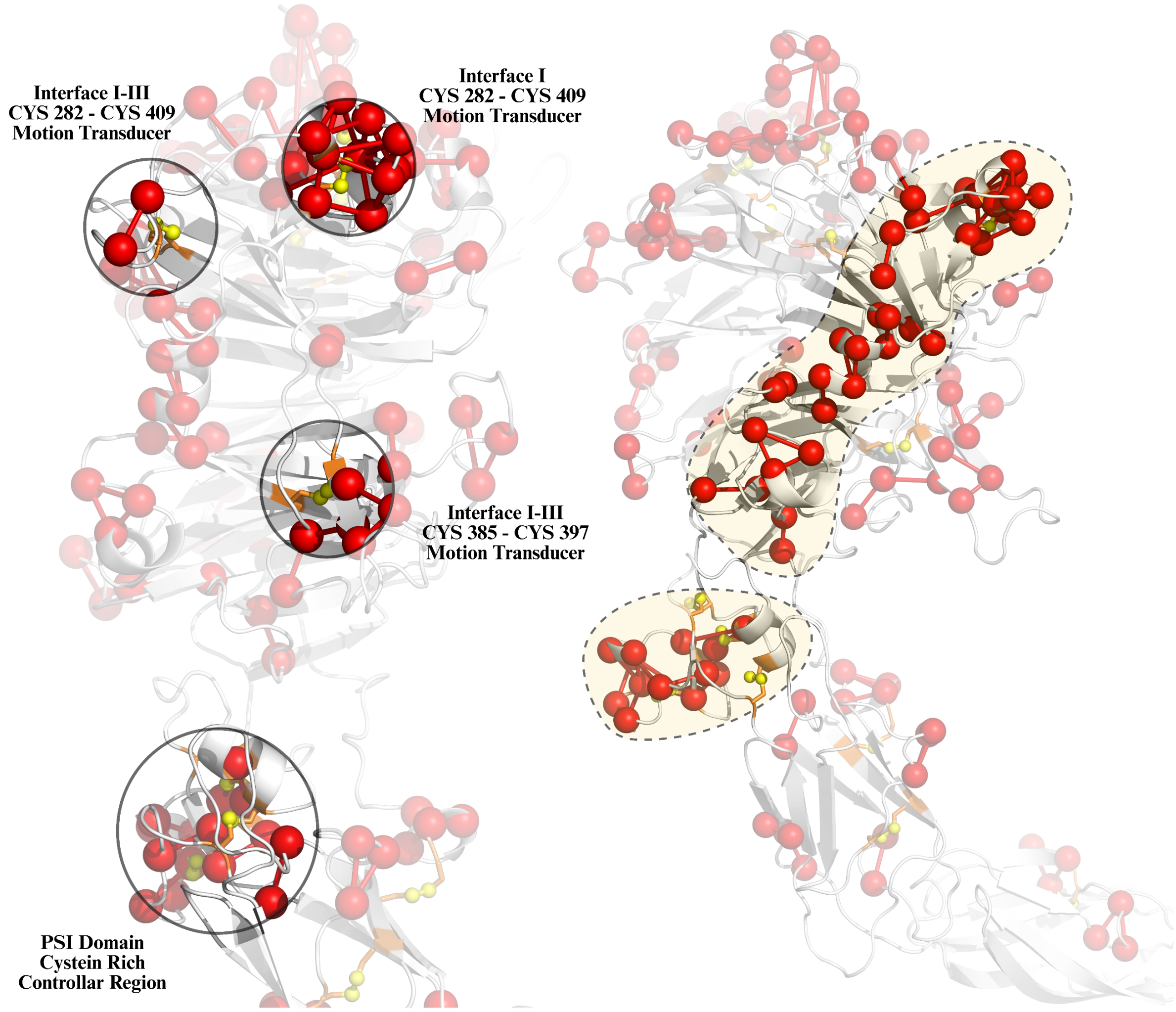


**Figure S20.** Structural organization of cystine groups, pivot regions, and highly frustrated interaction patterns. (A) Cystine groups involved in the identified interaction pathways are highlighted, illustrating their spatial arrangement and potential role in motion transmission. (B) Distribution of two pivot points together with other highly frustrated regions across the structure. Interaction pathways form distinct patterns converging toward the pivot centers, including those extending into the PSI domain.

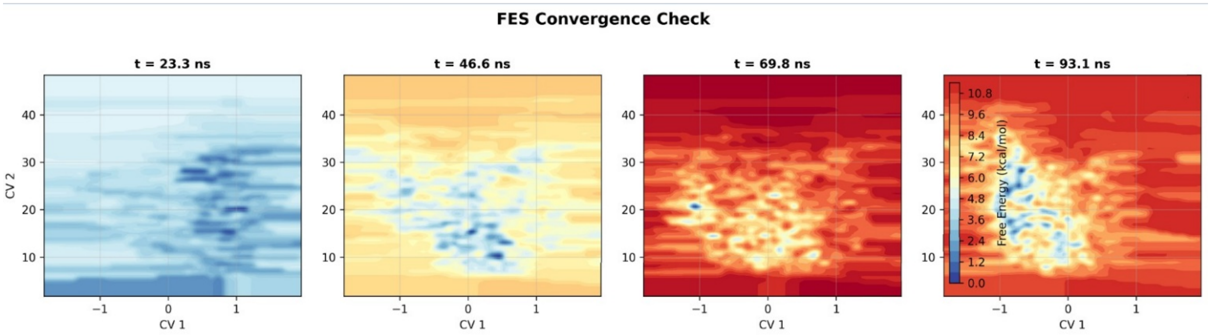


**Figure S21: FES convergence across simulation quarters.** Time-resolved (FES) reconstructed from the metadynamics trajectory split into four consecutive quarters (t = 23.3, 46.6, 69.8 and 93.1 ns), enabling assessment of progressive stabilization and convergence of the sampled landscape.

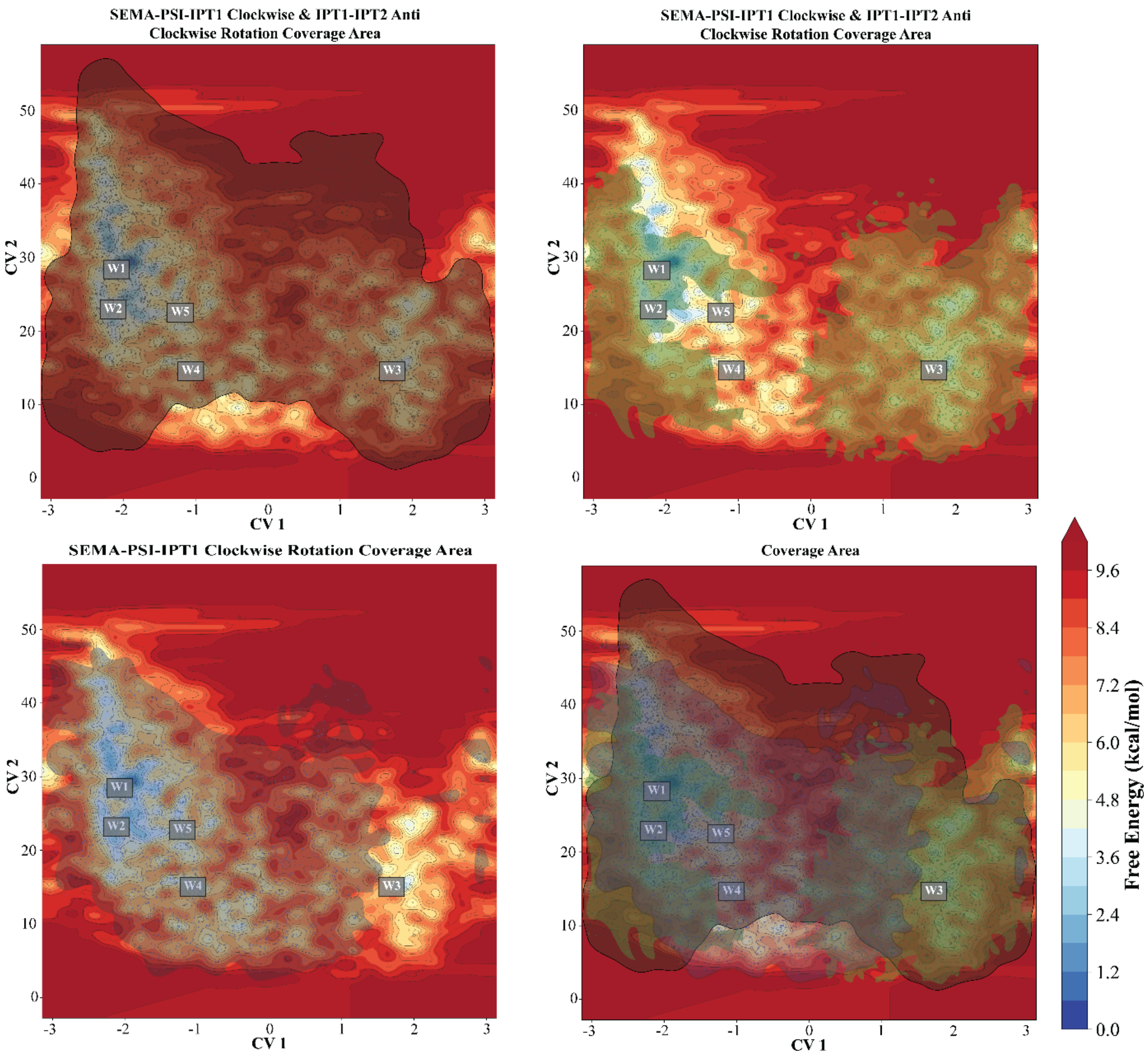


**Figure S22.** Three different Metadynamics simulation replicas for Apo form of the c-MET. All Well definations show with numbers by W1 to W5. In all FES, CV1 corresponds the radial coordinate that is coming from the dihedral angels. CV2 corresponds the distance (Å). The color bar at the bottom right indicates the free energy scale and applies to all plots. A) It include conformations that carry SEMA-PSI-IPT1 Clockwise rotation and IPT1-IPT2 anticlockwise rotation. Its coverage area shows with transparent black overlay. B) SEMA-PSI-IPT1 Clockwise rotation and IPT1-IPT2 clockwise rotation data include conformation that define the clockwise rotations by utilized transparent green overlay. C) The FES elucidat only SEMA-PSI-IPT1 clockwise rotation Dynamics and for illustration of the coverage are of the data, Dark blue overlay is used. D) The FES data and overlayers shows whole conformations of the data for clear seperation of the essential Dynamics types.

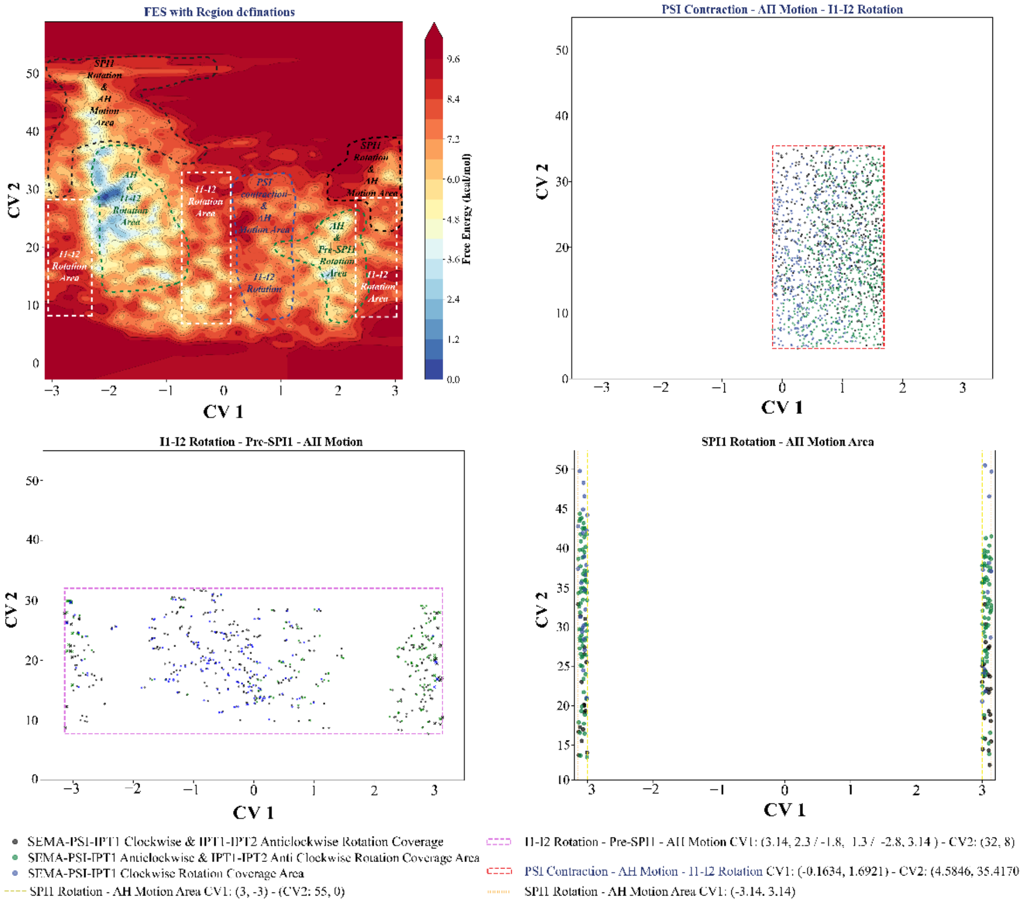


**Figure S23.** The figure aims explanation of the indicates of the region that shows essential dynamics. CV1 include distance data by Å unit and CV2 includes of the radial coordinates (Å). Black dots show SEMA-PSI-IPT1 clockwise rotation and IPT1-IPT2 anticlockwise rotation structures. Green dots show SEMA-PSI-IPT1 clockwise rotation and IPT1-IPT2 clockwise rotation. Dark blue dots show SEMA-PSI-IPT1 Clockwise rotation. A) To define the essential dynamics, the FES was partitioned into discrete dynamical regions by grouping free-energy wells, transition barriers, and metastable basins. B) Plot include different Metadynamics simulation structures by using special colour coded dots. Red dash box rectangle encompass PSI contraction-stabilization frames mostly. Also because of the complex structure of the protein IPT1-IPT2 (I1-I2) rotation and as a stabilization contributor angular hinge (AH) motion dynamics are included. C) The illustration provide the seperation of the dominant I1-I2 rotations (see Panel A) by pink dash box area. D) SEMA-PSI-IPT1 (SPI1) rotation is show between CV1 3.14/3 : -3.14/-3 and CV2 30 Å – 50 Å by seperation yellow and orange dash box.

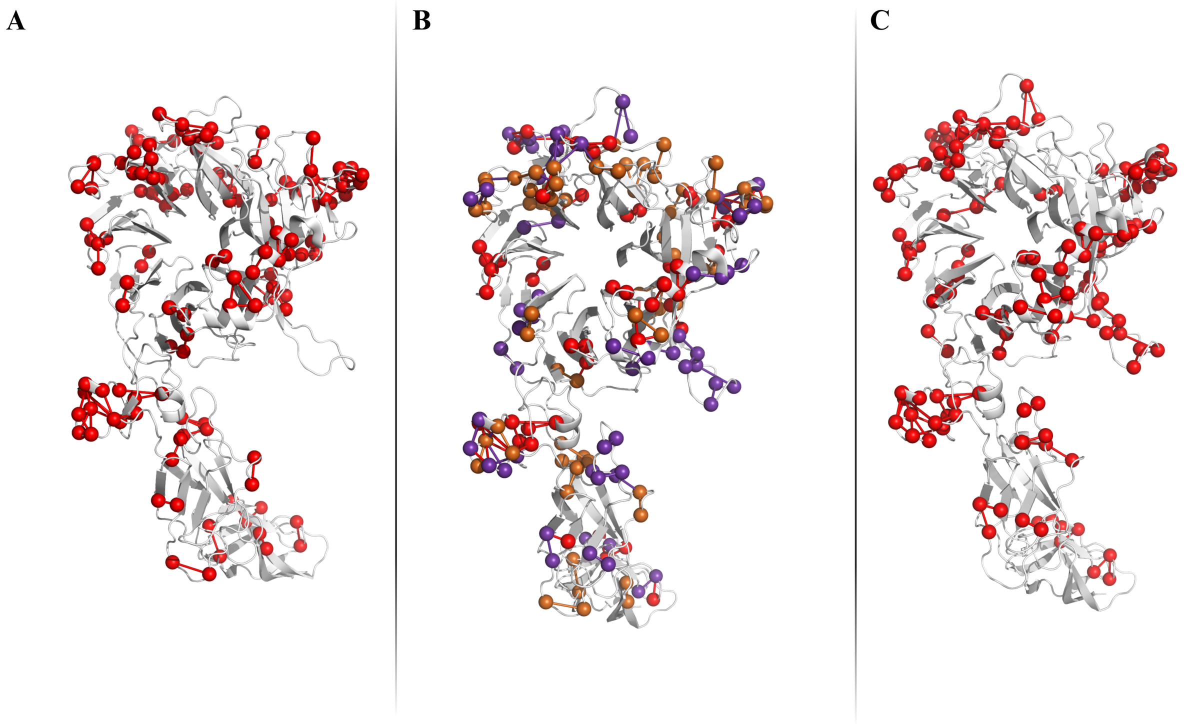


**Figure S22.** Frustration profiles derived from representative metadynamics simulations. **A)** Frustration profile of the holo conformation. Red pairs indicate the top 100 most frequently observed residue–residue interactions classified as highly frustrated along the trajectory. **B)** Spatial distribution of the top 100 most frequently observed highly frustrated pairs mapped onto the same conformation for both holo and apo metadynamics simulations. Shared highly frustrated regions are shown in red, while regions specific to the holo and apo forms are highlighted in gold and purple, respectively. **C)** Top 100 most frequently observed highly frustrated residue pairs along the trajectory of the apo-initiated simulation.

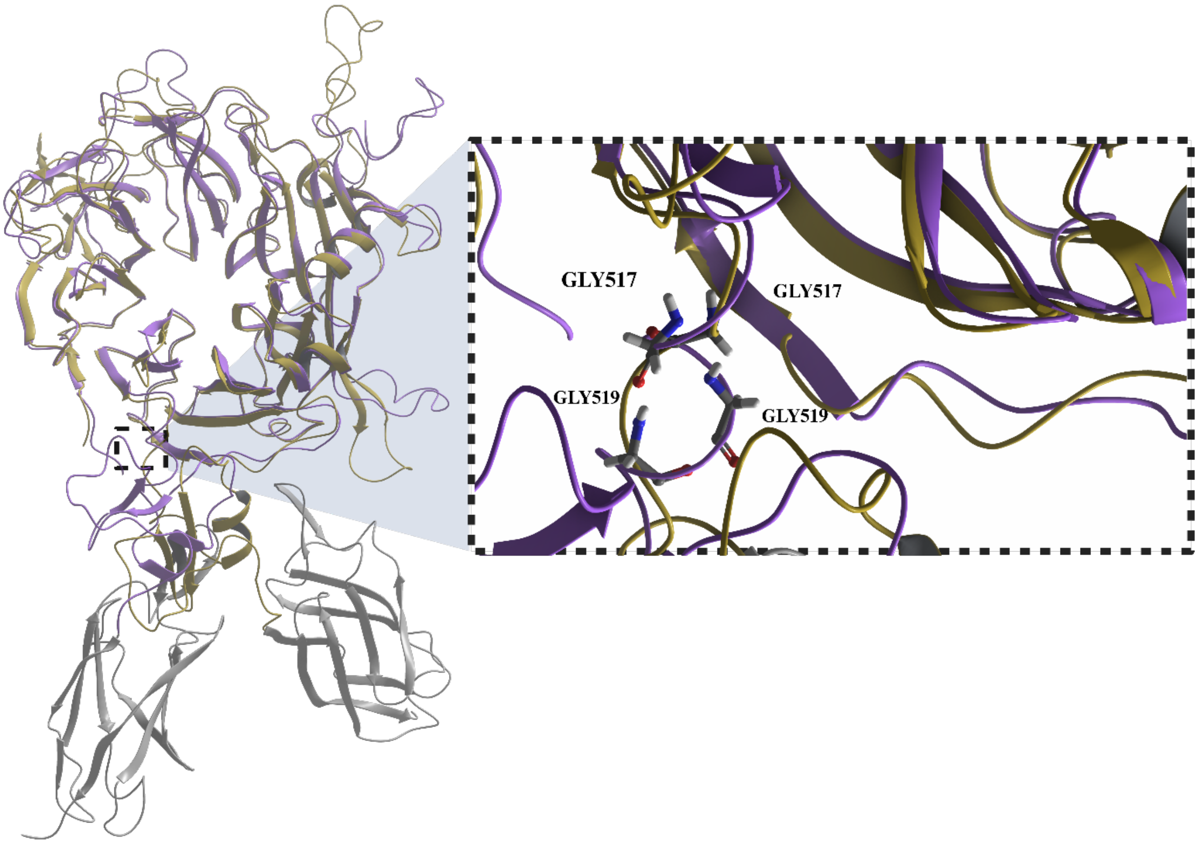


**Figure S23.** Hypothesis of the c-MET activation and inactivation mechanism that is coming from the experimental literature information: Active Conformation (PDB:2UZX) is illustrated with orange colour. Inactivation liked conformation is showed by purple (PDB: 1SYH).
